# A non-enzymatic role for METTL3 as an Androgen Receptor co-regulator that promotes prostate cancer proliferation

**DOI:** 10.64898/2026.07.08.737095

**Authors:** Raymond J. Kostlan, John T. Phoenix, Audris Budreika, Carli D. Deegan, Marina G. Ferrari, Elise T. Warren, Pushpinder S. Bawa, Charles S. Rogers, Divya Dureja, Muzna Ali, Govinda R. Hancock, Kristen S. Young, Gopal Gupta, Abhishek Solanki, Donald J. Vander Griend, Sean W. Fanning, Steven Kregel

**Author notes:** Corresponding Author: Steven Kregel, Department of Cancer Biology, Loyola University Chicago, Stritch School of Medicine Health Sciences Division, Cardinal Bernardin Cancer Center, 2160 South First Avenue Building 112, Room 205, Maywood, IL 60153.

## Abstract

Metastatic prostate cancer (PCa) continues to be a major cause of death in males, despite advances in treatment. Most treatment focuses on targeting the Androgen Receptor (AR), the main oncogene responsible for driving most prostate tumors. Despite these therapies targeting AR, the majority of patients still succumb to AR-driven disease. Therefore, there is a critical need for understanding how AR functions to promote prostate cancer growth and identify alternative therapeutic targets in AR-driven PCa. One avenue garnering attention is targeting epigenetic regulators that promote AR-activity; however, the importance of epitranscriptomic regulators, like those that modify mRNAs, is not well understood. Here, we identify a new role for the key catalytic subunit of the RNA N6-methyladenosine (m^6^A) transferase complex, METTL3, as an AR-coregulator. METTL3 is overexpressed in prostate tumors compared to normal tissue, and METTL3 protein is elevated in AR-expressing cell lines. Depletion of METTL3 significantly reduces proliferation of cancer cells and has no effect on the growth of non-transformed prostate epithelial cells, despite decreasing global m^6^A levels on mRNA. The catalytic activity of METTL3 is dispensable for the growth of both non-transformed and PCa cell lines, as pharmacologic inhibition of METTL3 does not inhibit proliferation, despite the reduction of global m^6^A on mRNA. Overexpression of both wild-type and catalytically inactive METTL3 mutants enhances cell viability and rescues cells in which METTL3 is knocked down. Finally, we report on direct interaction between AR and METTL3, their co-localization on chromatin, and reduced AR-cistromic occupancy within cells with METTL3 knockdown. Together, these findings identify a non-enzymatic role for METTL3 in supporting AR-driven transcriptional programs and PCa proliferation.

## Introduction

Globally, prostate cancer (PCa) mortality ranks second only to lung cancer in men, with nearly 400,000 deaths reported worldwide (1). If diagnosed early, patients with nonmetastatic prostate cancer exhibit a high survival rate; however, if distant metastasis occurs, the 5-year survival rate is reduced to 37% (2). Surgical or chemical castration targeting the androgen receptor (AR) signaling axis has been the mainstay of prostate cancer treatment (3), but the effects of castration are temporary and prostate cancer will progress to a “castration-resistant” disease (4). Progression to metastatic castration resistant prostate cancer (mCRPC) is often driven by the maintenance of AR-signaling despite the standard-of-care AR-targeted treatments such as castration and AR-antagonists like enzalutamide (5,6). Therefore, there is a critical need to define additional biochemical mechanisms that contribute to PCa progression, resistance to AR-targeted therapies, and to identify new therapeutic targets to manage this fatal disease.

An increasingly attractive approach for inhibiting AR signaling after failure of AR-targeted therapies has been to target the epigenetic co-regulators essential for its activity (7–9). Most epigenetic enzymes modify DNA or histones, often through methylation or acetylation, however some like p300, but can modify AR itself (10,11). Enzymes involved in the epigenetic control of gene expression, including p300 (12,13) and readers, like BET (Bromodomain and Extra-Terminal motif) proteins (14,15), are essential AR-coregulators and investigated targets in CRPC. While these approaches show promise, therapies directed at epigenetic regulators have not yet translated into significant clinical benefits. (16).

A new approach would involve targeting epitranscriptomic regulators, like those associated with N6-methyladenosine (m^6^A) RNA modifications. M^6^A is the most commonly observed, evolutionarily-conserved, reversible epigenetic modification found on both coding and non-coding transcripts (17); it plays a role in mRNA stability and translation (18,19) and may be important for a host of cellular functions such as increasing translation efficiency as well as regulating transcript degradation and sequestration (18,19). This post-transcriptional modification occurs transcriptome-wide at a consensus motif that is best represented as DRACH (D=A, G or U; H=A, C or U), with the center adenosine being modified (20). M^6^A modifications are also enriched in the 3’-untranslated-regions (UTRs) of mRNAs and can enhance protein binding and prevent microRNAs from targeting these transcripts (21–23). Furthermore, m^6^A modifications may be altered in cancer this alteration is frequently found on transcripts in pathways regulating cell growth, the cell cycle, the response to DNA-damage, and chromatin modification (24).

Similar to the other epigenetic processes, m^6^A modification has its own “writer,” “eraser,” and “reader” proteins. Of these proteins, the “writers” are composed of the proteins methyltransferase-like 3 (METTL3), METTL14, and Wilms tumor-1 associated protein (WTAP) – with METTL3 being the core catalytic subunit of the complex that methylates adenosine in RNA (25,26). METTL3, in particular, is emerging as an oncogene in a number of tumor types, including prostate cancer (27). METTL3 promotes the translation of many oncogenes in human cancer cell lines through modification of their corresponding mRNAs (28–31), and serves as a transcriptional co-activator by directly binding to chromatin as well as interacting with tissue-specific transcription factors at the transcriptional start sites of genes, which may ultimately facilitate target mRNA methylation (32). METTL3 is expressed at higher levels in prostate cancer compared to normal prostate tissue, other tumors, and tissue types, providing the first evidence that it contributes to oncogenesis in prostate cancer and role in the tissue-specific activity of AR (33,34).

METTL3 and m^6^A modification regulate a number of important oncogenes in prostate cancer, including members of the hedgehog signaling pathway (35), histone methylation (36), extracellular matrix proteins (37), a number of long-non-coding RNAs (lncRNAs) (38,39), LEF1 (40), AKT (41), and importantly MYC (34) and AR (42) itself. Regarding castration resistance, m^6^A contributes to guiding differential splicing and may control alterative splicing (43) such as events that create constitutively active AR-splice variants that lack its ligand binding domain (LBD) (44–50). However, m^6^A methylation on mRNA regulates translation of both oncogenes and tumor suppressors (51), and the functional and clinical significance of m^6^A modifications in prostate cancer is unclear (34,35,37,52–57).

METTL3 also functions independent of methylation, driving transcription and translation through binding interactions (58–60). METTL3 expression is elevated in primary prostate tumors compared to non-malignant prostate tissue (61) much like AR (62), and is overexpressed without reciprocal interaction of its partners in the methyltransferase complex, METTL14 or WTAP (33,34). Given its role as a transcriptional regulator in addition to its methyltransferase activity, METTL3 may interact directly with AR to promote its oncogenic functions. Here, we illustrate the non-catalytic, oncogenic role of METTL3 as an AR-transcriptional co-regulator.

## Methods

### Cell lines and culture

22Rv1 cells were obtained from ATCC (Manassas VA, USA). CWR-R1, VCaP, LAPC4, LNCaP, and their enzalutamide-resistant derivatives, in addition to BPH-1, 957E/hTERT, NCI-H660 (H660), LASCPC-01, HEK293, PC3, DU145, PNT-2, and RWPE1 cells, were generously provided by Dr. Vander Griend at the University of Illinois at Chicago and have been previously characterized and cultured as described (6,63,64). Dr. Peter Nelson at the Fred Hutchinson Cancer Center supplied LNCaP-shAR and LNCaP-APIPC cell lines (65). Enzalutamide resistant cell lines were cultured in ATCC RPMI (30-2001, ATCC) containing 10% FBS supplemented with 20 µM enzalutamide (MDV3100) (S1250, Selleck Chemicals). LNCaP NSC/shMETTL3 lines were cultured in ATCC RPMI with 10% FBS with 1µg/ml puromycin (A11138-03, Gibco). All cell lines were screened to confirm the absence of mycoplasma contamination utilizing an ATCC Universal Mycoplasma Detection Kit (30-1012K, ATCC). LNCaP cells overexpressing METTL3 were cultured in 3µg/ml blasticidin (A1113903, Invitrogen) and LNCaP-GFP were cultured with the addition of 1µg/ml puromycin. Combined LNCaP shMETTL3-NSC lines were cultured in 1µg/ml puromycin and 3µg/ml blasticidin in ATCC RPMI with 10% FBS. R1881 (metribolone) was purchased from (R0908, Sigma-Aldrich, St. Louis, MO) and stored at −20°C in ethanol and enzalutamide (MDV3100) were stored at −80°C in DMSO (D2438, Sigma-Aldrich), respectively. FBS was obtained from BioWest (S1480, Bradenton, FL).

### Cell Viability

Viability was assessed utilizing the CellTiter-Glo® Luminescent Cell Viability Assay kit (G7570, Promega, Madison, WI, USA) using the company provided protocol. Cells were seeded at 2500-5000 cells/well in 96 well white opaque plates (Corning, Cat: 07-200-336) and cultured for 5 days for drug-dose response experiments or to 7 days for cell proliferation experiments and results obtained on days 0, 1, 3, 5 and 7. shRNA transcription was induced with 1µg/ml doxycycline added on day 0 (Sigma-Aldrich, Cat: D3447). Luminescence measurements were obtained using the Synergy H1 Plate Reader (BioTek, Winooski, VT, USA). Readouts were adjusted to the day 0 readout to obtain adjusted luminescence values or further normalized to a control.

### Quantitative Reverse Transcription PCR (Q-RT-PCR)

Cells were cultured for 3 days as described above followed by isolation and purification of RNA using a Qiagen RNeasy Mini Kit (Qiagen, Valencia, CA, USA). RNA quality was evaluated using a NanoDrop 2000 Spectrophotometer (Thermofisher). The isolated RNA was subsequently reverse-transcribed to cDNA utilizing the High-Capacity cDNA Reverse Transcription Kit (Thermofisher, Cat: 4368814). Levels of the mRNAs for *METTL3*, *AR, ALOXE3*, *TLE1*, and *ILF2* were quantified utilizing Fast SYBR® Green Master Mix (Invitrogen) with custom primers (Integrated DNA Technologies, Coralville, IA) see **Table S1A** for sequences. Standard curves were used to verify primer efficiency and average change in threshold cycle (ΔCT) values determined for each sample relative to endogenous β-actin (*ACTB*) and compared to siNSC control (ΔΔCT). Analysis was performed using QuantStudio™ Real-Time PCR Software.

### Western Blots

Whole-cell lysates obtained from cells cultured in a 6-well plate (Becton, Dickinson and Company, Franklin Lakes, New Jersey) were subjected to lysis utilizing RIPA buffer containing 150 mM sodium chloride, 1.0% Igepal CA-630 (Sigma-Aldrich), 0.5% sodium deoxycholate, 0.1% SDS, 50 mM Tris, pH 8.0, supplemented with a protease/phosphatase inhibitor cocktail (Roche Molecular Biochemicals; Penzberg, Germany), followed by scraping and sonication (Fisher Scientific; Hampton, NH; model FB-120 Sonic Dismembrator). Protein concentration was determined using a BCA assay (Thermo-Fisher Scientific), 30µg of protein was diluted with ultrapure water (Corning) and 5x Lamelli buffer supplemented with 2-mercaptoethanol (444203, Millipore) to a mixture with a final concentration of 1x Laemmli buffer. Solutions were boiled at 95°C for 5 minutes, mixed, centrifuged, and electrophoresed. Membranes were blocked with a 5% milk blocking buffer in TBS. Antibodies employed included: anti-AR (D6F11 XP®, Cell Signaling Technology, Danvers, MA); anti-Beta Actin (AC-15, Sigma-Aldrich); anti-NKX3.1 [D2Y1A XP ^®^, Cell Signaling Technology, (Danvers, MA, USA) and anti-PSA (KLK3), (D11E1 XP^®^, Cell Signaling Technology); anti-METTL3 (EPR18810, Abcam); anti-METTL14 (PA5-117138, Invitrogen, Rockford, IL); anti-WTAP (ERP18744, Abcam), and anti-Flag (F1804, R&D, USA). Secondary antibodies; Licor IRDye 680RD Donkey Anti-Mouse Secondary Antibody (926–68072), Licor IRDye 800CW Donkey Anti-Rabbit Secondary Antibody (926-32213, LICORbio), and nitrocellulose membranes were obtained from Bio-Rad (1620115) and developed using the Licor Odyssey M system for both the 700 and 800nm channels (Lincoln, NE).

### siRNA Knockdown

Transient knockdown of METTL3 by siRNA transfection was performed using Silencer^®^ Select siRNAs (Invitrogen, Waltham, MA, USA with Silencer® and Select Negative Control No. 1 (43-908-43), siMETTL3-1 (s32141), siMETTL3-2 (s32142), or siMETTL3-3 (s32143) (Figure S1: Table S1B for sequences). Cells were plated in 6-well plates at 500,000 cells per well for protein and RNA analysis and in 96-well plates at 2500 cells/well for cell viability analysis. Immortalized Prostate Epithelial lines RWPE1 and 957E-hTERT cells were plated at 250,000 cells/well in a 6 well plate for protein/RNA analysis and 2000 cells/well in a 96 well plate for viability. The transfection of siRNAs was performed using Lipofectamine™ RNAiMAX (13778150, Invitrogen) following the manufacturer’s protocol. Transient knockdown of METTL3-overexpressing LNCaP cells was performed by siRNA transfection using siMETTL3-3, siMETTL3-4 (n322814) and siMETTL3-5 (n322813) and cell viability was measured as described above.

### STM2457 dose-response experiments

The efficacy of STM2457 (S9870, Selleck chem) on LNCaP, LNCaP Enz^R^, VCaP, VCaP-Enz^R^, CWR-R1, CWR-R1-Enz^R^, RWPE1, and 957E-hTERT viability was measured in using an initial concentration of 50µM and subsequent 1:5 dilutions with an equalized amount of vehicle control. Cells were plated at 2500 cells/well and STM2457 was added the following day with or without vehicle or enzalutamide. Cultures were equalized to contain the same concentration of vehicle, as both STM2457 and enzalutamide dissolved in DMSO. At the day 5 endpoint, viability was measured using the CellTiter-Glo® Luminescent Cell Viability Assay kit.

### m^6^A quantification

LNCaP or 957e-hTERT cells were treated with DMSO or STM2457, or Select Negative Control No. 1 (43-908-43), siMETTL3-1 (s32141), siMETTL3-2 (s32142) or siMETTL3-3 (s32143) (**Table S1B**). Cells with 3 biological replicates in 6 well plates were cultured using the method described above. RNA was harvested after 3 days. Using 5µg of total RNA, mature mRNA was isolated with the NEBNext® Poly(A) mRNA Magnetic Isolation Module kit (NEB #E3370) according to the provided protocol. The m^6^A levels on isolated mRNA were assayed using the EpiQuick m^6^A Methylation Quantification Colorimetric kit (P-9005-48) with 100ng of mRNA used per well. The readout was detected using the Synergy H1 Plate Reader (BioTek, Winooski, VT, USA) at 450nm.

### Generation of doxycyline-inducible shRNA and METTL3 overexpressing cell lines

Cells containing the doxycycline inducible shMETTL3 and control cell lines were generated via stable transduction of SMARTvector Inducible Lentiviral plasmids shNSC (VSC6570) and shMETTL3 Plasmids (sh-METTL3-1: V3SH11252-229451971, sh-METTL3-2: V3SH11252-229373728, sh-METTL3-3: V3SH11252-228569617) purchased from Horizon Discovery (Cambridge, UK). Tet-pLKO-puro-Scrambled control plasmids were a gift from Charles Rudin (Addgene plasmid # 47541; http://n2t.net/addgene:47541; RRID:Addgene 47541). Lentiviral plasmids were transfected into HEK293-T cells utilizing Lipofectamine 3000 with the expression vector plasmids, pCMV-dR8.2 (Addgene, #8455) and pVSV-G (Addgene, #8454). pCMV-VSV-G and pCMV-dR8.2 being a gift from Bob Weinberg (Addgene plasmid # 8454 ; http://n2t.net/addgene:8454 ; RRID:Addgene 8454; Addgene plasmid # 8455 ; http://n2t.net/addgene:8455 ; RRID:Addgene8455). Media containing virus particleswas filtered and used to infect LNCaP cells with 8 μg/ml polybrene (TR-1003-G, Sigma-Aldrich). Cells were subsequently selected with 1μg/ml puromycin. The parental LNCaP-METTL3 overexpressing cell lines (Table S1) were generated via stable transduction using the method described above into parental LNCaP cells: (eGFP-LV105, GeneCopoeia), or pLV[Exp]-Bsd-CMV>3xFLAG

/{hMETTL3[NM_019852.5](D395A)} (VB251009-1391vwr), pLV[Exp]-Bsd-CMV>3xFLAG

/{hMETTL3[NM_019852.5](D395A)} (VB251009-1508sru) or pLV[Exp]-Bsd-CMV>3xFLAG

/{hMETTL3[NM_019852.5](D395A,W398A)} (VB251009-1513wzv) (Vector Builder). METTL3 overexpressing cells were selected with antibiotic. For the METTL3 rescue study LNCaP cell lines were transduced with: pLX311-Cas9 control, the wild-type METTL3 and the D395A W398A plasmids mentioned above (Table S1). The pLX311-Cas9 plasmid was a gift from William Hahn & David Root (Addgene plasmid # 118018: http://n2t.net/addgene:118018 ; RRID:Addgene_118018).

### Co-Immunoprecipitation assays

LNCaP, LNCaP Enz^R^, and CWR-R1 cells were plated at 20 million cells per plate in a 15cm^3^ dish with one dish per immunoprecipitation. Cells were fixed in media containing a final concentration of 1% formaldehyde (ICN19404780, ThermoFisher) in the media and quenched with 1M glycine (A13816.36, ThermoFisher) with a final concentration of 125mM. Cells were then washed in 1x PBS (SH3025602, HyClone) and harvested. Cells were lysed using CellLytic M Lysis Reagent (C2978, Sigma Aldrich) and sonicated using the BioRupter® Pico Sonicator (B01080010, Diagenode, Denville, NJ, USA) as described in (66). Co-IP was done using Dynabeads Protein G IP Kit (10007D, ThermoFisher). Pulldown was performed using rabbit IgG control antibodies (Cell Signaling, Cat: 2729), anti-AR (PG-21 (Millipore, Burlington, MA, USA), and anti-METTL3 (Abcam ab195352, EPR18810, Cambridge, UK). Laemmli buffer with 2-mercaptoethanol was used to elute protein and were subsequently analyzed via Western blot.

### NanoBiT Assay

Similar to that performed in Budreika *et al*. 2025 (67), the NanoBiT^®^ (NanoLuc^®^ Binary Technology, Naperville, IL, USA) bioluminescent split luciferase assay, pcDNA3.1 vectors for AR, METTL3 and SOX2 (see **Table S1** for sequences) with either small or large BiT tags at their amino (N) and carboxy ©-terminus were sourced from GeneScript Biotech (Piscataway, NJ, USA). The NanoBiT^®^ Protein-Protein Interaction (PPI) Starter Kit (Promega) was used following the manufacturer’s protocol. HEK293-T cells were seeded at a density of 9000 cells per well in a 96-well clear bottom plates coated with poly-d-lysine (3603, Corning), isolated and purified BiT plasmids were co-transfected using Turbofectin 8.0 reagent (TF81001, OriGene) at a final concentration of 100ng plasmid/well for each plasmid. Cells were treated with DMSO or enzalutamide for 9 hours or starved with 10% CSS (S11650, R&D Systems) in Phenol-red free RPMI-1640 (11835030, Gibco) for 12 hours and then treated with EtOH or 1nM R1881 for 3 hours. Luminescence was quantified with a BioTek BioSpa Live Cell Analysis System (Agilent, Santa Clara, CA, USA). Treatments were performed in triplicate. Background signal was subtracted.

### Chromatin immunoprecipitation and sequencing (ChIP-seq)

LNCaP and 22Rv1 cells were plated and cultured at 4.1 million cells per dish for each IP in 15cm^3^ dishes and incubated at 37°C at 5% CO_2_ for 24 hours in 10% FBS ATCC modified RPMI. After 12 hours cells were treated with DMSO (Sigma-Aldrich, D2438) and then harvested after 24 hours of treatment. Cells were then fixed with 1% w/v formaldehyde for 8 minutes followed by quenching 125mM glycine for 5 minutes. Chromatin immunoprecipitation, size selection, and purification were performed according to manufacturer’s protocol using the Diagenode iDeal ChIP kit for Transcription Factors (Diagenode, C01010055). LNCaP cells were transduced with (shMETTL3-1 Catalog #: V3SH11252-229451971), (shMETTL3-2 Catalog #: V3SH11252-229373728) or (shMETTL3-3 Catalog #: V3SH11252-228569617) for doxycycline induced knockdown of METTL3. Cells were plated and cultured at 4.1 million cells per dish for each IP in 15cm^3^ dishes and incubated at 37°C at 5% CO_2_ for 24 hours in 10% ATCC modified RPMI. Cells were treated with DMSO (Sigma-Aldrich, D2438) or 1ug/ml doxycycline (D3447-500MG, Sigma-Aldrich) after 24 hours. After 48 hours cells were treated with ATCC modified RPMI phenol red free media supplemented with 10% charcoal stripped serum (CSS) while maintaining DMSO or doxycycline treatment. After 21 hours, cells were then treated with fresh ATCC modified RPMI phenol red free media with 10% CSS (BioWest, S11650) with or without 1 nM R1881 or equalized with EtOH (A962F-1GAL, Thermo Fisher) and cultured for 3 hours. Cells were then fixed 1% w/v formaldehyde for 8 minutes followed by quenching 125mM glycine for 5 minutes. Chromatin immunoprecipitation, size selection, and purification were performed according to manufacturer’s protocol using the Diagenode iDeal ChIP kit for Transcription Factors (C01010055, Diagenode). Cells were fixed with 1% w/v formaldehyde followed by quenching in 125mM glycine. Chromatin immunoprecipitation, size selection, and purification were performed according to manufacturer’s protocol using the Diagenode iDeal ChIP kit for Transcription Factors. Polyclonal rabbit anti-AR antibody (06-680, Millipore), METTL3 antibody (EPR18810, abcam) or the species-matched rabbit IgG control antibody (2729, Cell Signaling) were used in the immunoprecipitation. Chromatin shearing was carried out over 40 cycles (30 seconds on/30 seconds off) with a Diagenode Pico sonicator. A resulting size range of 100-500bp was confirmed by both 1% agarose DNA gel electrophoresis and Agilent 2100 bioanalyzer (G2939A, Agilent). The preparation of libraries and subsequent sequencing was conducted at the Roy J. Carver Biotechnology Center at the University of Illinois Urbana-Champaign (UIUC) utilizing the Ovation^®^ Ultralow System V2 DNA-Seq Library Preparation Kit (402666, Tecan). Libraries were combined, quantified via qPCR, and sequenced using a NovaSeq X Plus with V1.0 sequencing kits (NovaSeq). Fastq files were obtained using bcl-convert v4.1.7 Conversion Software (Illumina). Analysis was done using standard software including BWA MEM for alignment to reference human genome hg38 (68), MACS2 for peak calling https://pypi.org/project/MACS2/ (69), BEDTools for identifying regions of colocalization (70,71), https://meme-suite.org for motif analysis using the HOCOMOCOv11 core HUMAN mono meme format using MEME-ChIP version 5.5.7 (72), Cistrome-GO (73) identification of genes for which there is regulatory potential using the default cutoff of <10kb from Transcriptional Start Site (TSS). Lists of genes with an adjusted RP score of greater than or equal to 0.01 were identified for each condition and analyzed for Gene Ontology Biological Processes using GO Enrichment Analysis at GeneOntology.org (74,75) (https://geneontology.org). Track plots were visualized using IGV.

### Statistical Analysis

Statistical analysis was performed using GraphPad Prism software version 9.0 (GraphPad Software, La Jolla, CA, USA). Mean and mean standard error were determined for replicate qPCR samples, and one-way ANOVA was used to determine statistical significance. ΔCt values were relative to the control and data are presented as fold change (2^− ΔΔCt^) normalized to NSC. For cell viability data (n = 5 or 6) mean and SEM were determined, and one-way ANOVA was used to determine statistical significance. One-way ANOVA was used to determine significance for NanoBiT signal. Empiria studio v2.3 was used for densitometry calculations.

### Data and code availability

Datasets for this study have been deposited in the NCBI Gene Expression Omnibus (GEO) database and are available through accession number GSE335783. ChIP-seq analysis code was employed as previously described in Phoenix et al 2025 (76) and can be found at https://github.com/1jphoenix/Phoenix-SOX2-FOXA1.

## Results

### METTL3 is elevated in PCa cell lines compared to immortalized prostate derived cell lines and is higher when AR is higher

First, given its elevated mRNA expression in tumors (61), we evaluated the protein levels of METTL3, and its known partners in the m^6^A methyltransferase complex, METTL14 and WTAP, and compared it to AR in a panel of non-transformed (benign), AR-positive adenocarcinoma and AR-negative PCa cell lines (**Figure 1A**). Consistent with what was identified in patients (33,34), METTL3 is higher in the prostate cancer cell lines, and in AR-positive cells, METTL3 is higher where AR is also high. This also occurs when examining matched, isogeneic enzalutamide-resistant counterparts, when both AR and METTL3 are elevated (**Figure 1B**), consistent with a potential role of METTL3 in AR-driven PCa.

**Figure 1.**
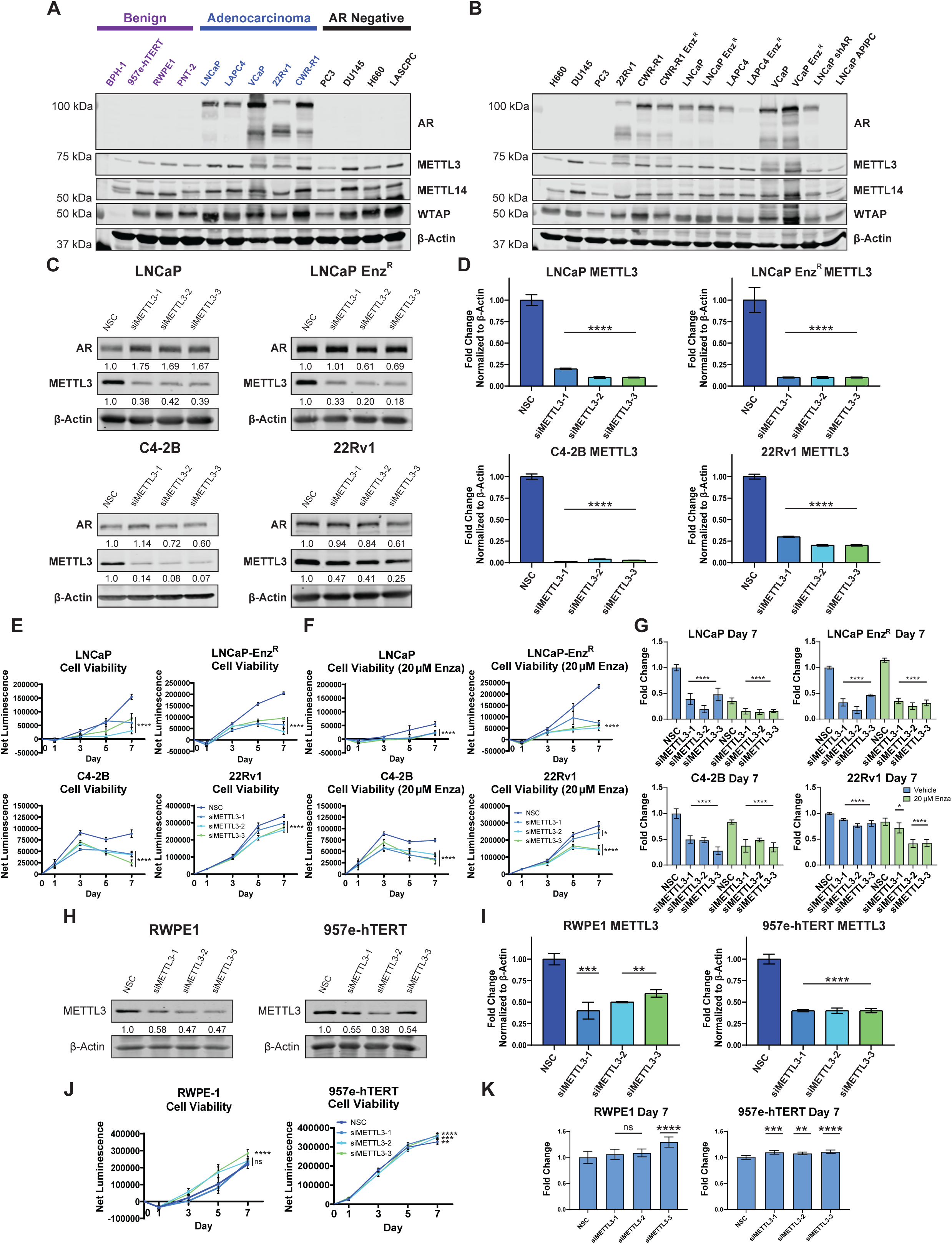
METTL3 is elevated in PCa cell lines, and METTL3 siRNA knockdown inhibits proliferation in PCa cell lines, but not in immortalized non-transformed prostate epithelial cells. **A)** Western blot of AR, METTL3, METTL14 and WTAP protein in non-transformed, immortalized epithelial cells (benign), AR-positive prostate cancer cells (adenocarcinoma) and AR-negative prostate cancer cell lines with β-actin as a loading control. **B)** Western blot of AR, METTL3, METTL14 and WTAP protein in prostate cancer cells, both enzalutamide naïve and resistant. **C)** Western blots of METTL3 confirming knockdown by three siRNAs directed toward METTL3 in LNCaP, LNCaP Enz^R^, C4-2B, and 22Rv1 cell lines. **D)** Expression of METTL3 mRNA in LNCaP, LNCaP Enz^R^, C4-2B, and 22Rv1 cell lines quantified by qPCR after transfection of METTL3-targeting siRNAs. Expression is relative to non-silencing control (NSC) and calculated using the ΔΔCt method. **E-F)** Cell viability of prostate adenocarcinoma cell lines LNCaP, LNCaP Enz^R^, C4-2B, and 22Rv1 after transfection of METTL3-targeting or NSC siRNAs over a 7-day time course in the presence or absence of 20 µM enzalutamide. Viability was evaluated using the CellTiter-Glo® Luminescent Cell Viability Assay kit (Promega, Madison, WI, USA). **G)** Cell viability of LNCaP, LNCaP Enz^R^, C4-2B, and 22Rv1 cells at day 7 after transfection with METTL3-targeting or NSC siRNAs with or without enzalutamide (20 µM) treatment**. H)** Western blots demonstrating knockdown of METTL3 by three siRNAs directed toward METTL3 in the immortalized prostate epithelial cell lines RWPE1 and 957e-hTERT. **I)** Expression of METTL3 mRNA in the benign prostate cell lines RWPE1 and 957e-hTERT quantified by qPCR after transfection of METTL3-targeting siRNAs. Expression is relative to NSC and calculated using the ΔΔCt method. **J)** Cell viability of the benign prostate cell lines RWPE1 and 957e-hTERT after transfection of METTL3-targeting or NSC siRNAs over a 7-day time course. Viability was evaluated using the CellTiter-Glo® Luminescent Cell Viability Assay kit (Promega, Madison, WI, USA). **K)** Cell survival the benign prostate cell lines RWPE1 and 957e-hTERT at day 7 after transfection with METTL3-targeting or NSC siRNAs.

### Reducing METTL3 inhibits proliferation in malignant cell lines, but not in immortalized prostate epithelial cells

A functional phenotypic role of METTL3 in PCa cells and benign controls was evaluated by measuring the effect of METTL3 depletion on proliferation. Using siRNA knockdown of METTL3, significant depletion of METTL3 in LNCaP, LNCaP Enz^R^, C4-2B, and 22Rv1 (**Figure 1C-D**) and VCaP, CWR-R1 and CWR-R1 Enz^R^ (**Figure S2H-M**), was evident both at the mRNA (**Figure 1D**, **S2A-C**) and protein levels (**Figure 1D**, **S2E-G**). Concordantly, METTL3 depletion results in reduced proliferation in PCa cell lines, with significant decrease in proliferation observed on day 5 or 7 in LNCaP, LNCaP Enz^R^, C4-2B, 22Rv1, VCaP, CWR-R1 and CWR-R1 Enz^R^ (**Figure 1E-G**; **S2H-M**). The effects of METTL3 knockdown were additive to growth inhibition observed with enzalutamide treatment (**Figure 1F-G**; **S2K-M**).

In contrast, when METTL3 was knocked-down in the AR-negative, immortalized epithelial cell lines RWPE1 and 957e-hTERT (**Figure 1H-I**), proliferation was unaffected (**Figure 1J-K**). Conversely, there was a modest, yet statistically significant increase in proliferation observed when METTL3 was reduced on day 7 (**Figure 1J-K**). These results indicate that METTL3 can affect the proliferation of AR-positive PCa cells, and not in benign controls.

The effect of METTL3 knockdown in LNCaP cells using a doxycycline inducible model to deplete METTL3 via shRNA was also determined, allowing for strong knockdown that could be obtained with desired timing and performed at scale for other downstream analyses. Doxycycline induced METTL3 shRNA knockdown was achieved at both the protein (**Figure 2A**) and mRNA levels using this approach (**Figure 2B**). The shRNA knockdown of METTL3 significantly reduced cell viability compared to controls (**Figure 2C-D**), and consistent with the siRNA experimental results (**Figure 1F**), the effect was additive with enzalutamide treatment (**Figure 2C-D**).

**Figure 2.**
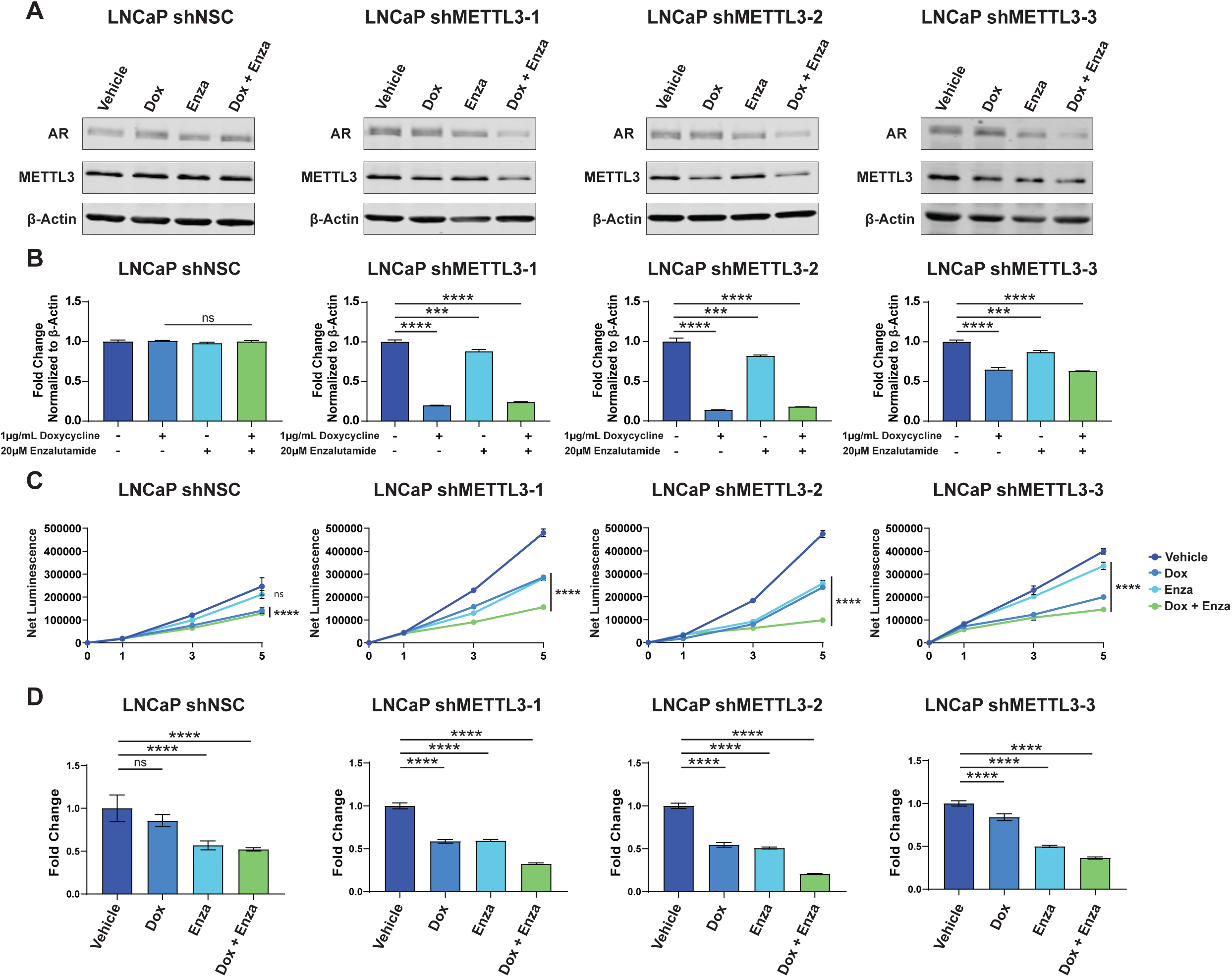
Doxycycline-induced shRNA knockdown of METTL3 reduces cell proliferation in LNCaP cells and is additive with enzalutamide. LNCaP cells were stably transduced with doxycycline-inducible shRNA targeting METTL3. **A)** Western blots demonstrating knockdown of METTL3 by 1µg/ml doxycycline treatment in LNCaP cells stably transduced with doxycycline-inducible shRNA targeting METTL3 compared to NSC. Cells were cultured with or without enzalutamide (20 µM). **B)** Expression of METTL3 mRNA in LNCaP cells stably transduced with doxycycline-inducible shRNA targeting METTL3 and cultured with or without doxycycline (1µg/ml) quantified by qPCR. Cells were also cultured with or without enzalutamide (20 µM). Expression is relative to NSC and calculated using the ΔΔCt method. **C)** Cell viability over a 5-day time course of LNCaP cells stably transduced with doxycycline-inducible shRNA targeting METTL3 and cultured with or without doxycycline (1µg/ml). Cells were also cultured with or without enzalutamide (20 µM). Viability was evaluated using the CellTiter-Glo® Luminescent Cell Viability Assay kit (Promega, Madison, WI, USA). **D)** Cell survival in LNCaP cells stably transduced with doxycycline-inducible shRNA targeting METTL3 and cultured with or without doxycycline (1µg/ml) and/or enzalutamide (20 µM).

### METTL3 enzymatic inhibition reduces m^6^A modification with marginal impact on the proliferation of both benign and malignant PCa lines

To understand if the effects of METTL3 on PCa cell viability are dependent on its enzymatic activity, we compared the results of knocking down METTL3 to the effect of inhibiting its enzymatic function with the METTL3 inhibitor STM2457. STM2457 is a potent S-adenosyl methionine (SAM)-competitive inhibitor of METTL3 catalytic activity with high affinity (1.4 nM Kd) and an IC_50_ of 16.9 nM (77). Unlike depletion of METTL3 by siRNA knockdown, treatment of several PCa cell lines with STM2457 showed only minimal effect on proliferation except at high doses approaching the limits of solubility (**Figure 3A for LNCaP, LNCaP-Enz^R^; S3A-B for VCaP, VCaP-Enz^R^, CWR-R1, CWR-R1-Enz^R^**). That lack of an impact of STM2457 on proliferation was evident for both transformed and non-transformed cells as well as enzalutamide-resistant cells (**Figure 3A RWPE-1, 957E/hTERT; S3A-B**). The lack of an effect seen with STM2457 occurred even though inhibition of mRNA adenosine methylation was observed in both the PCa cell line LNCaP and the benign cell line 957e-hTERT (**Figure 3B-C**). The level of m^6^A inhibition by STM2457 was comparable to the inhibition seen when depletion of METTL3 was achieved by treatment with siRNA (**Figure 3B-C**).

**Figure 3.**
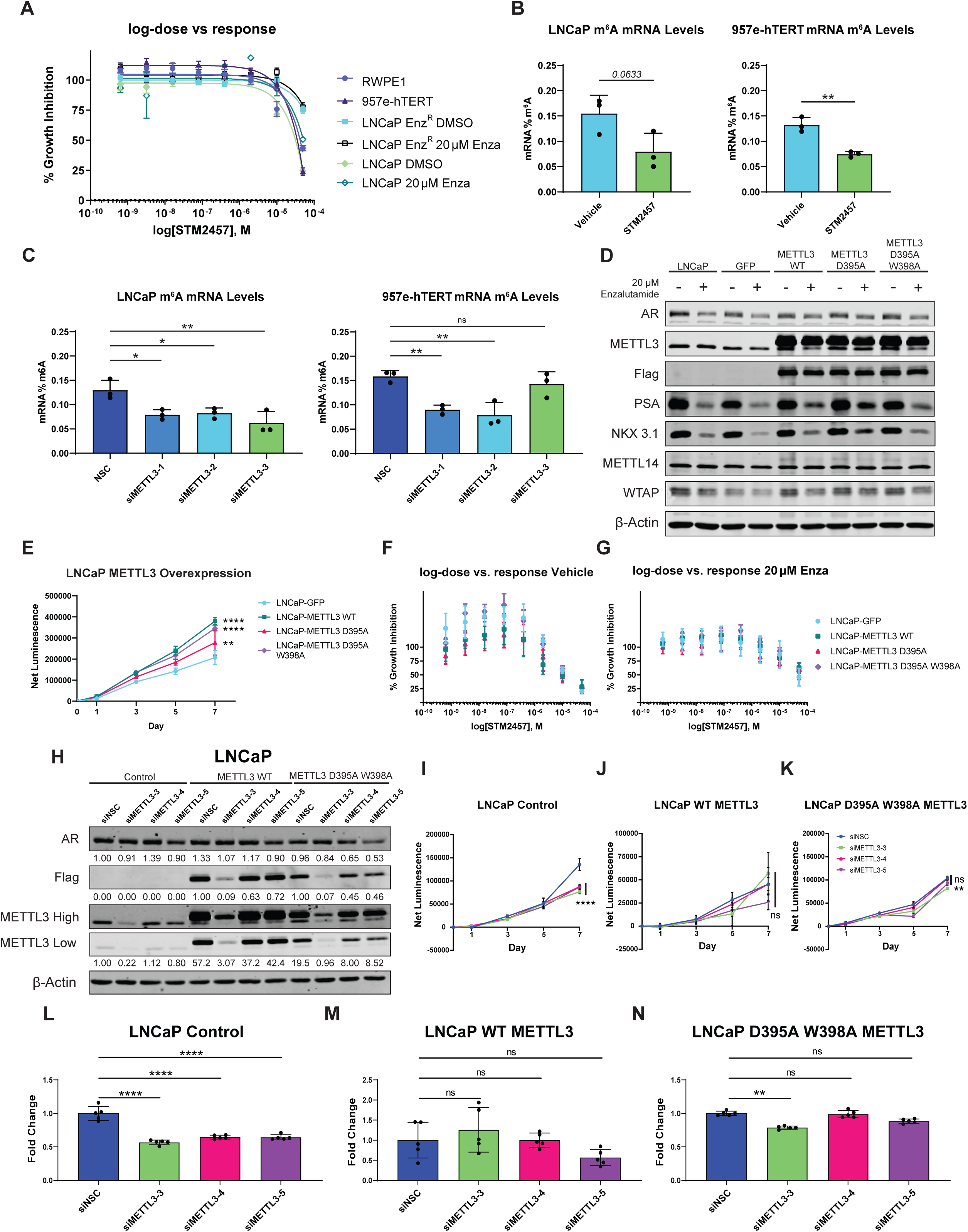
METTL3 enzymatic inhibition has little impact on proliferation on both benign and malignant PCa lines despite inhibition of m^6^A. **A)** Comparison of dose response curves of METTL3 inhibitor STM2457 in prostate adenocarcinoma cell lines and benign prostate cell lines in the luminescence cell viability assay at day 5. **B)** Effect of METTL3 inhibitor STM2457 on mRNA m^6^A levels in LNCaP and 957e-hTERT cell lines. Levels of m^6^A in mRNA was measured using the EpiQuick m^6^A Methylation Quantification Colorimetric kit. **C)** Effect of siRNA knockdown of METTL3 on mRNA m^6^A levels in LNCaP and 957e-hTERT cell lines. Levels of m^6^A in mRNA was measured using the EpiQuick m^6^A Methylation Quantification Colorimetric kit. **D)** LNCaP cells were transduced with FLAG-tagged WT METTL3 or catalytically inactive mutants (D395A and D395A/W398A), with GFP lentivirus as a control. Protein expression was assessed by Western blot using antibodies against METTL3, FLAG, AR, PSA, NKX3.1, METTL14, and WTAP, with β-actin as a loading control. AR modulation was evaluated by culturing cells in the presence or absence of enzalutamide (20 µM). **E)** Cell viability of LNCaP cells overexpressing FLAG-tagged WT METTL3 or catalytically inactive mutants (D395A and D395A/W398A) was evaluated over a 7-day period using the CellTiter-Glo® Luminescent Cell Viability Assay (Promega). **F-G)** The dose–response curve for the METTL3 inhibitor STM2457 was assessed in cells overexpressing FLAG-tagged WT METTL3 or catalytically inactive mutants (D395A, D395A/W398A), in the presence or absence of enzalutamide (20 µM). Cell viability rescue from shRNA knockdown of METTL3 by overexpression of METTL3. H-M) LNCaP cells stably transduced with doxycycline-inducible shMETTL3-1, shMETTL3-2 or shNSC (control) and treated with or without doxycycline were subjected to overexpression METTL3 by transduction of WT METTL3, catalytically inactive mutant (D395A/W398A) or vector control (Cas9). Cell viability was evaluated over a 7-day period (**H-J**) using the CellTiter-Glo® Luminescent Cell Viability Assay (Promega). Cell survival at day 7 (**K-M**) is shown relative to vector control.

Overexpression of either wild-type or catalytic inactive mutant METTL3 rescues the anti-proliferative effects of endogenous METTL3 knockdown.

Given the comparable sensitivities to STM2457 and loss of m6A modification with either STM2457 or siRNA knockdown, it was hypothesized that the catalytic function of METTL3 was dispensable for its effect on PCa proliferation reported above. To evaluate the requirement of the catalytic function of METTL3 on cell proliferation, FLAG-tagged WT METTL3 along with two catalytically inactive mutants (*METTL3* D395A, which disrupts most catalytic activity, and *METTL3* D395A, W398A, which disrupts activity and m^6^A binding (31)) were over-expressed in LNCaP cells via lentiviral transduction (**Figure 3D**). Enzalutamide treatment alone resulted in slightly decreased METTL3 expression. METTL3 overexpression had little effect on AR levels or on AR targets PSA and NKX 3.1. There was no effect on expression of METTL14 with METTL3 overexpression with or without enzalutamide treatment. Compared to the GFP control, METTL3 overexpression slightly increased WTAP while enzalutamide slightly decreased WTAP expression (**Figure 3D**). Overexpression of METTL3, in both WT and inactive mutants, increased cell viability compared to the GFP control in LNCaP (**Figure 3E**). Overexpression of METTL3 (WT or mutants) had no impact differences in viability or sensitivity to STM2457 in LNCaP cells treated with either vehicle or enzalutamide (**Figure 3F-G**).

To determine if overexpression of WT or mutant METTL3 could rescue the effects on cell viability seen with siRNA knockdown, we utilized two more siRNAs that targeted an intron of METTL3 (**Figure S1A**). It was first determined that targeting the intron reduced endogenous METTL3 without affecting the levels of the ectopically expressed protein (**Figure 3H; S3C-E**). We then compared these siRNAs to the growth of siRNA-3 in METTL3 over-expressed cells and determined that either wild-type or catalytic dead mutant METTL3 overexpression rescued the effects of the siRNA knockdown of endogenous METTL3 (**Figure 3I-N**). These data demonstrate the functionality of METTL3 independent of its catalytic activity, for the proliferation of AR-positive prostate cancer cells and not AR-negative immortalized prostate epithelial cells.

### METTL3 and AR protein physically interact

Given the key role of AR in PCa proliferation and progression, and sensitivity to METTL3 knockdown but not to enzymatic inhibition in AR-positive cancer cell lines, we investigated the direct binding interaction between METTL3 and AR. METTL3 and AR coimmunoprecipitate (Co-IP) in multiple AR-positive PCa cell lines. We observed CoIP of METTL3 and AR in LNCaP, LNCaP Enz^R^ and CWR-R1 (**Figure 4A-C**; **Figure S2A**). To further validate the interaction, we utilized a NanoBiT ^®^ bioluminescent protein-protein split-luciferase reporter system to confirm the METTL3/AR interaction as well as to evaluate the effect of modulating the AR activity with the synthetic androgen R1881 or enzalutamide antagonism. We measured increased luminescence in cells co-transfected with plasmids for AR and METTL3 NanoBiT fusion proteins, with highest signal coming from the N-terminus of AR interacting with the C-terminus of METTL3, regardless of hormone manipulation (**Figure 4D-G**). Taken together, here are the first data illustrating an interaction between AR and METTL3.

**Figure 4.**
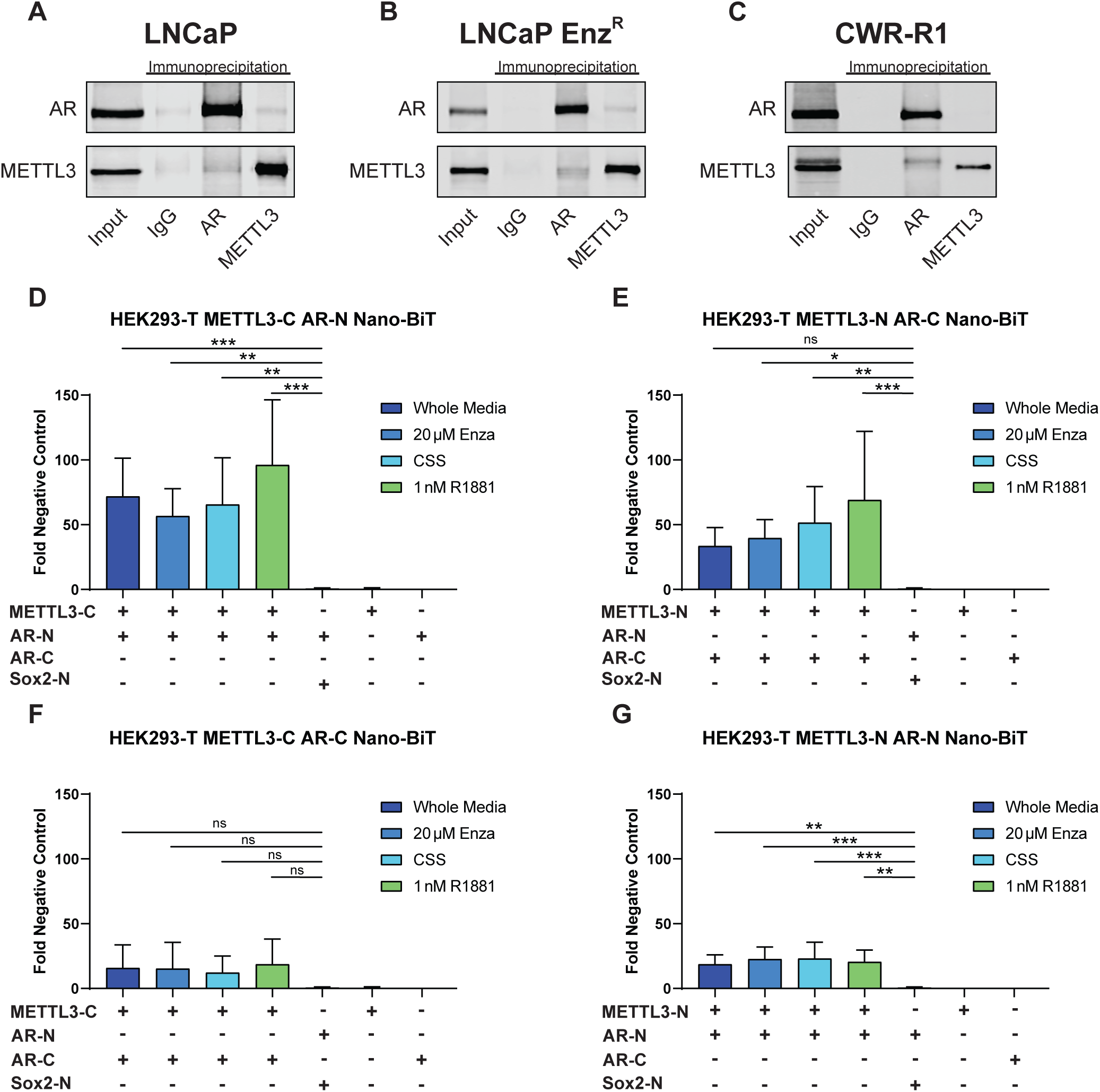
METTL3 and AR show evidence of interaction. **A)** Western blot demonstrating co-immunoprecipitation of AR and METTL3 in LNCaP, LNCaP Enz^R^ and CWR-R1 cell lines using IgG antibodies as a negative control. **B)** Luminescence relative to negative control from NanoBiT® assays in HEK293-T cells evaluating the interaction of LgBiT and SmBiT fragments fused to AR, METTL3, and/or Sox2 (negative control) at either the N- or C-terminus. Cells were cultured in complete media or under AR-modulating conditions [charcoal stripped serum (CSS), enzalutamide (20 µM), or R1881 (1 nM)].

### METTL3 and AR colocalize on chromatin throughout the genome

To functionally validate our findings of a protein-protein interaction between METTL3 and AR, we performed ChIP-seq AR and METTL3 from LNCaP and 22Rv1 cells (**Figure 5**), as METTL3 has previously been identified to act as a transcriptional co-regulator on the chromatin (32). METTL3 and AR colocalized (overlap of at least 1 base pair) at 53% of METTL3 IP peaks in LNCaP (**Figure 5A**) and 43% in 22Rv1 (**Figure 5E**). DNA-binding motif discovery (**Figure 5B-D; 5F-H**, see **Supplemental MEME Output Motif Files**) utilizing the MEME-ChIP Suite for METTL3 and AR ChIP-seq peaks indicates that METTL3 bound to motifs in LNCaP cells (**Figure 5B**) of nuclear hormone receptors such as the Vitamin D Receptor (VDR) and NF2FL2 or NRF2, the antioxidant and redox master transcriptional regulator, both important for prostate cancer biology (78,79). We also identify a sequence (NA) that is reminiscent of the DRACH RNA-binding motif (D = A/G/U, R = A/G, H = A/C/U) that METTL3 utilize at m^6^A targeted sequences (20). As expected, AR and its pioneering cofactor Forkhead box A1 (FOXA1) (80,81), as well as other typical nuclear hormone and NR3C AR-like motifs (GR, PR, ER) (82) were also identified in our LNCaP-AR ChIP-seq peaks (**Figure 5C**). The co-bound AR and METTL3 peaks are enriched for E2F motifs as well (**Figure 5D**), indicating a likely a role in controlling proliferation (83).

**Figure 5.**
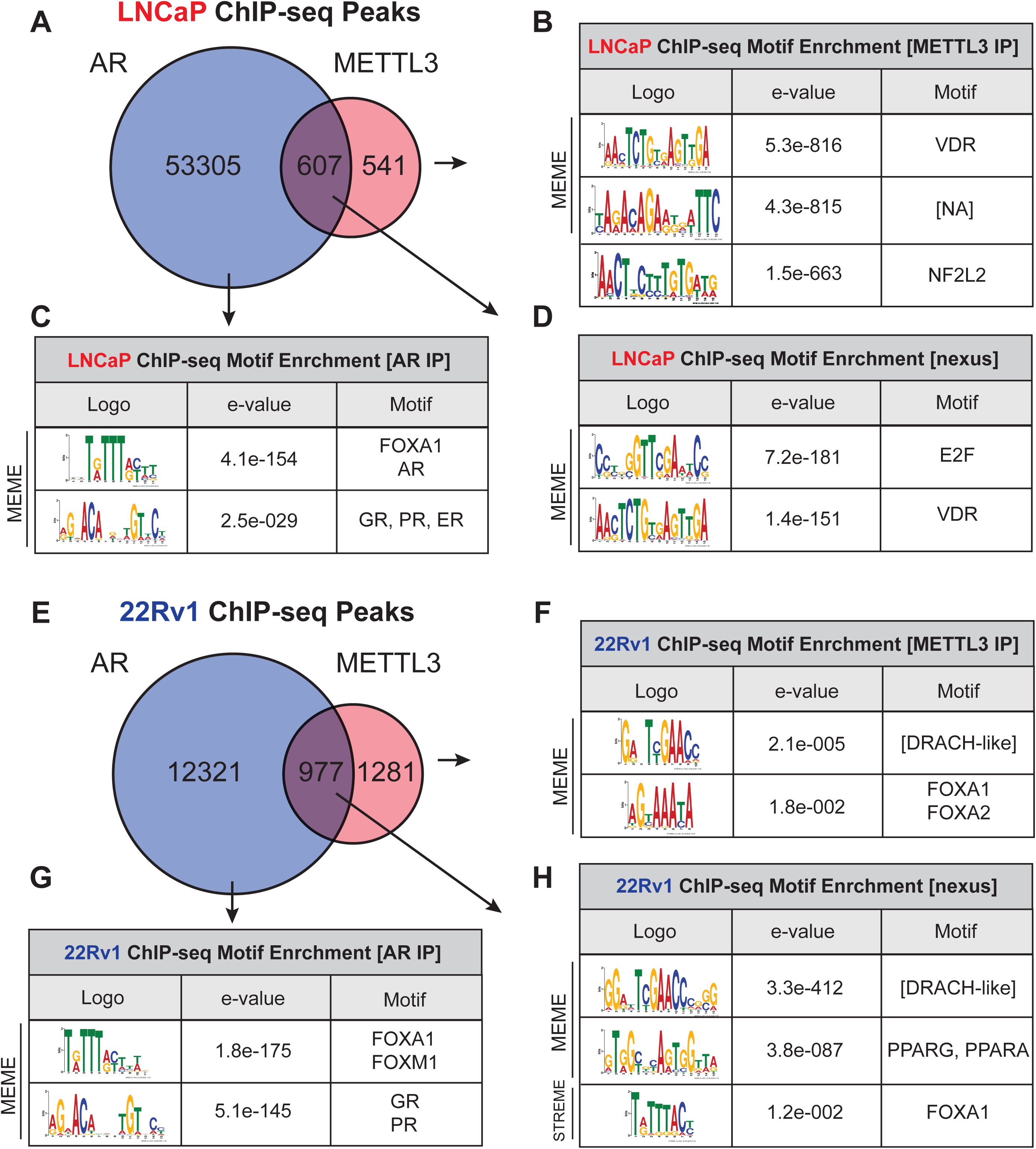
METTL3 and AR co-occupy sites on DNA in LNCaP and 22Rv1 prostate adenocarcinoma cells. **A)** Venn diagram of AR and METTL3 ChIP seq peaks in LNCaP cells grown in whole media. Shared regions signify called peaks with >= 1bp of overlap. **B-D)** Binding motif enrichment near called ChIP-seq peaks of AR and METTL3 in LNCaP cells. Both MEME and STREME motif discovery analyses were included. Full lists of motifs discovered and run parameters can be found in Supplemental Figure 5. **E)** Venn diagram of AR and METTL3 ChIP seq peaks in 22Rv1 CRPC cells grown in whole media. Shared regions signify called peaks with >= 1bp of overlap. **F-H)** Binding motif enrichment near called ChIP-seq peaks of AR and METTL3 in 22Rv1 cells. Both MEME and STREME motif discovery analyses were included.

Consistent with what we identified in LNCaP cells, we find a true-DRACH-like motif as well as FOXA1 motifs from METTL3-bound DNA, in both peaks identified without AR (**Figure 5F**), as well as co-localized with AR (**Figure 5H**), implicating direct interactions with AR and its other coregulators such as FOXA1. And as with LNCaP cells (**Figure 5C**), the same FOXA1 and NR3C AR-like motifs with AR alone (**Figure 5G**) were identified. Taken together, these results illustrate that METTL3 displays DNA sequence specificity with similarities to its known RNA-binding motif recognized by the methyltransferase complex and co-occupies many of the same sites that AR binds in prostate cancer cells, which are enriched for FOXA1 motifs.

To identify genes that may be regulated by colocalized binding of METTL3 and AR, we analyzed the METTL3 IP and AR IP peaks bed files utilizing CistromeGO for genes with a regulatory potential (RP) score ≥0.01 (see **Supplemental Table 2** for full list). In LNCaP (**Figure 6A**) and 22Rv1 (**Figure 6B**), 660 and 538 genes were identified respectively. Gene Ontology analysis of these shared regulatory potential genes in LNCaP cells identified biological processes relating to protein localization to the chromosome, hormone metabolic processes and reactive oxygen species/lipoxygenase pathways (**Figure 6C**). In 22Rv1 cells (**Figure 6D**) biological processes relating to T-cell differentiation, as well as processes relating to signal transduction by P53, the TGF-β signaling pathway and the MAPK cascade were identified for AR/METTL3 shared regulatory potential genes. AR/METTL3 shared Biological Processes common to both LNCaP and 22Rv1 included processes relating to protein localization to the chromosome and T-cell activation and differentiation (**Figure 6E-F**).

**Figure 6.**
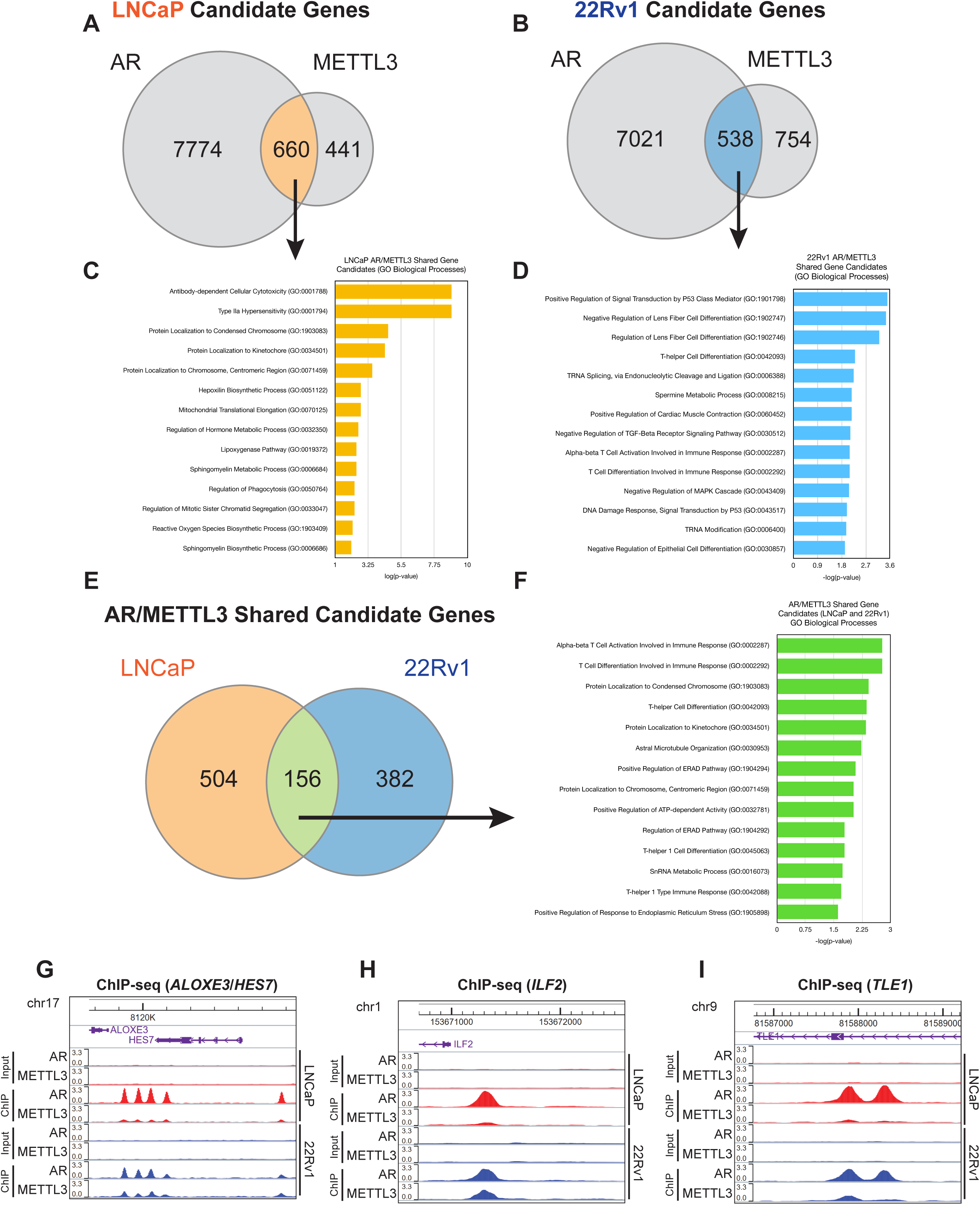
METTL3 and AR co-occupy sites on DNA near genes involved in several biological processes. A-B) Venn diagram of AR and METTL3 potential target genes in LNCaP and 22Rv1, respectively. ChIP-seq peaks were analyzed using Cistrome-GO to identify genes that had AR and METTL3 called binding peaks less than 10kb from a TSS and an adjusted regulatory potential (RP) score of >0.01. **C-D)** Gene Ontology (GO - Biological Processes) analyses for the shared AR and METTL3 candidate target genes in LNCaP and 22Rv1, respectively. GO enrichment was performed using Enrichr. **E)** Venn diagram of the LNCaP and 22Rv1 candidate gene nexuses. **F)** Gene Ontology (GO - Biological Processes) analyses for the shared AR and METTL3 candidate target genes shared in both LNCaP and 22Rv1 cell lines. **G-I)** ChIP-seq track plots of AR and METTL3 binding in both LNCaP and 22Rv1. Peaks were visualized using the WashU Genome Browser v61.1.0 and mapped to the human hg38 chromosome build.

Next, we assessed protein coding genes near colocalized binding of METTL3 and AR in both 22Rv1 and LNCaP as candidate target genes for their regulation. We focused on a key genes with a high regulatory potential score that have been implicated in cancer development and progression. Colocalized binding of METTL3 and AR occurs near *ALOXE3*, which had the highest RP score in 22Rv1 (**Figure 6G**). Other genes with oncogenic potential such as *ILF2* (**Figure 6H**)*, TLE1* (**Figure 6I**), and *DPP9* also demonstrated colocalized binding of METTL3 and AR.

Next, we evaluated the effect of METTL3 knockdown on expression of these genes by qPCR in LNCaP cells that had been stably transduced with doxycycline (dox) inducible shRNA (**Figure S5**). Expression of *ALOXE3* (**Figure S5A**) and *ILF2* (**Figure S5B**) is decreased by METTL3 knockdown with doxycycline treatment to a similar extent as seen by the addition of enzalutamide. Expression of *ALOX3E* was decreased by the combination of METTL3 knockdown and enzalutamide treatment compared to either doxycycline or enzalutamide alone, indicating these are AR and METTL3 upregulated gene targets. Conversely, enzalutamide treatment increases the expression of *TLE1* (**Figure S5C**), METTL3 knockdown similarly increases *TLE1*-expression, indicating that AR is a negative regulator of *TLE1* and METTL3 is essential to this regulation. Taken together, these results illustrate direct positive (*ALOX3E*, *ILF2*) and negative (*TLE1*) co-regulation of gene targets of the AR-METTL3 complex.

### METTL3 knockdown reduces the AR-bound cistrome

To evaluate the effects of METTL3 knockdown on AR chromatin binding, we utilized our doxycycline inducible shMETTL3-LNCaP cells and performed ChIP-seq on AR and METTL3 with or without doxycycline treatment and under hormone-starved then androgen (R1881) - stimulated conditions. We cultured two of our sh-METTL3 LNCaP lines, sh-1 and sh-2 (**Figure 2** and for sh-sequences see **Figure S1**) cells the presence or absence of doxycycline for 72 hours and in charcoal stripped media for 48 hours, followed by stimulation with 1 nM R1881 for 3 hours. These conditions result in the modulation of both androgen and METTL3 levels prior to chromatin immunoprecipitation.

METTL3 knockdown successfully achieved in dox treated cells (**Figure S4D-E**). As before, we observed colocalized binding of METTL3 and AR with >50% of METTL3 binding peaks colocalized with AR binding under all conditions (**Figure 7A-B**). In general, there is increased AR-bound ChIP-seq peaks with the addition of the AR agonist as expected, and fewer METTL3 peaks were recovered, again indicating minor direct interaction with the DNA. However, in androgen stimulated R1881 treated cells, dox treatment to deplete METTL3 greatly reduced AR ChIP-seq peaks (**Figure 7A-B**). These effects are more apparent in heatmaps (**Figure 7C-D**) which show AR binding at the top 10000 AR peaks. There is increased AR cistromic occupancy with the addition of the agonist R1881, with or without doxycycline (METTL3 knockdown). In the METTL3 ChIP-seq, there is also increased METTL3 binding with the addition of R1881 (**Figure 7E-F**) in METTL3 heatmaps. These data illustrate that decreases of METTL3 leads to reciprocal decreases in AR chromatin binding, providing evidence of METTL3 as an essential AR-coregulator in PCa cells.

**Figure 7.**
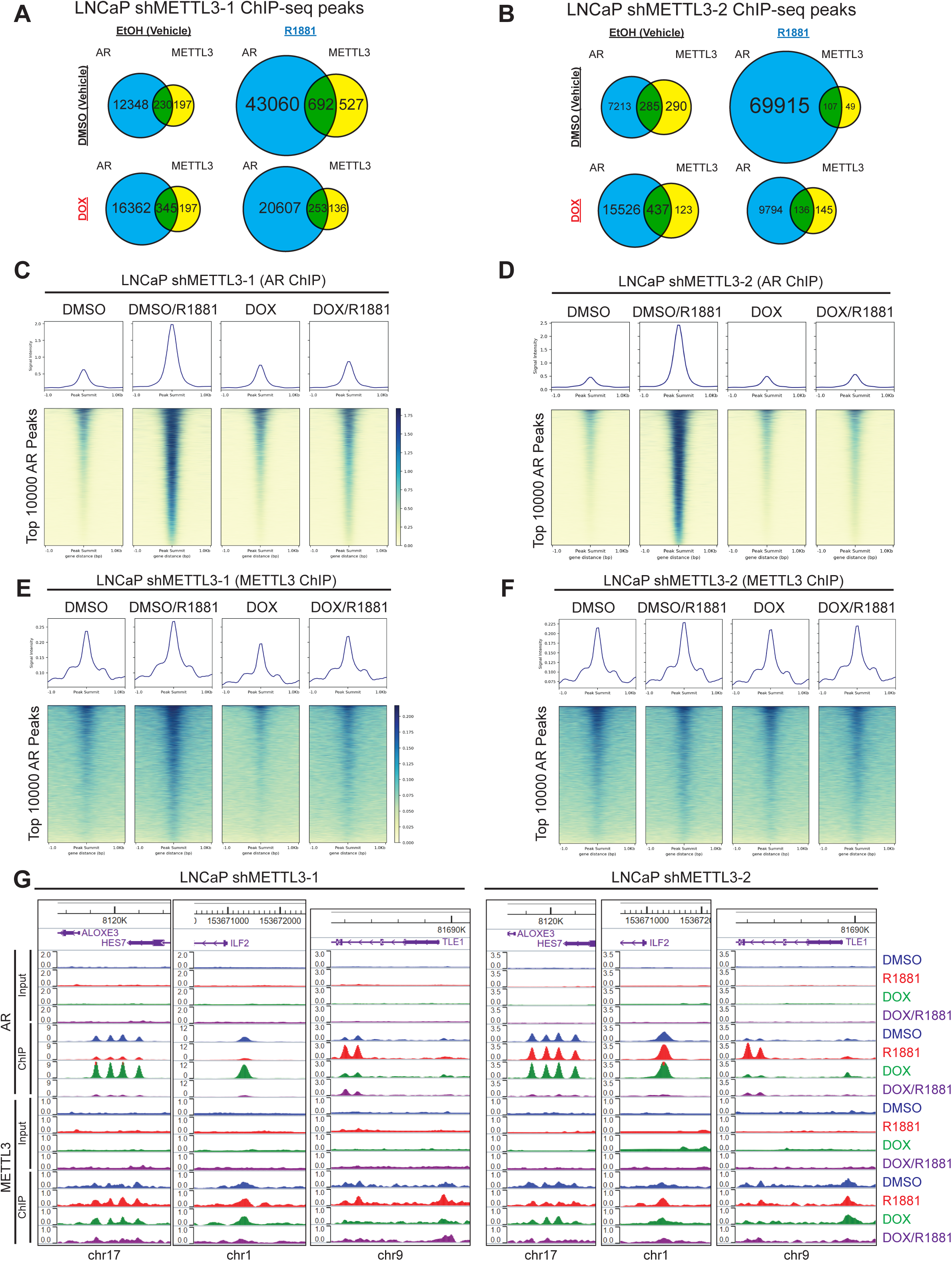
Inducible knockdown of METTL3 in LNCaP cells results in a decreased AR cistrome. A-B) Venn diagrams representing AR and METTL3 ChIP-seq binding peaks in LNCaP cells undergoing AR pathway and METTL3 modulation. **C-F)** Heatmaps of AR and METTL3 DNA binding in LNCaP cells. The vertical axis of each heatmap represents the top 1000 AR binding sites by peak score in both clones of LNCaP cells. **G)** ChIP-seq track plots of AR and METTL3 binding in LNCaP cells undergoing AR pathway and METTL3 modulation. Peaks were visualized using the WashU Genome Browser v61.1.0 and mapped to the human hg38 chromosome build.

When at individual candidate AR-METTL3 target genes (**Figure 7G**); METTL3 knockdown in hormone-starved conditions (in the absence of AR agonist) did show increased AR binding at *ALOXE3* and *ILF2*, but whereas AR binding at *TLE1* was increased with R1881 treatment (**Figure 7G**). METTL3 knockdown in the presence of R1881 decreased AR binding at all 3 genes (**Figure 7G**). For AR target genes *KLK3*, *TMPRSS2*, and *C1orf116* (SARG) (82) there is minimal METTL3 binding, but as expected, AR binding is increased with the addition of R1881 alone (**Figure S6**). However, METTL3 knockdown eliminates the AR binding observed with R1881 (**Figure S6**). These data indicate that METTL3 may also be necessary for the broader AR-mediated transcriptional complex without directly binding to the chromatin itself. Taken together, these results illustrate the importance of METTL3 in regulating the AR-cistrome and establishes METTL3 as an AR-transcriptional co-regulator essential for prostate cancer cell survival.

## Discussion

Here, we present the first work illustrating the direct interaction of METTL3 and AR and its importance in prostate cancer cells. METTL3 has various functions outside of the m^6^A methyltransferase complex, and we have identified another function as an AR-coregulator. These results rectify the somewhat confusing roles of METTL3 in prostate cancer. While it is increased transcriptionally in primary prostate cancers (33), it is initially decreased in enzalutamide resistance (37), which would mirror the levels of AR itself in PCa (84). Various groups have reported METTL3 to have both tumor suppressive and oncogenic functions *in vitro* and differential clinical correlations with both m^6^A RNA modifications and METTL3 expression suggest that there is a difference between the importance of the enzyme activity versus protein. Given the dispensable nature of METTL3 in the non-malignant prostate cancer cell lines and the essentiality in AR-positive PCa cells, we have illustrated an oncogenic role for it as an important AR-interactor.

We identified a few genes regulated by the AR-METTL3 complex that may have translational relevance: *ALOX3*, *ILF2*, and *TLE1*. Arachidonate epidermal lipoxygenase 3 (*ALOX3*) is an enzyme whose expression predicts poor prognosis in colon cancer (85) and has been linked to ferroptosis (86). Interleukin Enhancer-binding Factor 2 (*ILF2*) is highly expressed in prostate cancer and is implicated in poor clinical outcomes (87). These two genes were putative oncogenes we found upregulated by AR and METTL3. Finally, a repressed gene, Groucho (Gro)/Transducin-like enhancer of split-1 (*TLE1*) has mixed roles as a transcriptional repressor in multiple tumor types (88). Future work will be focused on the overall role of these targets, and other downstream targets of the AR-METTL3 complex, and validation of other genes and pathways we identified through ChIP-seq analysis.

Our results of this noncanonical role of METTL3 outside of its enzymatic activity have broad implications for the clinical relevance of targeting METTL3. First, the lack of efficacy of enzymatic inhibition in prostate cancer cells would suggest targeting the protein, perhaps through newly developed proteolysis targeting chimera (PROTAC) molecules (89–91). However, there are potential off target effects of m^6^A-sensitive tissues with enzymatic inhibitors, and targeting the AR-METTL3 interaction through small molecule peptide disruption may prove to be a better, PCa-specific strategy. Ultimately, we suggest METTL3, through its interaction with AR, remains an attractive therapeutic target for PCa.

## Acknowledgements

We would like to thank the Prostate Cancer Foundation, and Drs. Howard Soule, PhD and Andrea K. Miyahira PhD. We would also really like to thank the Department of Cancer Biology at Loyola University Chicago and the support of the chair, Dr. Nancy Zeleznik-Le, PhD. We would also like to thank the generous support of the Cardinal Bernadin Cancer Center, and its director William Small Jr., MD, FACRO, FACR, FASTRO. We thank Dr. Peter Nelson, MD of UW Fred Hutchinson Cancer Center for providing LNCaP-APIPC and LNCaP-shAR cells. We thank Nick Achille, Dr. Alvaro Hernandez, PhD of the UIUC Roy J. Carver Biotechnology Center, Carlos Martinez and Dr. Mark Maienschein-Cline, PhD of the Research Informatic Core at UIC for their technical support with ChIP-sequencing. We would finally like to thank Dr. Arul Chinnaiyan, MD/PhD and Dr. Sethuramasundaram Pitchiaya, PhD at the University of Michigan, and Dr. Russell Z. Szmulewitz, MD at the University of Chicago for reagents, important discussions, and guidance. And Dr. Alan Diamond, PhD at University of Illinois at Chicago, for discussions, guidance and critical review of the manuscript.

## Conflicts of Interest

The authors declare no conflicts of interest.

**Supplementary Figure S1. siRNA target sites and complementarity. A)** Schematic of METTL3 sequence with sites targeted by siRNAs and shRNAs indicated. **B)** The siRNA sequence of siMETTL3-1, siMETTL3-2, siMETTL3-3, siMETTL3-4, and siMETTL3-5 and sequences of shMETTL3-1, shMETTL3-2 and shMETTL3-3 are shown labeled as “query’”. All complementary sequences (7 base pairs and up) of in the human (hMETTL3) coding sequence are shown. siMETTL3-4 and siMETTL3-5 target only non-coding regions of hMETTL3. All complementary sequences (7 base pairs and up) of in the full hMETTL3 sequence are shown. The fully identical sequences occur in intron 6 just upstream to the start of exon 7. Blastn off-target search of the guide (antisense) strand sequences of siMETTL3-1, siMETTL3-2, siMETTL3-3, siMETTL3-4, and siMETTL3-5 (RefSeq_RNA database) identified no non-target transcripts with greater than 17 contiguous nucleotides of homology to the 19mer siRNAs. **C)** The 580 amino acid sequence of the canonical hMETTL3 with the target regions of the siRNAs and shRNAs highlighted. The methyl transferase domain, active site DPPW and CCCH Zn finger domains are indicated for reference.

**Supplementary Figure S2. METTL3 siRNA knockdown inhibits proliferation in VCaP, CWR-R1 and CWR-R1 Enz^R^ cell lines. A-C)** Expression of METTL3 mRNA in VCaP, CWR-R1 and CWR-R1 Enz^R^ cell lines quantified by qPCR after transfection of METTL3-targeting siRNAs. Expression is relative to non-silencing control (NSC) and calculated using the ΔΔCt method. **E-G)** Western blots demonstrating knockdown of METTL3 by three siRNAs directed toward METTL3 in VCaP, CWR-R1 and CWR-R1 Enz^R^ cell lines. **H-M)** Cell viability of prostate adenocarcinoma cell lines VCaP, CWR-R1 and CWR-R1 Enz^R^ after transfection of METTL3-targeting or NSC siRNAs over a 7-day time course in the presence or absence of 20 µM enzalutamide Viability was evaluated using the CellTiter-Glo® Luminescent Cell Viability Assay kit (Promega, Madison, WI, USA).

**Supplementary Figure S3. STM2457 efficacy in CWR-R1, CWR-R1 Enz^R^, VCaP and VCaP Enz^R^ cell lines and METTL3 protein levels in LNCaP shNSC cells overexpressing WT or Mutant D395A W398A METTL3 A-B)** Log-dose response curves of METTL3 inhibitor STM2457 with or without enzalutamide (20 µM) in CWR-R1, CWR-R1 Enz^R^, VCaP and VCaP Enz^R^ prostate adenocarcinoma cell lines in the luminescence cell viability assay at day 5. **C-E)** Western blots showing levels of METTL3 in LNCaP shNSC cells and in LCaP shNSC cells overexpressing WT or Mutant D395A W398A METTL3 and treated with or without METTL3 targeting siRNAs. Cells were treated with either siMETTL3-3 which targets a coding region of METTL3 or with METTL3-4 orMETTL3-5 which target non-coding regions of METTL3 present in the overexpression vectors.

**Supplementary Figure S4. METTL3 and AR CoIP: higher exposure A-C)** Higher Exposure of Western blot demonstrating co-immunoprecipitation of AR and METTL3 in LNCaP, LNCaP Enz^R^ and CWR-R1 cell lines using IgG antibodies as a negative control. **Western blot of ChIP lysates from doxycycline-inducible shMETTL3 LNCaP cells. D-E)** ChIP lysates from METTL3 IP, AR IP and control IP (Rabbit IgG IP) in LNCaP cells transduced with doxycycline-inducible shMETTL3-1 or shMETTL3-2 and treated with or without doxycycline and/or R1881 were analyzed by Western blot. Levels of METTL3, AR, were evaluated in the input, immunoprecipitated (IP), and corresponding post-IP (clear) fractions under each condition.

**Supplementary Figure S5. Effect of METTL3 knockdown with or without enzalutamide treatment on expression of target genes. A-C)** Expression of *ALOXE3*, *ILF2*, and *TLE1* mRNA in LNCaP cells stably transduced with doxycycline-inducible shRNA targeting METTL3 and cultured with or without doxycycline (1µg/ml) quantified by qPCR. Cells were also cultured with or without enzalutamide (20 µM). Expression is relative to NSC and calculated using the ΔΔCt method.

**Supplementary Figure S6. ChIP-seq track plots of METTL3 at known AR target genes.**

ChIP-seq track plots of AR and METTL3 binding at AR target genes *KLK3*, *TMPRSS2*, and

*c1orf116 (SARG)* in LNCaP cells expressing two doxycycline-inducible shMETTL3 constructs, **A)** sh-METTL3-1 and **B)** sh-METTL3-2. Peaks were visualized using the Integrative Genomics Viewer (IGV) (https://igv.org/).

**Supplemental Table 1: Primer, RNAi, and Vector sequences.** Supplemental Table **1A**: qPCR Primer sequences. Supplemental Table **1B**: RNAi Sequences, Supplemental Table **1C**: METTL3 Overexpression Vectors. Supplemental Table **1D**: NanoBit Vectors.

**Supplemental Table 2: Full Gene Ontology (GO - Biological Processes) analyses for AR and METTL3 in LNCaP and 22Rv1 cells**. Gene Ontology/EnrichR outputs corresponding to **Figure 6**. For all experiments, ChIP-seq peaks were analyzed using Cistrome-GO to identify genes that had transcription factor binding sites less than 10kb from a TSS and an adjusted regulatory potential (RP) score of >0.01. For the resulting gene lists, GO enrichment was performed using Enrichr. **A-B)** Full Gene Ontology (GO) pathway analysis list and bar plot for AR candidate target genes in LNCaP. **C-D)** Full Gene Ontology (GO) pathway analysis list and bar plot for METTL3 candidate target genes in LNCaP. **E-F)** Full Gene Ontology (GO) pathway analysis list and bar plot for shared AR and METTL3 candidate target genes in LNCaP. **G)** Venn diagram of AR and METTL3 potential target genes in LNCaP. **H-I)** Full Gene Ontology (GO) pathway analysis list and bar plot for AR candidate target genes in 22Rv1. **J-K)** Full Gene Ontology (GO) pathway analysis list and bar plot for METTL3 candidate target genes in 22Rv1. **L-M)** Full Gene Ontology (GO) pathway analysis list and bar plot for shared AR and METTL3 candidate target genes in 22Rv1. **N)** Venn diagram of AR and METTL3 potential target genes in 22Rv1. **O-P)** Full Gene Ontology (GO) pathway analysis list and bar plot for the shared AR and METTL3 candidate target genes shared in both LNCaP and 22Rv1 cell lines. **Q)** Venn diagram of the LNCaP and 22Rv1 candidate gene nexuses. **R-S)** Full Gene Ontology (GO) pathway analysis list and bar plot for the shared AR and METTL3 candidate target genes shared in both LNCaP-shMETTL3-1 and LNCaP-shMETTL3-2 cell lines in the DMSO/EtOH treatment condition.

**Supplemental MEME Motif Output files.**

---

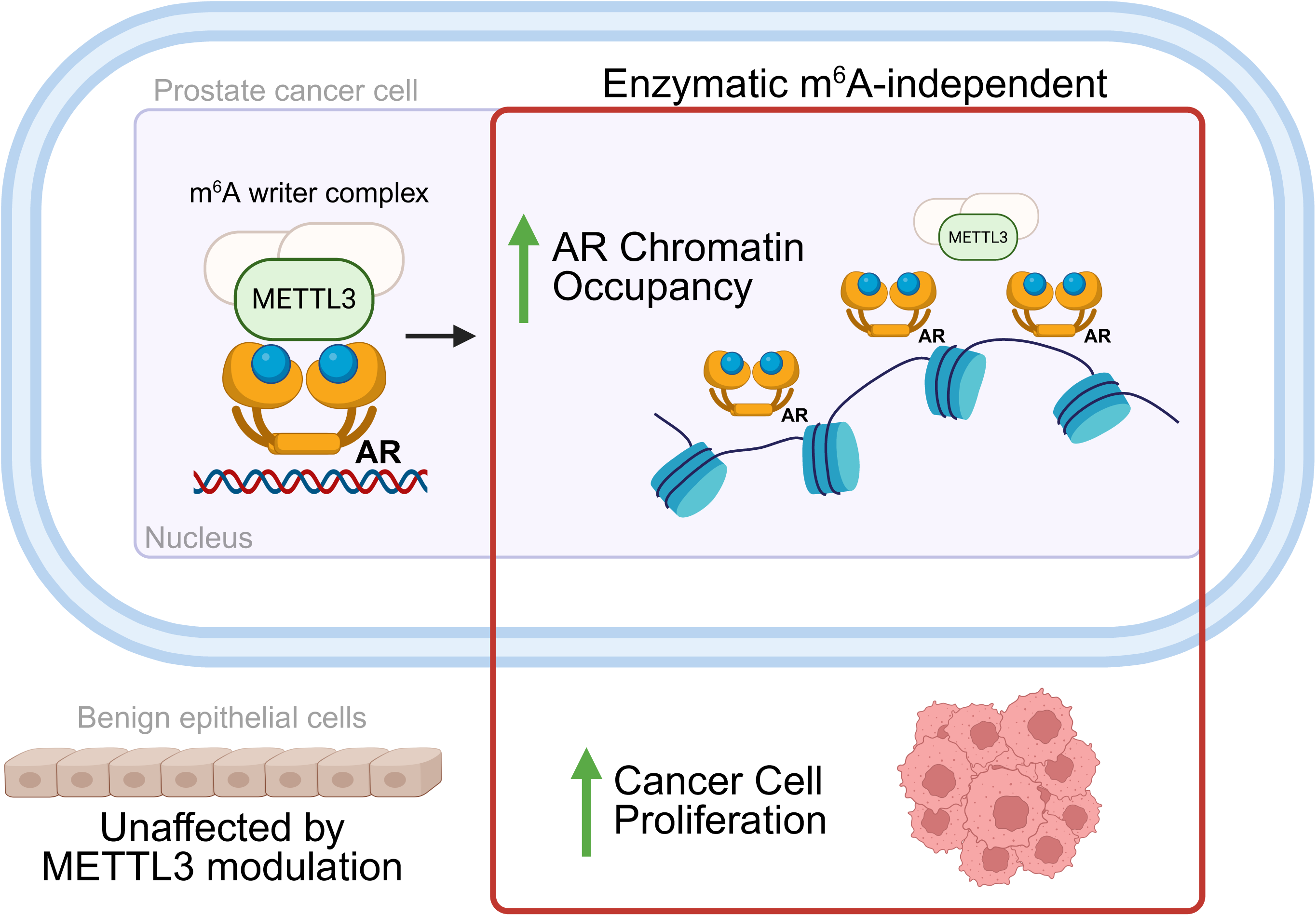

