## Supplemental Files 1-6 and Table 1 and 2 for "A non-enzymatic role for METTL3 as an Androgen Receptor co-regulator that promotes prostate cancer proliferation"

**A**

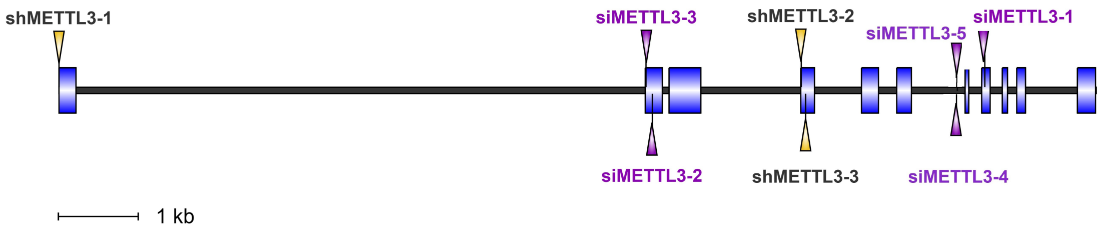


**B**

siMETTL3-1

| Query | 1 | GAACGGGTAGATGAAATTATT | 21 |
|---|---|---|---|
| Complete Coding Sequence | 1348 | GAACGGGTAGATGAAATTATT | 1368 |
| | 1542 | AGATGAAAT | 1550 |
| | 992 | ATGAAATT | 999 |
| | 344 | GGGTAGA | 350 |
| | 1262 | ACGGGTA | 1256 |

shMETTL3-1

| Query | 1 | TCGGACACGTGGAGCTCTA | 19 |
|---|---|---|---|
| Complete Coding Sequence | 4 | TCGGACACGTGGAGCTCTA | 22 |
| | 318 | CGTGGAG | 312 |
| | 746 | AGCTCTA | 740 |
| | 1401 | TCGGACA | 1407 |

shMETTL3-2

| Query | 1 | AGATCCTAGAGCTATTAAA | 19 |
|---|---|---|---|
| Complete Coding Sequence | 734 | AGATCCTAGAGCTATTAAA | 752 |
| | 22 | TAGAGCT | 16 |
| | 222 | AGATCCT | 228 |

siMETTL3-2

| Query | 1 | GATCCTGAGTTAGAGAAGA | 19 |
|---|---|---|---|
| Complete Coding Sequence | 223 | GATCCTGAGTTAGAGAAGA | 241 |
| | 380 | TCCTGAG | 374 |
| | 1593 | TGAGTTA | 1599 |
| | 1727 | TTAGAGA | 1721 |

shMETTL3-3

| Query | 1 | ACAGCCAAGGAACAATCCA | 19 |
|---|---|---|---|
| Complete Coding Sequence | 760 | ACAGCCAAGGAACAATCCA | 778 |
| | 703 | AAGGAACAA | 711 |
| | 626 | CAGCCAAG | 633 |
| | 1438 | AAGGAACA | 1445 |
| | 1500 | ACAATCCA | 1493 |
| | 638 | CAAGGAA | 644 |
| | 839 | CCAAGGA | 845 |

siMETTL3-3

| Query | 1 | GCAGTTCCTGAATTAGCTA | 19 |
|---|---|---|---|
| Complete Coding Sequence | 202 | GCAGTTCCTGAATTAGCTA | 220 |
| | 1213 | GCAGTTCC | 1206 |

siMETTL3-4

| Query | 1 | GCATACAAATAGTTGTTTA | 19 |
|---|---|---|---|
| hMETTL3 GeneID:56339 | 11504 | GCATACAAATAGTTGTTTA | 11522 |
| | 1950 | TAGTTGTTT | 1958 |
| | 3476 | CATACAAAT | 3484 |
| | 3028 | ATACAAAT | 3021 |
| | 5688 | CATACAAA | 5695 |
| | 12476 | TACAAATA | 12483 |
| | 13155 | TACAAATA | 13148 |
| | 13192 | ATACAAAT | 13185 |
| | 933 | AAATAGT | 939 |
| | 980 | ACAAATA | 974 |
| | 2159 | ATACAAA | 2153 |
| | 2390 | GTTGTTT | 2396 |
| | 2397 | GTTGTTT | 2403 |
| | 2604 | ATACAAA | 2598 |
| | 2871 | GCATACA | 2865 |
| | 3838 | ATACAAA | 3832 |
| | 4271 | TAGTTGT | 4277 |
| | 4427 | CATACAA | 4421 |
| | 4439 | CATACAA | 4445 |
| | 4947 | ATACAAA | 4941 |
| | 5420 | TACAAAT | 5414 |
| | 6569 | ATACAAA | 6563 |
| | 6717 | TTGTTTA | 6711 |
| | 7201 | ATACAAA | 7195 |
| | 8943 | ACAAATA | 8937 |
| | 9271 | TAGTTGT | 9265 |
| | 11141 | ATACAAA | 11135 |

siMETTL3-5

| Query | 1 | GAGTCCTAATGCCTGTTTA | 19 |
|---|---|---|---|
| hMETTL3 GeneID:56339 | 11280 | GAGTCCTAATGCCTGTTTA | 11298 |
| | 1192 | GAGTCCTA | 1185 |
| | 2576 | ATGCCTGT | 2569 |
| | 4583 | ATGCCTGT | 4576 |
| | 5163 | TCCTAATG | 5170 |
| | 6843 | GTCCTAAT | 6850 |
| | 206 | GAGTCCT | 200 |
| | 1578 | CCTGTTT | 1584 |
| | 1718 | AGTCCTA | 1712 |
| | 1735 | ATGCCTG | 1741 |
| | 2274 | ATGCCTG | 2280 |
| | 3218 | ATGCCTG | 3224 |
| | 3260 | CCTGTTT | 3254 |
| | 3816 | ATGCCTG | 3822 |
| | 4438 | TGCCTGT | 4432 |
| | 4592 | ATGCCTG | 4598 |
| | 5559 | AATGCCT | 5553 |
| | 6160 | TGCCTGT | 6154 |
| | 7225 | TCCTAAT | 7219 |
| | 7996 | GAGTCCT | 7990 |
| | 8251 | CTGTTTA | 8257 |
| | 8853 | AATGCCT | 8859 |
| | 9998 | CCTGTTT | 9992 |
| | 10845 | TGCCTGT | 10839 |
| | 11105 | TGCCTGT | 11099 |
| | 12338 | GCCTGTT | 12332 |
| | 12747 | CTAATGC | 12741 |

**C**

MSDTWSSIQAHKKQLDSLRERLQRRRKQDSGHLDLRNPEAALSPTFRSDSPVPTAPTSGGPKPSTASAVPELATDPELEKKLLHHLSDLALTLPTDAVSICLAISTPDAPATQDGVESLLQKFA
AQELIEVKRGLLQDDAHPTLVTYADHSKLSAMMGAVAEKKGPGEVAGTVTGQKRRAEQDSTTVAAFASSLVSGLNSSASEPAKEPAKKSRKHAASDVDLEIESLLNQQSTKEQQSKKVSQEIL
ELLNTTTAKEQSIVEKFRSRGRAQVQEFCDYGTKEECMKASDADRPCRKLHFRRIINKHTDESLGDCSFLNTCFHMDTCKYVHYEIDACMDSEAPGSKDHTPSQELALTQSVGGDSSADRL
FPPQWICCDIRYLDVSILGKFAVVMADPPWDIHMELPYGTLTDDEMRRLNIPVLQDDGFLFLWVTGRAMELGRECLNLWGYERVDEIIWVKTNQLQRIIRTGRTGHWLNHGKEHCLVGVKGNP
QGFNQGLDCDVIVAEVRSTSHKPDEIYGMIERLSPGTRKIELFGRPHNVQPNWITLGNQLDGIHLLDPDVVARFKQRYPDGIISKPKNL

Methyl Transferase Domain
CCCH Zn Finger Domains
DPPW Active site DPPW

siMETTL3-1
siMETTL3-2
siMETTL3-3

SDTWSS shMETTL3-1
ILELL shMETTL3-2
TAKEQS shMETTL3-3

Kostlan, et. al (Figure S1)

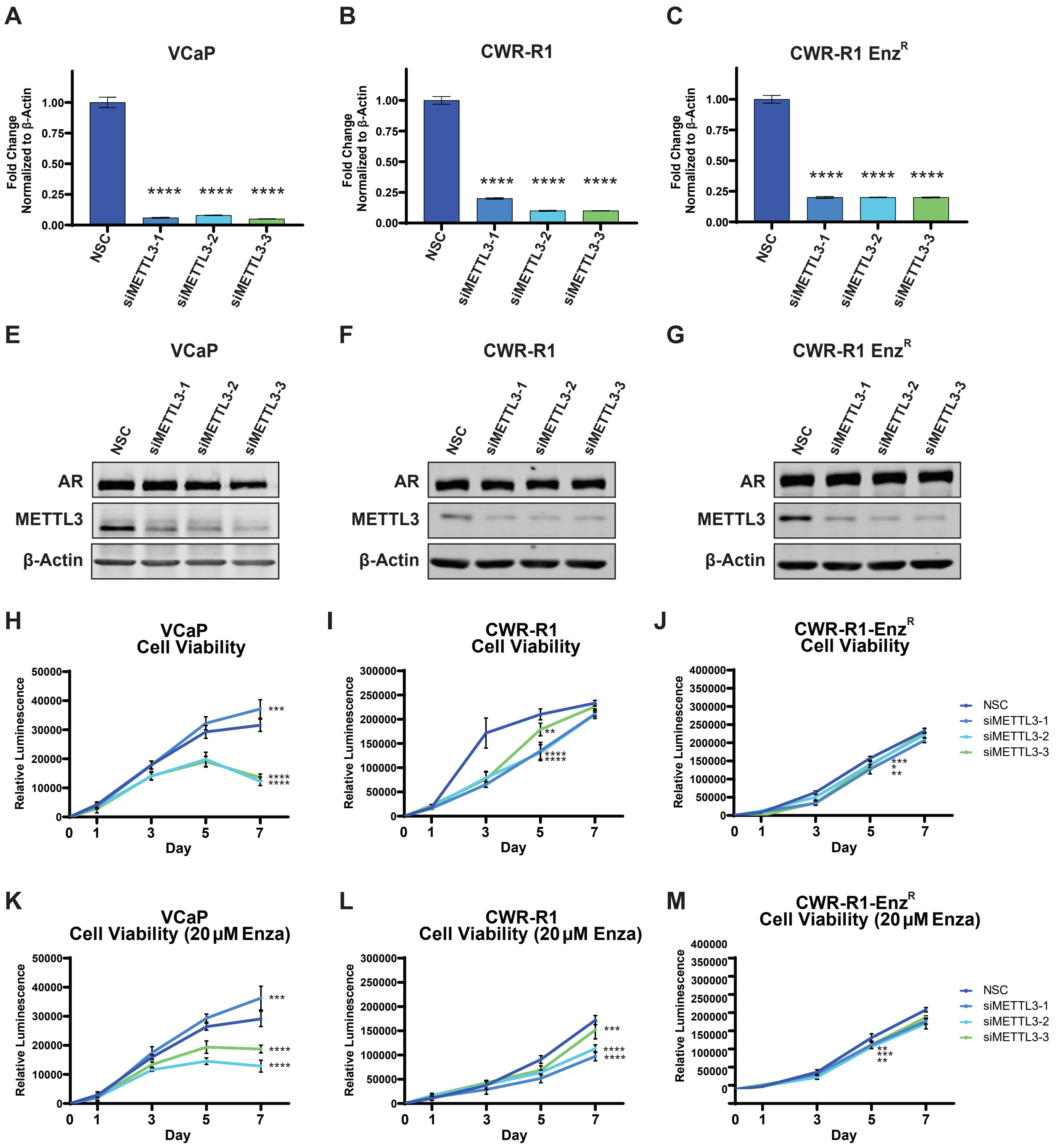


Kostlan, et. al (Figure S2)

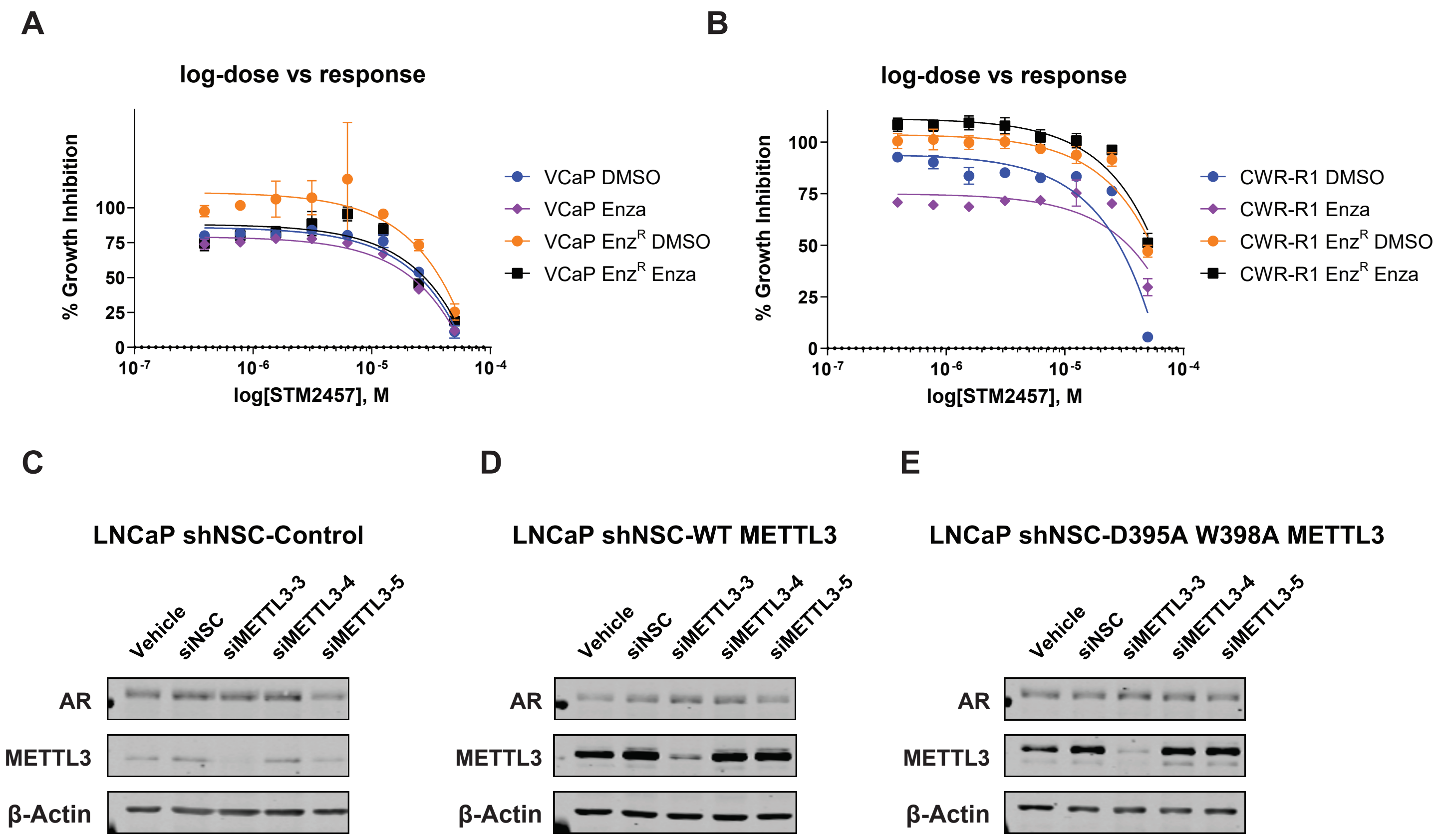
A
log-dose vs response
% Growth Inhibition
log[STM2457], M
VCaP DMSO
VCaP Enza
VCaP Enz^R DMSO
VCaP Enz^R Enza
B
log-dose vs response
% Growth Inhibition
log[STM2457], M
CWR-R1 DMSO
CWR-R1 Enza
CWR-R1 Enz^R DMSO
CWR-R1 Enz^R Enza
C
LNCaP shNSC-Control
D
LNCaP shNSC-WT METTL3
E
LNCaP shNSC-D395A W398A METTL3
Vehicle
siNSC
siMETTL3-3
siMETTL3-4
siMETTL3-5
AR
METTL3
β-Actin

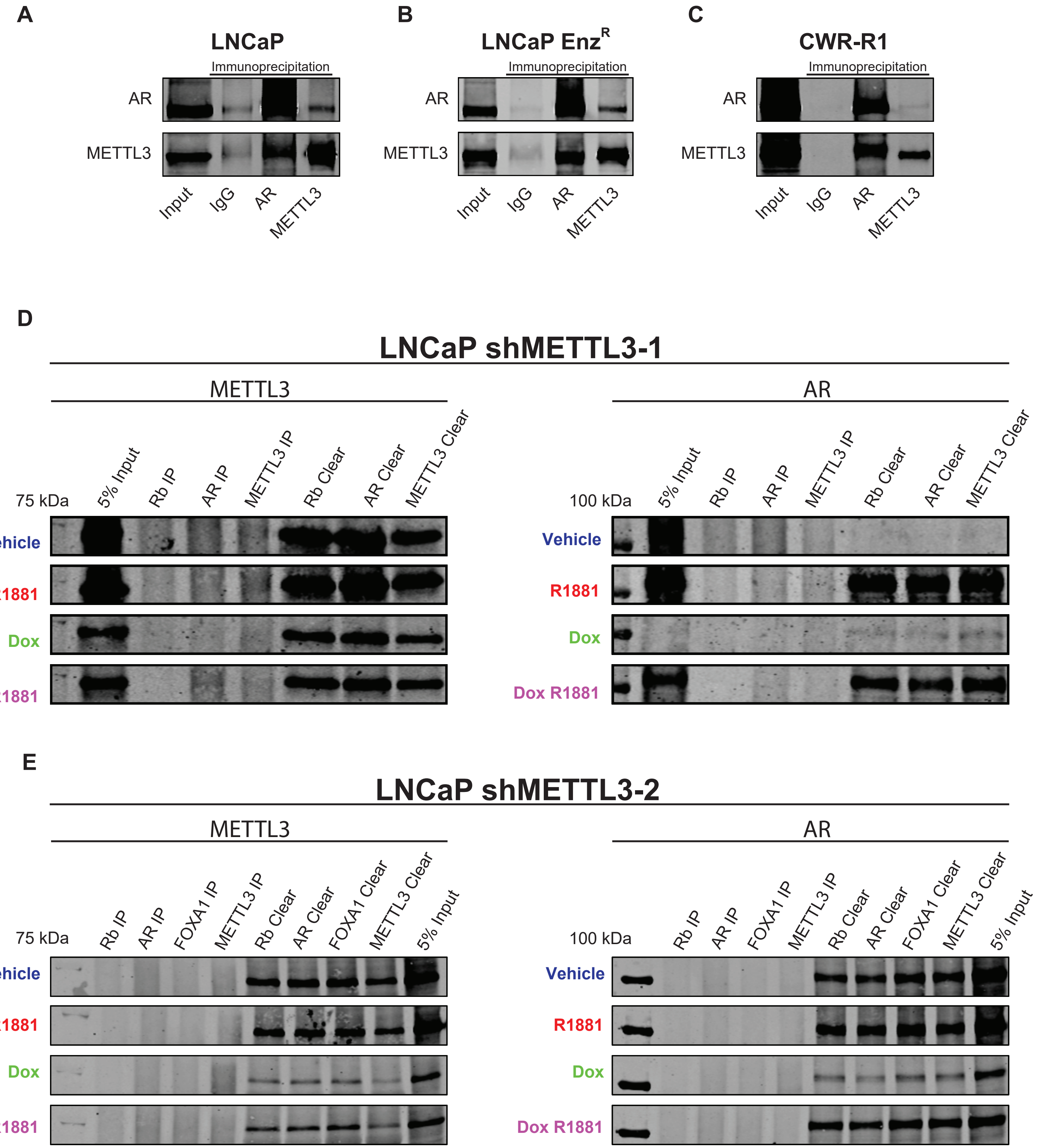


Kostlan, et. al (Figure S4)

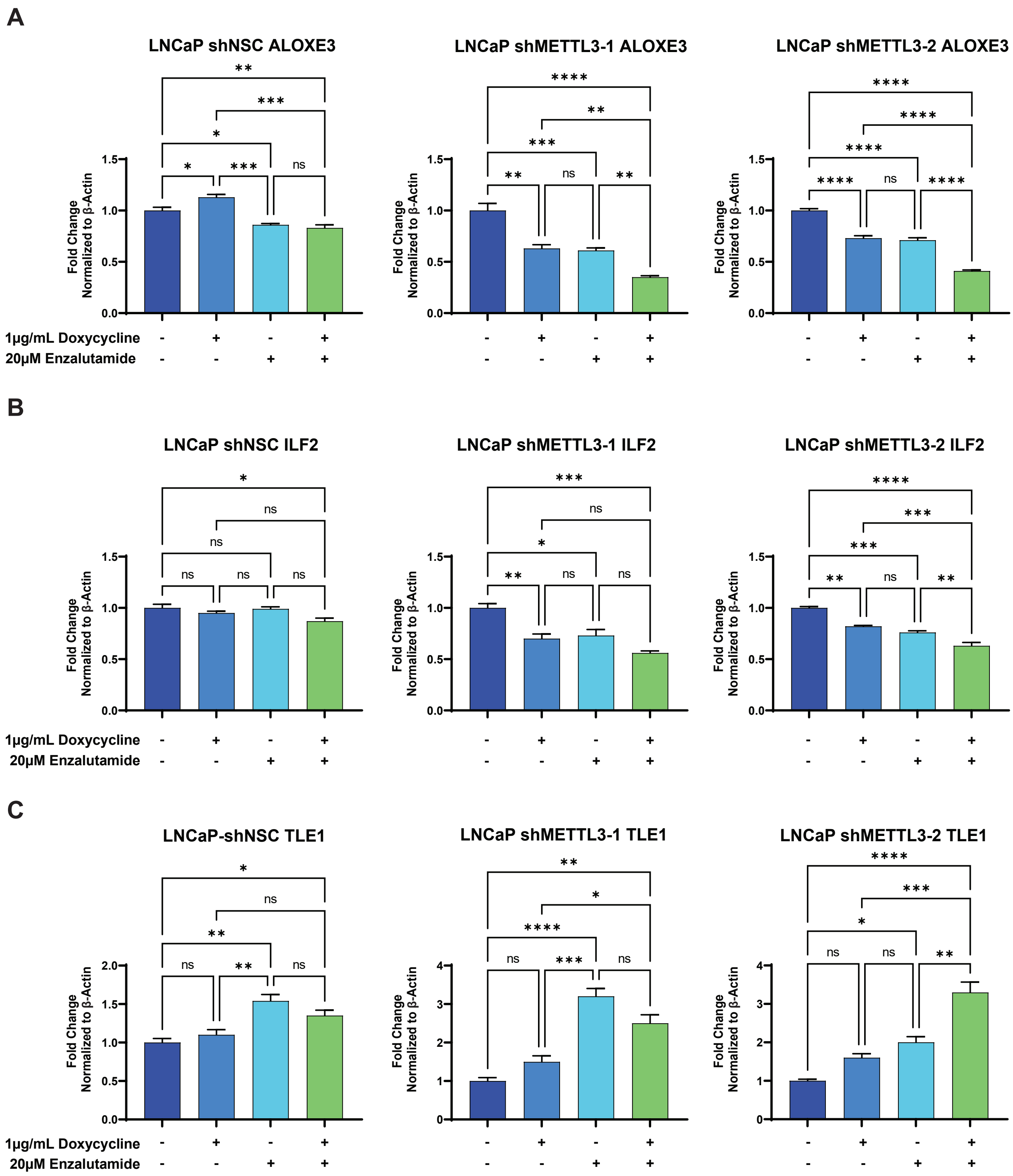
A
LNCaP shNSC ALOXE3
LNCaP shMETTL3-1 ALOXE3
LNCaP shMETTL3-2 ALOXE3
Fold Change Normalized to β-Actin
1µg/mL Doxycycline
20µM Enzalutamide
B
LNCaP shNSC ILF2
LNCaP shMETTL3-1 ILF2
LNCaP shMETTL3-2 ILF2
Fold Change Normalized to β-Actin
1µg/mL Doxycycline
20µM Enzalutamide
C
LNCaP-shNSC TLE1
LNCaP shMETTL3-1 TLE1
LNCaP shMETTL3-2 TLE1
Fold Change Normalized to β-Actin
1µg/mL Doxycycline
20µM Enzalutamide

A

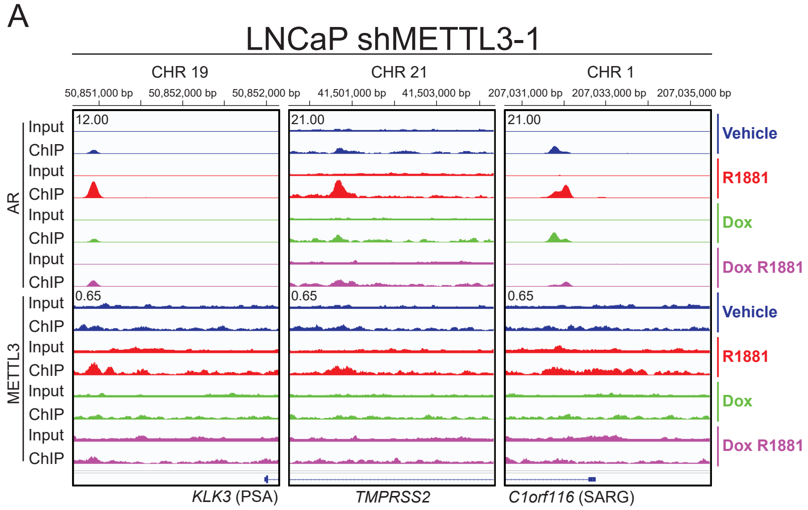
LNCaP shMETTL3-1
CHR 19
CHR 21
CHR 1
50,851,000 bp
50,852,000 bp
50,852,000 bp
41,501,000 bp
41,503,000 bp
207,031,000 bp
207,033,000 bp
207,035,000 bp
AR
METTL3
Input
ChIP
12.00
21.00
21.00
0.65
0.65
0.65
Vehicle
R1881
Dox
Dox R1881
KLK3 (PSA)
TMPRSS2
C1orf116 (SARG)

B

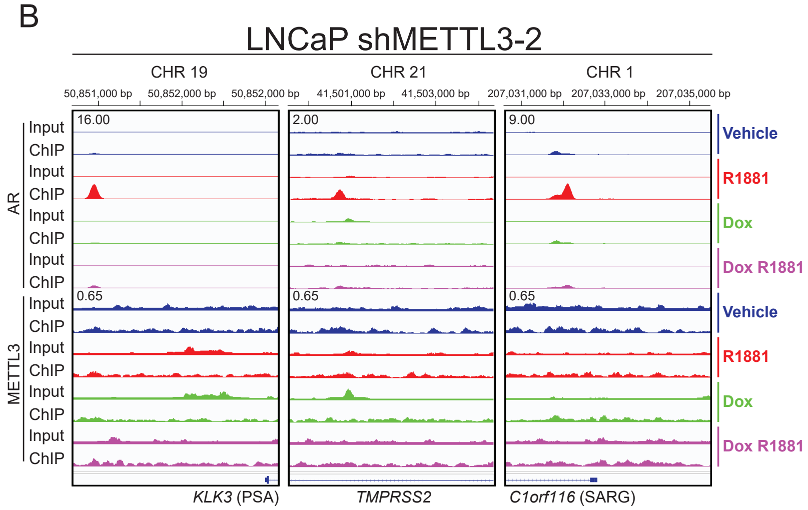
LNCaP shMETTL3-2
CHR 19
CHR 21
CHR 1
50,851,000 bp
50,852,000 bp
50,852,000 bp
41,501,000 bp
41,503,000 bp
207,031,000 bp
207,033,000 bp
207,035,000 bp
AR
METTL3
Input
ChIP
16.00
2.00
9.00
0.65
0.65
0.65
Vehicle
R1881
Dox
Dox R1881
KLK3 (PSA)
TMPRSS2
C1orf116 (SARG)

**Supplemental Table 1:**

**Supplemental Table 1A: qPCR Primers**

| qPCR Primers (Source: IDT) | | |
|---|---|---|
| Gene | Forward Primer | Reverse Primer |
| *AR* | 5′-CGGAAGCTGAAGAAACTTGG-3′ | 5′-ATGGCTTCCAGGACATTCAG-3′ |
| *METTL3* | 5′-TGGGAAGGAACACTGCTTGGTTG-3′ | 5′-AATCCAGACCCTGGTTGAAGCC-3′ |
| *ALOXE3* | 5′-GGTCCTGGAGCGGAAAGAAT-3′ | 5′-GGAGCCTGGGTTCCACTCAT-3′ |
| *TLE1* | 5′-GGCGCAGTATCACAGCCTTA-3′ | 5′-CAACTGCTGCTGCTGTTTGT-3′ |
| *ILF2* | 5′-GTTTTAAGGCGCCATGAGGG-3′ | 5′-GGGAAAGGCCATTTCACACAAAT-3′ |
| *ACTB* | 5′-CACCATTGGCAATGAGCGGTT-3′ | 5′-AGGTCTTTGCGGATGTCCACGT-3′ |

**Supplemental Table 1B: iRNA Sequences**

| iRNA Sequences | | | | |
|---|---|---|---|---|
| | Sense Sequence | Position | Catalog # | Source |
| *NSC* | Silencer™ Select Negative Control No. 1 siRNA | N/A | 43-908-43 | Invitrogen |
| *siMETTL3-1* | 5′- GAACGGGUAGAUGAAAUUAtt -3′ | Exon 8 | s32141 | Invitrogen |
| *siMETTL3-2* | 5′- GAUCCUGAGUUAGAGAAGAtt -3′ | Exon 2 | s32142 | Invitrogen |
| *siMETTL3-3* | 5′- GCAGUUCCUGAAUUAGCUAtt -3′ | Exon 2 | s32143 | Invitrogen |
| *siMETTL3-4* | 5′- GCAUACAAAUAGUUGUUUAtt -3′ | Intron 6 | n322814 | Invitrogen |
| *siMETTL3-5* | 5′- GAGUCCUAAUGCCUGUUUAtt -3′ | Intron 6 | n322813 | Invitrogen |
| *shNSC* | Tet-pLKO-puro-shScrambled | N/A | 47541 | Addgene (Charles Rudin) |
| *shMETTL3-1* | 5′- TCGGACACGTGGAGCTCTA-3′ | Exon 1 | V3SH11252-229451971 | Horizon Discovery |
| *shMETTL3-2* | 5′- AGATCCTAGAGCTATTAAA -3′ | Exon 4 | V3SH11252-229373728 | Horizon Discovery |
| *shMETTL3-3* | 5′- ACAGCCAAGGAACAATCCA -3′ | Exon 4 | V3SH11252-228569617 | Horizon Discovery |

**Supplemental Table 1C: METTL3 Overexpression Vectors**

| METTL3 Overexpression Vector | | Catalog # | Source |
|---|---|---|---|
| METTL3 WT | pLV[Exp]-Bsd-CMV>3xFLAG/hMETTL3[NM_019852.5] | VB251009-1391vwr | Vector Builder |
| METTL3 D395A | pLV[Exp]-Bsd-CMV>3xFLAG /{hMETTL3[NM_019852.5](D395A)} | VB251009-1508sru | Vector Builder |
| METTL3 D395A,W398A | pLV[Exp]-Bsd-CMV>3xFLAG /{hMETTL3[NM_019852.5](D395A,W398A)} | VB251009-1513wzv | Vector Builder |
| Control Vector | GFP control | eGFP-LV105 | GeneCopoeia |

**Supplemental Table 1D: NanoBiT® Vectors**

| NanoBiT® Vectors (GeneScript Biotech (Piscataway, NJ, USA) | |
|---|---|
| METTL3-lg-N cDNA (cloned into pcDNA3.1(+) backbone) | ATGGTCTTCACACTCGAAGATTTCGTTGGGGACTGGGAACAGACAGCCGCCTACAA<br>CCTGGACCAAGTCCTTGAACAGGGAGGTGTGTCCAGTTTGCTGCAGAATCTCGCCG<br>TGTCCGTAACTCCGATCCAAAGGATTGTCCGGAGCGGTGAAAATGCCCTGAAGATC<br>GACATCCATGTCATCATCCCGTATGAAGGTCTGAGCGCCGACCAAATGGCCCAGAT<br>CGAAGAGGTGTTTAAGGTGGTGTACCCTGTGGATGATCATCACTTTAAGGTGATCC<br>TGCCCTATGGCACACTGGTAATCGACGGGGTTACGCCGAACATGCTGAACTATTTC<br>GGACGGCCGTATGAAGGCATCGCCGTGTTCGACGGCAAAAAGATCACTGTAACAGG<br>GACCCTGTGGAACGGCAACAAAATTATCGACGAGCGCCTGATCACCCCCGACGGCT<br>CCATGCTGTTCCGAGTAACCATCAACAGCTCTGGCTCGAGCGGTGGTGGCGGGAGC<br>GGAGGTGGAGGGTCGTCAGGTACCTCGGACACGTGGAGCTCTATCCAGGCCCACAA<br>GAAGCAGCTGGACTCTCTGCGGGAGAGGCTGCAGCGGAGGCGGAAGCAGGACTCGG<br>GGCACTTGGATCTACGGAATCCAGAGGCAGCATTGTCTCCAACCTTCCGTAGTGAC<br>AGCCCAGTGCCTACTGCACCCACCTCTGGTGGCCCTAAGCCCAGCACAGCTTCAGC<br>AGTTCCTGAATTAGCTACAGATCCTGAGTTAGAGAAGAAGTTGCTACACCACCTCT<br>CTGATCTGGCCTTAACATTGCCCACTGATGCTGTGTCCATCTGTCTTGCCATCTCC<br>ACGCCAGATGCTCCTGCCACTCAAGATGGGGTAGAAAGCCTCCTGCAGAAGTTTGC<br>AGCTCAGGAGTTGATTGAGGTAAAGCGAGGTCTCCTACAAGATGATGCACATCCTA<br>CTCTTGTAACCTATGCTGACCATTCCAAGCTCTCTGCCATGATGGGTGCTGTGGCA<br>GAAAAGAAGGGCCCTGGGGAGGTAGCAGGGACTGTCACAGGGCAGAAGCGGCGTGC<br>AGAACAGGACTCGACTACAGTAGCTGCCTTTGCCAGTTCGTTAGTCTCTGGTCTGA<br>ACTCTTCAGCATCGGAACCAGCAAAGGAGCCAGCCAAGAAATCAAGGAAACATGCT<br>GCCTCAGATGTTGATCTGGAGATAGAGAGCCTTCTGAACCAACAGTCCACTAAGGA<br>ACAACAGAGCAAGAAGGTCAGTCAGGAGATCCTAGAGCTATTAAATACTACAACAG<br>CCAAGGAACAATCCATTGTTGAAAAATTTCGCTCTCGAGGTCGGGCCCAAGTGCAA<br>GAATTCTGTGACTATGGAACCAAGGAGGAGTGCATGAAAGCCAGTGATGCTGATCG<br>ACCCTGTCGCAAGCTGCACTTCAGACGAATTATCAATAAACACACTGATGAGTCTT<br>TAGGTGACTGCTCTTTCCTTAATACATGTTTCCACATGGATACCTGCAAGTATGTT<br>CACTATGAAATTGATGCTTGCATGGATTCTGAGGCCCCTGGCAGCAAAGACCACAC<br>GCCAAGCCAGGAGCTTGCTCTTACACAGAGTGTCGGAGGTGATTCCAGTGCAGACC<br>GACTCTTCCCACCTCAGTGGATCTGTTGTGATATCCGCTACCTGGACGTCAGTATC<br>TTGGGCAAGTTTGCAGTTGTGATGGCTGACCCACCCTGGGATATTCACATGGAACT<br>GCCCTATGGGACCCTGACAGATGATGAGATGCGCAGGCTCAACATACCCGTACTAC<br>AGGATGATGGCTTTCTCTTCCTCTGGGTCACAGGCAGGGCCATGGAGTTGGGGAGA<br>GAATGTCTAAACCTCTGGGGGTATGAACGGGTAGATGAAATTATTTGGGTGAAGAC<br>AAATCAACTGCAACGCATCATTCGGACAGGCCGTACAGGTCACTGGTTGAACCATG<br>GGAAGGAACACTGCTTGGTTGGTGTCAAAGGAAATCCCCAAGGCTTCAACCAGGGT<br>CTGGATTGTGATGTGATCGTAGCTGAGGTTCGTTCCACCAGTCATAAACCAGATGA<br>AATCTATGGCATGATTGAAAGACTATCTCCTGGCACTCGCAAGATTGAGTTATTTG<br>GACGACCACACAATGTGCAACCCAACTGGATCACCCTTGGAAACCAACTGGATGGG<br>ATCCACCTACTAGACCCAGATGTGGTTGCACGGTTCAAGCAAAGGTACCCAGATGG<br>TATCATCTCTAAACCTAAGAATTTATAG |
| METTL3-lg-C cDNA (cloned into pcDNA3.1(+) backbone) | ATGTCGGACACGTGGAGCTCTATCCAGGCCCACAAGAAGCAGCTGGACTCTCTGCG<br>GGAGAGGCTGCAGCGGAGGCGGAAGCAGGACTCGGGGCACTTGGATCTACGGAATC<br>CAGAGGCAGCATTGTCTCCAACCTTCCGTAGTGACAGCCCAGTGCCTACTGCACCC<br>ACCTCTGGTGGCCCTAAGCCCAGCACAGCTTCAGCAGTTCCTGAATTAGCTACAGA<br>TCCTGAGTTAGAGAAGAAGTTGCTACACCACCTCTCTGATCTGGCCTTAACATTGC<br>CCACTGATGCTGTGTCCATCTGTCTTGCCATCTCCACGCCAGATGCTCCTGCCACT<br>CAAGATGGGGTAGAAAGCCTCCTGCAGAAGTTTGCAGCTCAGGAGTTGATTGAGGT<br>AAAGCGAGGTCTCCTACAAGATGATGCACATCCTACTCTTGTAACCTATGCTGACC<br>ATTCCAAGCTCTCTGCCATGATGGGTGCTGTGGCAGAAAAGAAGGGCCCTGGGGAG<br>GTAGCAGGGACTGTCACAGGGCAGAAGCGGCGTGCAGAACAGGACTCGACTACAGT<br>AGCTGCCTTTGCCAGTTCGTTAGTCTCTGGTCTGAACTCTTCAGCATCGGAACCAG<br>CAAAGGAGCCAGCCAAGAAATCAAGGAAACATGCTGCCTCAGATGTTGATCTGGAG<br>ATAGAGAGCCTTCTGAACCAACAGTCCACTAAGGAACAACAGAGCAAGAAGGTCAG<br>TCAGGAGATCCTAGAGCTATTAAATACTACAACAGCCAAGGAACAATCCATTGTTG |

| | |
|---|---|
| | AAAAATTTCGCTCTCGAGGTCGGGCCCAAGTGCAAGAATTCTGTGACTATGGAACC<br>AAGGAGGAGTGCATGAAAGCCAGTGATGCTGATCGACCCTGTCGCAAGCTGCACTT<br>CAGACGAATTATCAATAAACACACTGATGAGTCTTTAGGTGACTGCTCTTTCCTTA<br>ATACATGTTTCCACATGGATACCTGCAAGTATGTTCACTATGAAATTGATGCTTGC<br>ATGGATTCTGAGGCCCCTGGCAGCAAAGACCACACGCCAAGCCAGGAGCTTGCTCT<br>TACACAGAGTGTCGGAGGTGATTCCAGTGCAGACCGACTCTTCCCACCTCAGTGGA<br>TCTGTTGTGATATCCGCTACCTGGACGTCAGTATCTTGGGCAAGTTTGCAGTTGTG<br>ATGGCTGACCCACCCTGGGATATTCACATGGAACTGCCCTATGGGACCCTGACAGA<br>TGATGAGATGCGCAGGCTCAACATACCCGTACTACAGGATGATGGCTTTCTCTTCC<br>TCTGGGTCACAGGCAGGGCCATGGAGTTGGGGAGAGAATGTCTAAACCTCTGGGGG<br>TATGAACGGGTAGATGAAATTATTTGGGTGAAGACAAATCAACTGCAACGCATCAT<br>TCGGACAGGCCGTACAGGTCACTGGTTGAACCATGGGAAGGAACACTGCTTGGTTG<br>GTGTCAAAGGAAATCCCCAAGGCTTCAACCAGGGTCTGGATTGTGATGTGATCGTA<br>GCTGAGGTTCGTTCCACCAGTCATAAACCAGATGAAATCTATGGCATGATTGAAAG<br>ACTATCTCCTGGCACTCGCAAGATTGAGTTATTTGGACGACCACACAATGTGCAAC<br>CCAACTGGATCACCCTTGGAAACCAACTGGATGGGATCCACCTACTAGACCCAGAT<br>GTGGTTGCACGGTTCAAGCAAAGGTACCCAGATGGTATCATCTCTAAACCTAAGAA<br>TTTAGGCTCGAGCGGTGGTGGCGGGAGCGGAGGTGGAGGGTCGTCAGGTGTCTTCA<br>CACTCGAAGATTTCGTTGGGGACTGGGAACAGACAGCCGCCTACAACCTGGACCAA<br>GTCCTTGAACAGGGAGGTGTGTCCAGTTTGCTGCAGAATCTCGCCGTGTCCGTAAC<br>TCCGATCCAAAGGATTGTCCGGAGCGGTGAAAATGCCCTGAAGATCGACATCCATG<br>TCATCATCCCGTATGAAGGTCTGAGCGCCGACCAAATGGCCCAGATCGAAGAGGTG<br>TTTAAGGTGGTGTACCCTGTGGATGATCATCACTTTAAGGTGATCCTGCCCTATGG<br>CACACTGGTAATCGACGGGGTTACGCCGAACATGCTGAACTATTTCGGACGGCCGT<br>ATGAAGGCATCGCCGTGTTCGACGGCAAAAAGATCACTGTAACAGGGACCCTGTGG<br>AACGGCAACAAAATTATCGACGAGCGCCTGATCACCCCCGACGGCTCCATGCTGTT<br>CCGAGTAACCATCAACAGCTAA |
| AR-smN cDNA (cloned into pcDNA3.1(+) backbone) | ATGGTGACCGGCTACCGGCTGTTCGAGGAGATTCTGTCTGGCTCGAGCGGTGGTGG<br>CGGGAGCGGAGGTGGAGGGTCGTCAGGTACCGAAGTGCAGTTAGGGCTGGGAAGGG<br>TCTACCCTCGGCCGCCGTCCAAGACCTACCGAGGAGCTTTCCAGAATCTGTTCCAG<br>AGCGTGCGCGAAGTGATCCAGAACCCGGGCCCCAGGCACCCAGAGGCCGCGAGCGC<br>AGCACCTCCCGGCGCCAGTTTGCTGCTGCTGCAGCAGCAGCAGCAGCAGCAGCAGC<br>AGCAGCAGCAGCAGCAGCAGCAGCAGCAGCAGCAGCAGCAGCAAGAGACTAGCCCC<br>AGGCAGCAGCAGCAGCAGCAGGGTGAGGATGGTTCTCCCCAAGCCCATCGTAGAGG<br>CCCCACAGGCTACCTGGTCCTGGATGAGGAACAGCAACCTTCACAGCCGCAGTCGG<br>CCCTGGAGTGCCACCCCGAGAGAGGTTGCGTCCCAGAGCCTGGAGCCGCCGTGGCC<br>GCCAGCAAGGGGCTGCCGCAGCAGCTGCCAGCACCTCCGGACGAGGATGACTCAGC<br>TGCCCCATCCACGTTGTCCCTGCTGGGCCCCACTTTCCCCGGCTTAAGCAGCTGCT<br>CCGCTGACCTTAAAGACATCCTGAGCGAGGCCAGCACCATGCAACTCCTTCAGCAA<br>CAGCAGCAGGAAGCAGTATCCGAAGGCAGCAGCAGCGGGAGAGCGAGGGAGGCCTC<br>GGGGGCTCCCACTTCCTCCAAGGACAATTACTTAGGGGGCACTTCGACCATTTCTG<br>ACAACGCCAAGGAGTTGTGTAAGGCAGTGTCGGTGTCCATGGGCCTGGGTGTGGAG<br>GCGTTGGAGCATCTGAGTCCAGGGGAACAGCTTCGGGGGGATTGCATGTACGCCCC<br>ACTTTTGGGAGTTCCACCCGCTGTGCGTCCCACTCCTTGTGCCCCATTGGCCGAAT<br>GCAAAGGTTCTCTGCTAGACGACAGCGCAGGCAAGAGCACTGAAGATACTGCTGAG<br>TATTCCCCTTTCAAGGGAGGTTACACCAAAGGGCTAGAAGGCGAGAGCCTAGGCTG<br>CTCTGGCAGCGCTGCAGCAGGGAGCTCCGGGACACTTGAACTGCCGTCTACCCTGT<br>CTCTCTACAAGTCCGGAGCACTGGACGAGGCAGCTGCGTACCAGAGTCGCGACTAC<br>TACAACTTTCCACTGGCTCTGGCCGGACCGCCGCCCCCTCCGCCGCCTCCCCATCC<br>CCACGCTCGCATCAAGCTGGAGAACCCGCTGGACTACGGCAGCGCCTGGGCGGCTG<br>CGGCGGCGCAGTGCCGCTATGGGGACCTGGCGAGCCTGCATGGCGCGGGTGCAGCG<br>GGACCCGGTTCTGGGTCACCCTCAGCCGCCGCTTCCTCATCCTGGCACACTCTCTT<br>CACAGCCGAAGAAGGCCAGTTGTATGGACCGTGTGGTGGTGGTGGGGGTGGTGGCG<br>GCGGCGGCGGCGGCGGCGGCGGCGGCGGCGGCGGCGGCGGCGGCGGCGAGGCGGGA<br>GCTGTAGCCCCCTACGGCTACACTCGGCCCCCTCAGGGGCTGGCGGGCCAGGAAAG<br>CGACTTCACCGCACCTGATGTGTGGTACCCTGGCGGCATGGTGAGCAGAGTGCCCT<br>ATCCCAGTCCCACTTGTGTCAAAAGCGAAATGGGCCCCTGGATGGATAGCTACTCC |

| | |
|---|---|
| | GGACCTTACGGGGACATGCGTTTGGAGACTGCCAGGGACCATGTTTTGCCCATTGA<br>CTATTACTTTCCACCCCAGAAGACCTGCCTGATCTGTGGAGATGAAGCTTCTGGGT<br>GTCACTATGGAGCTCTCACATGTGGAAGCTGCAAGGTCTTCTTCAAAAGAGCCGCT<br>GAAGGGAAACAGAAGTACCTGTGCGCCAGCAGAAATGATTGCACTATTGATAAATT<br>CCGAAGGAAAAATTGTCCATCTTGTCGTCTTCGGAAATGTTATGAAGCAGGGATGA<br>CTCTGGGAGCCCGGAAGCTGAAGAAACTTGGTAATCTGAAACTACAGGAGGAAGGA<br>GAGGCTTCCAGCACCACCAGCCCCACTGAGGAGACAACCCAGAAGCTGACAGTGTC<br>ACACATTGAAGGCTATGAATGTCAGCCCATCTTTCTGAATGTCCTGGAAGCCATTG<br>AGCCAGGTGTAGTGTGTGCTGGACACGACAACAACCAGCCCGACTCCTTTGCAGCC<br>TTGCTCTCTAGCCTCAATGAACTGGGAGAGAGACAGCTTGTACACGTGGTCAAGTG<br>GGCCAAGGCCTTGCCTGGCTTCCGCAACTTACACGTGGACGACCAGATGGCTGTCA<br>TTCAGTACTCCTGGATGGGGCTCATGGTGTTTGCCATGGGCTGGCGATCCTTCACC<br>AATGTCAACTCCAGGATGCTCTACTTCGCCCCTGATCTGGTTTTCAATGAGTACCG<br>CATGCACAAGTCCCGGATGTACAGCCAGTGTGTCCGAATGAGGCACCTCTCTCAAG<br>AGTTTGGATGGCTCCAAATCACCCCCCAGGAATTCCTGTGCATGAAAGCACTGCTA<br>CTCTTCAGCATTATTCCAGTGGATGGGCTGAAAAATCAAAAATTCTTTGATGAACT<br>TCGAATGAACTACATCAAGGAACTCGATCGTATCATTGCATGCAAAAGAAAAAATC<br>CCACATCCTGCTCAAGACGCTTCTACCAGCTCACCAAGCTCCTGGACTCCGTGCAG<br>CCTATTGCGAGAGAGCTGCATCAGTTCACTTTTGACCTGCTAATCAAGTCACACAT<br>GGTGAGCGTGGACTTTCCGGAAATGATGGCAGAGATCATCTCTGTGCAAGTGCCCA<br>AGATCCTTTCTGGGAAAGTCAAGCCCATCTATTTCCACACCCAGTGA |
| AR-smC cDNA (cloned into pcDNA3.1(+) backbone) | ATGGAAGTGCAGTTAGGGCTGGGAAGGGTCTACCCTCGGCCGCCGTCCAAGACCTA<br>CCGAGGAGCTTTCCAGAATCTGTTCCAGAGCGTGCGCGAAGTGATCCAGAACCCGG<br>GCCCCAGGCACCCAGAGGCCGCGAGCGCAGCACCTCCCGGCGCCAGTTTGCTGCTG<br>CTGCAGCAGCAGCAGCAGCAGCAGCAGCAGCAGCAGCAGCAGCAGCAGCAGCAGCA<br>GCAGCAGCAGCAGCAAGAGACTAGCCCCAGGCAGCAGCAGCAGCAGCAGGGTGAGG<br>ATGGTTCTCCCCAAGCCCATCGTAGAGGCCCCACAGGCTACCTGGTCCTGGATGAG<br>GAACAGCAACCTTCACAGCCGCAGTCGGCCCTGGAGTGCCACCCCGAGAGAGGTTG<br>CGTCCCAGAGCCTGGAGCCGCCGTGGCCGCCAGCAAGGGGCTGCCGCAGCAGCTGC<br>CAGCACCTCCGGACGAGGATGACTCAGCTGCCCCATCCACGTTGTCCCTGCTGGGC<br>CCCACTTTCCCCGGCTTAAGCAGCTGCTCCGCTGACCTTAAAGACATCCTGAGCGA<br>GGCCAGCACCATGCAACTCCTTCAGCAACAGCAGCAGGAAGCAGTATCCGAAGGCA<br>GCAGCAGCGGGAGAGCGAGGGAGGCCTCGGGGGCTCCCACTTCCTCCAAGGACAAT<br>TACTTAGGGGGCACTTCGACCATTTCTGACAACGCCAAGGAGTTGTGTAAGGCAGT<br>GTCGGTGTCCATGGGCCTGGGTGTGGAGGCGTTGGAGCATCTGAGTCCAGGGGAAC<br>AGCTTCGGGGGGATTGCATGTACGCCCCACTTTTGGGAGTTCCACCCGCTGTGCGT<br>CCCACTCCTTGTGCCCCATTGGCCGAATGCAAAGGTTCTCTGCTAGACGACAGCGC<br>AGGCAAGAGCACTGAAGATACTGCTGAGTATTCCCCTTTCAAGGGAGGTTACACCA<br>AAGGGCTAGAAGGCGAGAGCCTAGGCTGCTCTGGCAGCGCTGCAGCAGGGAGCTCC<br>GGGACACTTGAACTGCCGTCTACCCTGTCTCTCTACAAGTCCGGAGCACTGGACGA<br>GGCAGCTGCGTACCAGAGTCGCGACTACTACAACTTTCCACTGGCTCTGGCCGGAC<br>CGCCGCCCCCTCCGCCGCCTCCCCATCCCCACGCTCGCATCAAGCTGGAGAACCCG<br>CTGGACTACGGCAGCGCCTGGGCGGCTGCGGCGGCGCAGTGCCGCTATGGGGACCT<br>GGCGAGCCTGCATGGCGCGGGTGCAGCGGGACCCGGTTCTGGGTCACCCTCAGCCG<br>CCGCTTCCTCATCCTGGCACACTCTCTTCACAGCCGAAGAAGGCCAGTTGTATGGA<br>CCGTGTGGTGGTGGTGGGGGTGGTGGCGGCGGCGGCGGCGGCGGCGGCGGCGGCGG<br>CGGCGGCGGCGGCGGCGGCGAGGCGGGAGCTGTAGCCCCCTACGGCTACACTCGGC<br>CCCCTCAGGGGCTGGCGGGCCAGGAAAGCGACTTCACCGCACCTGATGTGTGGTAC<br>CCTGGCGGCATGGTGAGCAGAGTGCCCTATCCCAGTCCCACTTGTGTCAAAAGCGA<br>AATGGCCCCTGGATGGATAGCTACTCCGGACCTTACGGGGACATGCGTTTGGAGA<br>CTGCCAGGGACCATGTTTTGCCCATTGACTATTACTTTCCACCCCAGAAGACCTGC<br>CTGATCTGTGGAGATGAAGCTTCTGGGTGTCACTATGGAGCTCTCACATGTGGAAG<br>CTGCAAGGTCTTCTTCAAAAGAGCCGCTGAAGGGAAACAGAAGTACCTGTGCGCCA<br>GCAGAAATGATTGCACTATTGATAAATTCCGAAGGAAAAATTGTCCATCTTGTCGT<br>CTTCGGAAATGTTATGAAGCAGGGATGACTCTGGGAGCCCGGAAGCTGAAGAAACT<br>TGGTAATCTGAAACTACAGGAGGAAGGAGAGGCTTCCAGCACCACCAGCCCCACTG<br>AGGAGACAACCCAGAAGCTGACAGTGTCACACATTGAAGGCTATGAATGTCAGCCC |

| | |
|---|---|
| | ATCTTTCTGAATGTCCTGGAAGCCATTGAGCCAGGTGTAGTGTGTGCTGGACACGA<br>CAACAACCAGCCCGACTCCTTTGCAGCCTTGCTCTCTAGCCTCAATGAACTGGGAG<br>AGAGACAGCTTGTACACGTGGTCAAGTGGGCCAAGGCCTTGCCTGGCTTCCGCAAC<br>TTACACGTGGACGACCAGATGGCTGTCATTCAGTACTCCTGGATGGGGCTCATGGT<br>GTTTGCCATGGGCTGGCGATCCTTCACCAATGTCAACTCCAGGATGCTCTACTTCG<br>CCCCTGATCTGGTTTTCAATGAGTACCGCATGCACAAGTCCCGGATGTACAGCCAG<br>TGTGTCCGAATGAGGCACCTCTCTCAAGAGTTTGGATGGCTCCAAATCACCCCCCA<br>GGAATTCCTGTGCATGAAAGCACTGCTACTCTTCAGCATTATTCCAGTGGATGGGC<br>TGAAAAATCAAAAATTCTTTGATGAACTTCGAATGAACTACATCAAGGAACTCGAT<br>CGTATCATTGCATGCAAAAGAAAAAATCCCACATCCTGCTCAAGACGCTTCTACCA<br>GCTCACCAAGCTCCTGGACTCCGTGCAGCCTATTGCGAGAGAGCTGCATCAGTTCA<br>CTTTTGACCTGCTAATCAAGTCACACATGGTGAGCGTGGACTTTCCGGAAATGATG<br>GCAGAGATCATCTCTGTGCAAGTGCCCAAGATCCTTTCTGGGAAAGTCAAGCCCAT<br>CTATTTCCACACCCAGGGCTCGAGCGGTGGTGGCGGGAGCGGAGGTGGAGGGTCGT<br>CAGGTGTGACCGGCTACCGGCTGTTCGAGGAGATTCTGTAA |
| SOX2-smN cDNA (cloned into pcDNA3.1(+) backbone) | ATGGTGACCGGCTACCGGCTGTTCGAGGAGATTCTGTCTGGCTCGAGCGGTGGTGG<br>CGGGAGCGGAGGTGGAGGGTCGTCAGGTACCTACAACATGATGGAGACGGAGCTGA<br>AGCCGCCGGGCCCGCAGCAAACTTCGGGGGGCGGCGGCGGCAACTCCACCGCGGCG<br>GCGGCCGGCGGCAACCAGAAAAACAGCCCGGACCGCGTCAAGCGGCCCATGAATGC<br>CTTCATGGTGTGGTCCCGCGGGCAGCGGCGCAAGATGGCCCAGGAGAACCCCAAGA<br>TGCACAACTCGGAGATCAGCAAGCGCCTGGGCGCCGAGTGGAAACTTTTGTCGGAG<br>ACGGAGAAGCGGCCGTTCATCGACGAGGCTAAGCGGCTGCGAGCGCTGCACATGAA<br>GGAGCACCCGGATTATAAATACCGGCCCCGGCGGAAAACCAAGACGCTCATGAAGA<br>AGGATAAGTACACGCTGCCCGGCGGGCTGCTGGCCCCCGGCGGCAATAGCATGGCG<br>AGCGGGGTCGGGGTGGGCGCCGGCCTGGGCGCGGGCGTGAACCAGCGCATGGACAG<br>TTACGCGCACATGAACGGCTGGAGCAACGGCAGCTACAGCATGATGCAGGACCAGC<br>TGGGCTACCCGCAGCACCCGGGCCTCAATGCGCACGGCGCAGCGCAGATGCAGCCC<br>ATGCACCGCTACGACGTGAGCGCCCTGCAGTACAACTCCATGACCAGCTCGCAGAC<br>CTACATGAACGGCTCGCCCACCTACAGCATGTCCTACTCGCAGCAGGGCACCCCTG<br>GCATGGCTCTTGGCTCCATGGGTTCGGTGGTCAAGTCCGAGGCCAGCTCCAGCCCC<br>CCTGTGGTTACCTCTTCCTCCCACTCCAGGGCGCCCTGCCAGGCCGGGGACCTCCG<br>GGACATGATCAGCATGTATCTCCCCGGCGCCGAGGTGCCGGAACCCGCCGCCCCCA<br>GCAGACTTCACATGTCCCAGCACTACCAGAGCGGCCCGGTGCCCGGCACGGCCATT<br>AACGGCACACTGCCCCTCTCACACATGTGA |
| AR-lgN cDNA (cloned into pcDNA3.1(+) backbone) | ATGGTCTTCACACTCGAAGATTTCGTTGGGGACTGGGAACAGACAGCCGCCTACAA<br>CCTGGACCAAGTCCTTGAACAGGGAGGTGTGTCCAGTTTGCTGCAGAATCTCGCCG<br>TGTCCGTAACTCCGATCCAAAGGATTGTCCGGAGCGGTGAAAATGCCCTGAAGATC<br>GACATCCATGTCATCATCCCGTATGAAGGTCTGAGCGCCGACCAAATGGCCCAGAT<br>CGAAGAGGTGTTTAAGGTGGTGTACCCTGTGGATGATCATCACTTTAAGGTGATCC<br>TGCCCTATGGCACACTGGTAATCGACGGGGTTACGCCGAACATGCTGAACTATTTC<br>GGACGGCCGTATGAAGGCATCGCCGTGTTCGACGGCAAAAAGATCACTGTAACAGG<br>GACCCTGTGGAACGGCAACAAAATTATCGACGAGCGCCTGATCACCCCCGACGGCT<br>CCATGCTGTTCCGAGTAACCATCAACAGCTCTGGCTCGAGCGGTGGTGGCGGGAGC<br>GGAGGTGGAGGGTCGTCAGGTACCGAAGTGCAGTTAGGGCTGGGAAGGGTCTACCC<br>TCGGCCGCCGTCCAAGACCTACCGAGGAGCTTTCCAGAATCTGTTCCAGAGCGTGC<br>GCGAAGTGATCCAGAACCCGGGCCCCAGGCACCCAGAGGCCGCGAGCGCAGCACCT<br>CCCGGCGCCAGTTTGCTGCTGCTGCAGCAGCAGCAGCAGCAGCAGCAGCAGCAGCA<br>GCAGCAGCAGCAGCAGCAGCAGCAGCAGCAGCAGCAAGAGACTAGCCCCAGGCAGC<br>AGCAGCAGCAGCAGGGTGAGGATGGTTCTCCCCAAGCCCATCGTAGAGGCCCCACA<br>GGCTACCTGGTCCTGGATGAGGAACAGCAACCTTCACAGCCGCAGTCGGCCCTGGA<br>GTGCCACCCCGAGAGAGGTTGCGTCCCAGAGCCTGGAGCCGCCGTGGCCGCCAGCA<br>AGGGGCTGCCGCAGCAGCTGCCAGCACCTCCGGACGAGGATGACTCAGCTGCCCCA<br>TCCACGTTGTCCCTGCTGGGCCCCACTTTCCCCGGCTTAAGCAGCTGCTCCGCTGA<br>CCTTAAAGACATCCTGAGCGAGGCCAGCACCATGCAACTCCTTCAGCAACAGCAGC<br>AGGAAGCAGTATCCGAAGGCAGCAGCAGCGGGAGAGCGAGGGAGGCCTCGGGGGCT<br>CCCACTTCCTCCAAGGACAATTACTTAGGGGGCACTTCGACCATTTCTGACAACGC<br>CAAGGAGTTGTGTAAGGCAGTGTCGGTGTCCATGGGCCTGGGTGTGGAGGCGTTGG |

| | |
|---|---|
| | AGCATCTGAGTCCAGGGGAACAGCTTCGGGGGGATTGCATGTACGCCCCACTTTTG<br>GGAGTTCCACCCGCTGTGCGTCCCACTCCTTGTGCCCCATTGGCCGAATGCAAAGG<br>TTCTCTGCTAGACGACAGCGCAGGCAAGAGCACTGAAGATACTGCTGAGTATTCCC<br>CTTTCAAGGGAGGTTACACCAAAGGGCTAGAAGGCGAGAGCCTAGGCTGCTCTGGC<br>AGCGCTGCAGCAGGGAGCTCCGGGACACTTGAACTGCCGTCTACCCTGTCTCTCTA<br>CAAGTCCGGAGCACTGGACGAGGCAGCTGCGTACCAGAGTCGCGACTACTACAACT<br>TTCCACTGGCTCTGGCCGGACCGCCGCCCCCTCCGCCGCCTCCCCATCCCCACGCT<br>CGCATCAAGCTGGAGAACCCGCTGGACTACGGCAGCGCCTGGGCGGCTGCGGCGGC<br>GCAGTGCCGCTATGGGGACCTGGCGAGCCTGCATGGCGCGGGTGCAGCGGGACCCG<br>GTTCTGGGTCACCCTCAGCCGCCGCTTCCTCATCCTGGCACACTCTCTTCACAGCC<br>GAAGAAGGCCAGTTGTATGGACCGTGTGGTGGTGGTGGGGGTGGTGGCGGCGGCGG<br>CGGCGGCGGCGGCGGCGGCGGCGGCGGCGGCGGCGGCGGCGAGGCGGGAGCTGTAG<br>CCCCCTACGGCTACACTCGGCCCCCTCAGGGGCTGGCGGGCCAGGAAAGCGACTTC<br>ACCGCACCTGATGTGTGGTACCCTGGCGGCATGGTGAGCAGAGTGCCCTATCCCAG<br>TCCCACTTGTGTCAAAAGCGAAATGGGCCCCTGGATGGATAGCTACTCCGGACCTT<br>ACGGGGACATGCGTTTGGAGACTGCCAGGGACCATGTTTTGCCCATTGACTATTAC<br>TTTCCACCCCAGAAGACCTGCCTGATCTGTGGAGATGAAGCTTCTGGGTGTCACTA<br>TGGAGCTCTCACATGTGGAAGCTGCAAGGTCTTCTTCAAAAGAGCCGCTGAAGGGA<br>AACAGAAGTACCTGTGCGCCAGCAGAAATGATTGCACTATTGATAAATTCCGAAGG<br>AAAAATTGTCCATCTTGTCGTCTTCGGAAATGTTATGAAGCAGGGATGACTCTGGG<br>AGCCCGGAAGCTGAAGAAACTTGGTAATCTGAAACTACAGGAGGAAGGAGAGGCTT<br>CCAGCACCACCAGCCCCACTGAGGAGACAACCCAGAAGCTGACAGTGTCACACATT<br>GAAGGCTATGAATGTCAGCCCATCTTTCTGAATGTCCTGGAAGCCATTGAGCCAGG<br>TGTAGTGTGTGCTGGACACGACAACAACCAGCCCGACTCCTTTGCAGCCTTGCTCT<br>CTAGCCTCAATGAACTGGGAGAGAGACAGCTTGTACACGTGGTCAAGTGGGCCAAG<br>GCCTTGCCTGGCTTCCGCAACTTACACGTGGACGACCAGATGGCTGTCATTCAGTA<br>CTCCTGGATGGGGCTCATGGTGTTTGCCATGGGCTGGCGATCCTTCACCAATGTCA<br>ACTCCAGGATGCTCTACTTCGCCCCTGATCTGGTTTTCAATGAGTACCGCATGCAC<br>AAGTCCCGGATGTACAGCCAGTGTGTCCGAATGAGGCACCTCTCTCAAGAGTTTGG<br>ATGGCTCCAAATCACCCCCCAGGAATTCCTGTGCATGAAAGCACTGCTACTCTTCA<br>GCATTATTCCAGTGGATGGGCTGAAAAATCAAAAATTCTTTGATGAACTTCGAATG<br>AACTACATCAAGGAACTCGATCGTATCATTGCATGCAAAAGAAAAAATCCCACATC<br>CTGCTCAAGACGCTTCTACCAGCTCACCAAGCTCCTGGACTCCGTGCAGCCTATTG<br>CGAGAGAGCTGCATCAGTTCACTTTTGACCTGCTAATCAAGTCACACATGGTGAGC<br>GTGGACTTTCCGGAAATGATGGCAGAGATCATCTCTGTGCAAGTGCCCAAGATCCT<br>TTCTGGGAAAGTCAAGCCCATCTATTTCCACACCCAGTGA |

### Supplemental Table 2:

**A**

LNCaP AR Gene Candidates (GO Biological Processes)

| Gene Ontology (p<0.05) | -log(p-value) |
|---|---|
| Flavone Metabolic Process (GO:0051552) | 2.88151666 |
| Cellular Hyperosmotic Response (GO:0071474) | 2.663097017 |
| Monocarboxylic Acid Biosynthetic Process (GO:0072330) | 2.341359499 |
| Hyperosmotic Response (GO:0006972) | 2.213219495 |
| Regulation of Neuron Projection Arborization (GO:0150011) | 2.139703544 |
| Glucuronate Metabolic Process (GO:0019585) | 2.052273516 |
| Regulation of Skeletal Muscle Contraction by Calcium Ion Signaling (GO:0014722) | 2.052273516 |
| Skeletal Muscle Contraction via Release of Sequestered Calcium Ion (GO:0014809) | 2.052273516 |
| Estrogen Metabolic Process (GO:0008210) | 1.871172238 |
| Regulation of Fatty Acid Oxidation (GO:0046320) | 1.821619223 |
| Negative Regulation of Fatty Acid Metabolic Process (GO:0045922) | 1.812971436 |
| Positive Regulation of Carbohydrate Metabolic Process (GO:0045913) | 1.705645748 |
| Positive Regulation of Epithelial Tube Formation (GO:1905278) | 1.676115646 |
| NMDA Selective Glutamate Receptor Signaling Pathway (GO:0098989) | 1.444038425 |
| Negative Regulation of Very-Low-Density Lipoprotein Particle Remodeling (GO:0010903) | 1.444038425 |
| Retinoic Acid Metabolic Process (GO:0042573) | 1.397300754 |
| Positive Regulation of Glycolytic Process (GO:0045821) | 1.377837979 |
| Positive Regulation of Purine Nucleotide Catabolic Process (GO:0033123) | 1.377837979 |
| Embryonic Skeletal Joint Morphogenesis (GO:0060272) | 1.365195837 |
| Maintenance of Lens Transparency (GO:0036438) | 1.365195837 |
| Negative Regulation of B Cell Apoptotic Process (GO:0002903) | 1.365195837 |
| Positive Regulation of Neuron Projection Arborization (GO:0150012) | 1.365195837 |
| Regulation of Skeletal Muscle Fiber Development (GO:0048742) | 1.365195837 |
| Regulation of Very-Low-Density Lipoprotein Particle Remodeling (GO:0010901) | 1.365195837 |
| Regulation of Fatty Acid Metabolic Process (GO:0019217) | 1.35974487 |
| Regulation of Cellular Response to Insulin Stimulus (GO:1900076) | 1.329875175 |
| Xenobiotic Transport (GO:0042908) | 1.329646183 |
| Regulation of Dephosphorylation (GO:0035303) | 1.315817201 |

**B**

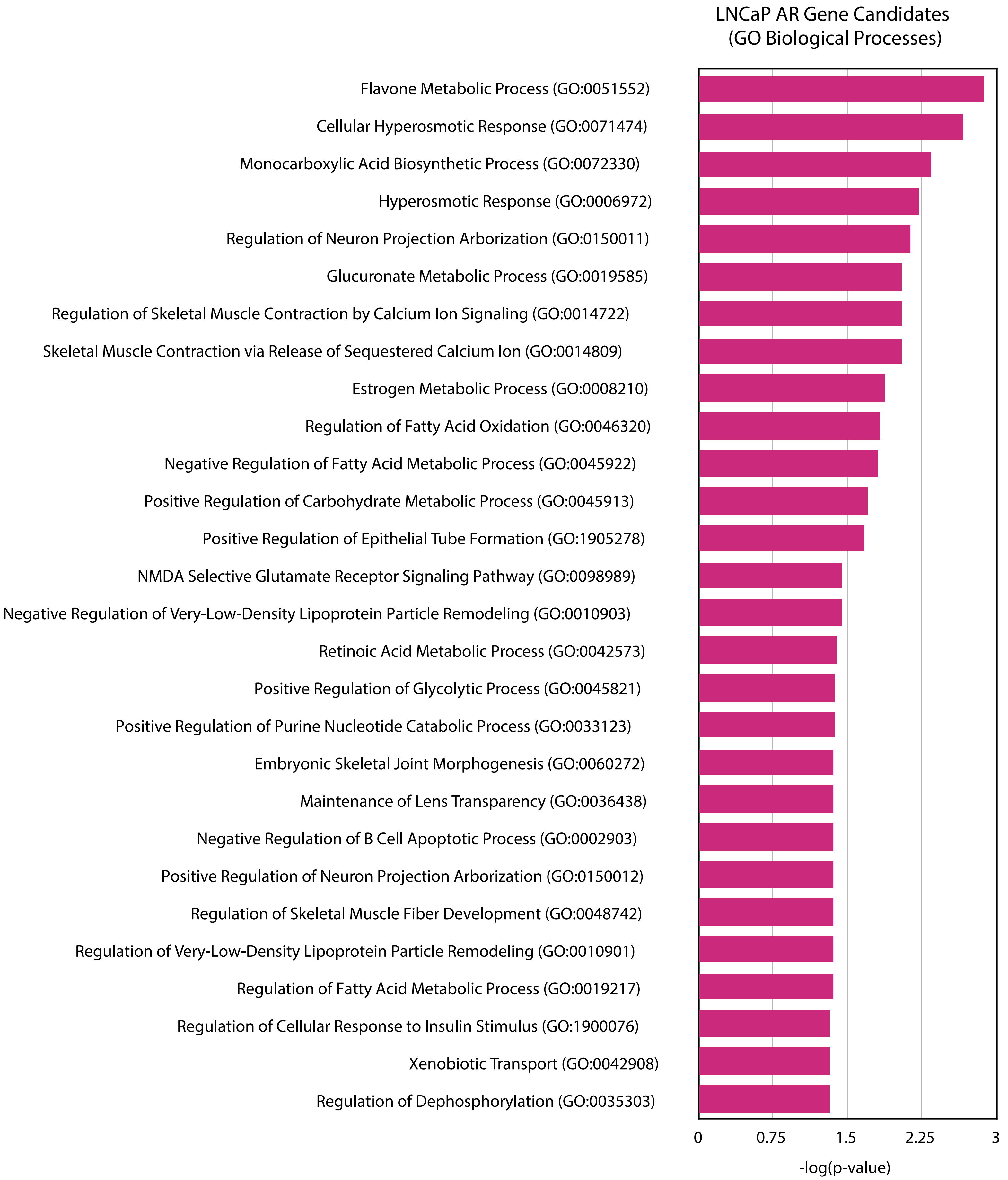

**C**

LNCaP METTL3 Gene Candidates (GO Biological Processes)

| Gene Ontology (p<0.05) | -log(p-value) |
|---|---|
| Negative Regulation of Locomotion (GO:0040013) | 2.587892911 |
| Negative Regulation of Response to Endoplasmic Reticulum Stress (GO:1903573) | 2.339196649 |
| Regulation of Antifungal Innate Immune Response (GO:1905034) | 2.333326392 |
| Tryptophan Transport (GO:0015827) | 2.333326392 |
| DNA Modification (GO:0006304) | 2.168057014 |
| Neutrophil-mediated Killing of Symbiont Cell (GO:0070943) | 2.163601811 |
| Nucleoside Triphosphate Biosynthetic Process (GO:0009142) | 2.163601811 |
| Fructose 1,6-Bisphosphate Metabolic Process (GO:0030388) | 2.163601811 |
| Positive Regulation of Proteasomal Protein Catabolic Process (GO:1901800) | 2.152828299 |
| Inner Mitochondrial Membrane Organization (GO:0007007) | 2.062127514 |
| Negative Regulation of ERAD Pathway (GO:1904293) | 2.023828795 |
| Neutrophil-mediated Killing of Bacterium (GO:0070944) | 2.023828795 |
| Glutathione Catabolic Process (GO:0006751) | 2.023828795 |
| Sulfur Compound Catabolic Process (GO:0044273) | 1.920469792 |
| Poly(A)-dependent snoRNA 3'-End Processing (GO:0071051) | 1.905233322 |
| U4 snRNA 3'-End Processing (GO:0034475) | 1.905233322 |
| SnRNA 3'-End Processing (GO:0034472) | 1.866812043 |
| Canonical Glycolysis (GO:0061621) | 1.815851822 |
| Glucose Catabolic Process to Pyruvate (GO:0061718) | 1.815851822 |
| Zinc Ion Transmembrane Transport (GO:0071577) | 1.815851822 |
| Anoikis (GO:0043276) | 1.802420288 |
| T Cell Apoptotic Process (GO:0070231) | 1.802420288 |
| Glycolytic Process Through Glucose-6-Phosphate (GO:0061620) | 1.767357824 |
| Zinc Ion Transport (GO:0006829) | 1.767357824 |
| Fat Cell Differentiation (GO:0045444) | 1.718522813 |
| Apoptotic DNA Fragmentation (GO:0006309) | 1.711829823 |
| Aromatic Amino Acid Transport (GO:0015801) | 1.711829823 |
| DNA Alkylation Repair (GO:0006307) | 1.711829823 |
| Nucleoside Triphosphate Metabolic Process (GO:0009141) | 1.711829823 |
| Fructose 6-Phosphate Metabolic Process (GO:0006002) | 1.711829823 |
| Regulation of Mesenchymal Stem Cell Differentiation (GO:2000739) | 1.630987257 |
| Nuclear mRNA Surveillance (GO:0071028) | 1.630987257 |
| Transferrin Transport (GO:0033572) | 1.630987257 |
| Defense Response to Symbiont (GO:0140546) | 1.558101638 |
| Sno(s)RNA 3'-End Processing (GO:0031126) | 1.558101638 |
| ADP Catabolic Process (GO:0046032) | 1.51831778 |
| Glycolytic Process (GO:0006096) | 1.51831778 |
| Regulation of DNA-templated Transcription Initiation (GO:2000142) | 1.49183457 |
| Negative Regulation of Chemotaxis (GO:0050922) | 1.49183457 |
| Brown Fat Cell Differentiation (GO:0050873) | 1.49183457 |
| DNA Deamination (GO:0045006) | 1.49183457 |
| Presynapse Organization (GO:0099172) | 1.49183457 |
| Mitochondrial Electron Transport, Ubiquinol to Cytochrome C (GO:0006122) | 1.49183457 |
| Nuclear RNA Surveillance (GO:0071027) | 1.44809175 |
| Cristae Formation (GO:0042407) | 1.431159313 |
| Peptidyl-L-cysteine S-palmitoylation (GO:0018230) | 1.431159313 |
| Peptidyl-S-diacylglycerol-L-cysteine Biosynthetic Process From Peptidyl-Cysteine (GO:0018231) | 1.431159313 |
| Interleukin-6-mediated Signaling Pathway (GO:0070102) | 1.431159313 |
| Purine Ribonucleoside Triphosphate Biosynthetic Process (GO:0009206) | 1.431159313 |
| Endocytic Recycling (GO:0032456) | 1.418809967 |
| Pyridine Nucleotide Catabolic Process (GO:0019364) | 1.414920832 |
| Vascular Transport (GO:0010232) | 1.38166313 |
| Cellular Response to Growth Hormone Stimulus (GO:0071378) | 1.375270833 |
| Early Endosome to Golgi Transport (GO:0034498) | 1.375270833 |
| Transport Across Blood-Brain Barrier (GO:0150104) | 1.345727134 |
| Neg Reg Endoplasmic Reticulum Stress-Induced Intrinsic Apoptotic Signaling Pathway (GO:1902236) | 1.323526169 |
| Nitric Oxide Biosynthetic Process (GO:0006809) | 1.323526169 |
| GTP Metabolic Process (GO:0046039) | 1.323526169 |
| Negative Regulation of Intrinsic Apoptotic Signaling Pathway (GO:2001243) | 1.320269801 |
| Positive Regulation of Ubiquitin-Dependent Protein Catabolic Process (GO:2000060) | 1.310945099 |

**D**

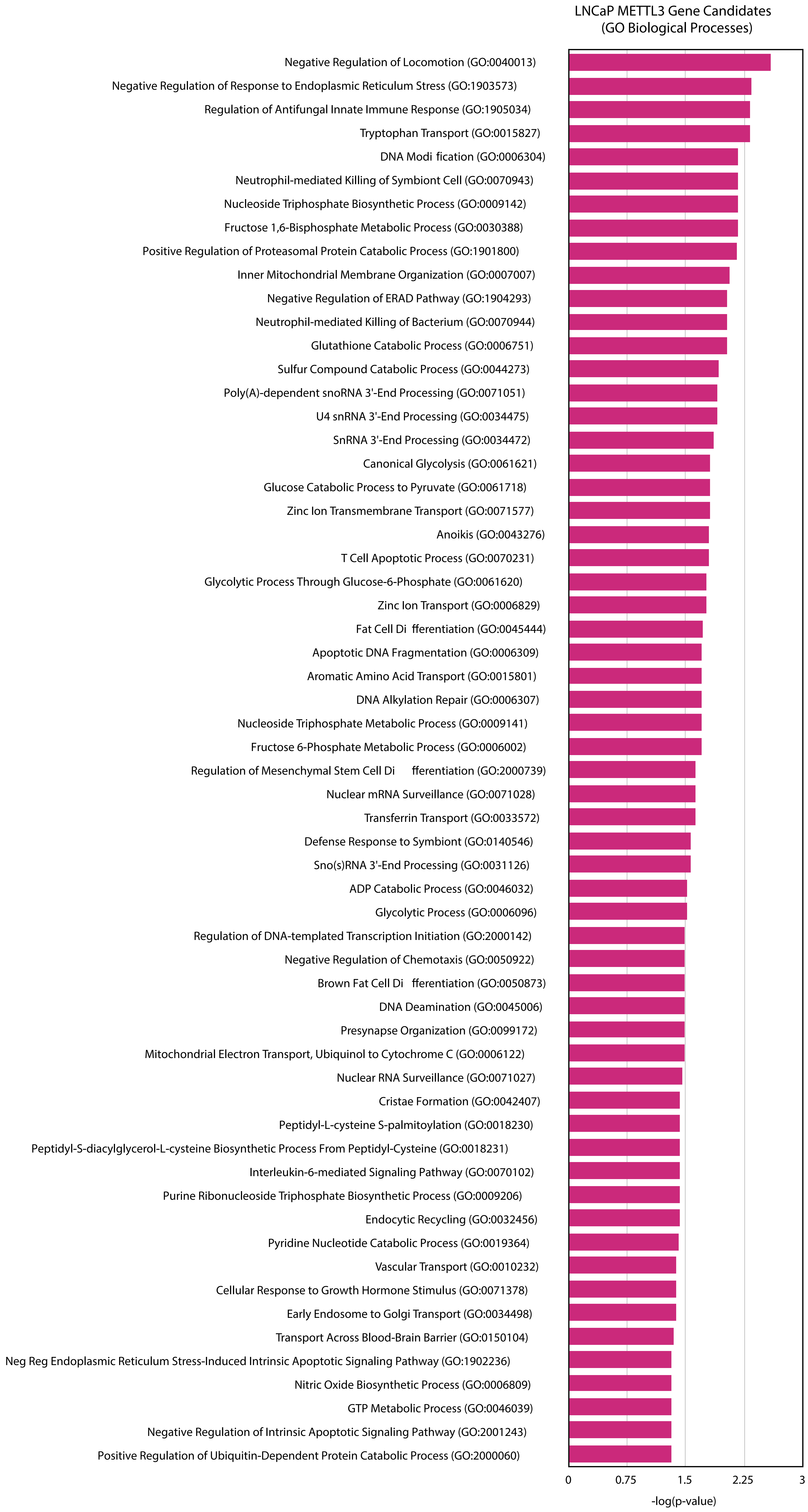

**E**

LNCaP AR/METTL3 Shared Gene Candidates (GO Biological Processes)

| Gene Ontology (p<0.05) | -log(p-value) |
|---|---|
| Antibody-dependent Cellular Cytotoxicity (GO:0001788) | 8.89858 |
| Type IIa Hypersensitivity (GO:0001794) | 8.89858 |
| Protein Localization to Condensed Chromosome (GO:1903083) | 4.59905 |
| Protein Localization to Kinetochore (GO:0034501) | 4.40015 |
| Protein Localization to Chromosome, Centromeric Region (GO:0071459) | 3.51574 |
| Hepoxilin Biosynthetic Process (GO:0051122) | 2.75219 |
| Mitochondrial Translational Elongation (GO:0070125) | 2.75219 |
| Regulation of Hormone Metabolic Process (GO:0032350) | 2.58684 |
| Lipoxygenase Pathway (GO:0019372) | 2.44266 |
| Sphingomyelin Metabolic Process (GO:0006684) | 2.42773 |
| Regulation of Phagocytosis (GO:0050764) | 2.33681 |
| Regulation of Mitotic Sister Chromatid Segregation (GO:0033047) | 2.31507 |
| Reactive Oxygen Species Biosynthetic Process (GO:1903409) | 2.20081 |
| Sphingomyelin Biosynthetic Process (GO:0006686) | 2.09754 |
| Adenosine Transport (GO:0032238) | 1.99243 |
| Regulation of Hydrogen Peroxide Biosynthetic Process (GO:0010728) | 1.99243 |
| Mitochondrial Translational Termination (GO:0070126) | 1.99243 |
| Positive Regulation of CD4-positive, Alpha-Beta T Cell Proliferation (GO:2000563) | 1.99243 |
| Organelle Transport Along Microtubule (GO:0072384) | 1.96559 |
| Cellular Response to Growth Hormone Stimulus (GO:0071378) | 1.91718 |
| Negative Regulation of Phagocytosis (GO:0050765) | 1.83761 |
| Pyrimidine Nucleoside Transport (GO:0015864) | 1.82589 |
| Uridine Transmembrane Transport (GO:0015862) | 1.82589 |
| Natural Killer Cell Degranulation (GO:0043320) | 1.82589 |
| Negative Regulation of Lamellipodium Organization (GO:1902744) | 1.82589 |
| Nucleoside Transport (GO:0015858) | 1.82589 |
| Podocyte Cell Migration (GO:0090521) | 1.82589 |
| Polyamine Transport (GO:0015846) | 1.82589 |
| Positive Regulation of Mitotic Sister Chromatid Segregation (GO:0062033) | 1.82589 |
| Purine Nucleobase Transmembrane Transport (GO:1904823) | 1.82589 |
| Positive Regulation of Tumor Necrosis Factor Production (GO:0032760) | 1.80249 |
| Sphingolipid Metabolic Process (GO:0006665) | 1.73095 |
| Positive Regulation of Tumor Necrosis Factor Superfamily Cytokine Production (GO:1903557) | 1.70745 |
| Regulation of Delayed Rectifier Potassium Channel Activity (GO:1902259) | 1.68929 |
| Negative Regulation of Humoral Immune Response (GO:0002921) | 1.68929 |
| Negative Regulation of Ruffle Assembly (GO:1900028) | 1.68929 |
| Pos Reg Phospholipase C-activating G Protein-Coupled Rec Signaling Pathway (GO:1900738) | 1.68929 |
| Protein Ufmylation (GO:0071569) | 1.68929 |
| Glutamate Metabolic Process (GO:0006536) | 1.63119 |
| Linoleic Acid Metabolic Process (GO:0043651) | 1.63119 |
| Positive Regulation of ATP-dependent Activity (GO:0032781) | 1.63119 |
| Regulation of Thyroid Hormone Generation (GO:2000609) | 1.57384 |
| Hydrogen Peroxide Biosynthetic Process (GO:0050665) | 1.57384 |
| Vacuole Organization (GO:0007033) | 1.57384 |
| Positive Regulation of Cellular Component Biogenesis (GO:0044089) | 1.53401 |
| Negative Regulation of Endocytosis (GO:0045806) | 1.52256 |
| Establishment of Skin Barrier (GO:0061436) | 1.51472 |
| Regulation of Metal Ion Transport (GO:0010959) | 1.48398 |
| Regulation of Lysosome Size (GO:0062196) | 1.47417 |
| Regulation of Phospholipase C-activating G Protein-Coupled Receptor Signaling Pathway (GO:1900736) | 1.47417 |
| Establishment of Vesicle Localization (GO:0051650) | 1.47417 |
| Negative Regulation of Leukocyte Proliferation (GO:0070664) | 1.47417 |
| Positive Regulation of Defense Response to Bacterium (GO:1900426) | 1.47417 |
| Positive Regulation of Hemostasis (GO:1900048) | 1.47417 |
| Positive Regulation of G2/M Transition of Mitotic Cell Cycle (GO:0010971) | 1.46154 |
| Protein Insertion Into Membrane (GO:0051205) | 1.46154 |
| Spindle Assembly (GO:0051225) | 1.45238 |
| Growth Hormone Receptor Signaling Pathway (GO:0060396) | 1.41132 |
| Synaptic Vesicle Cycle (GO:0099504) | 1.41084 |
| Regulation of Attachment of Mitotic Spindle Microtubules to Kinetochore (GO:1902423) | 1.38670 |
| Regulation of Defense Response to Bacterium (GO:1900424) | 1.38670 |
| Regulation of Endosome Size (GO:0051036) | 1.38670 |
| Negative Regulation of B Cell Receptor Signaling Pathway (GO:0050859) | 1.38670 |
| Neuroepithelial Cell Differentiation (GO:0060563) | 1.38670 |
| Positive Regulation of Membrane Depolarization (GO:1904181) | 1.38670 |
| Purine Nucleobase Transport (GO:0006863) | 1.38670 |
| Skin Epidermis Development (GO:0098773) | 1.36378 |
| Positive Regulation of Neural Precursor Cell Proliferation (GO:2000179) | 1.36378 |
| Regulation of ERAD Pathway (GO:1904292) | 1.31870 |
| Positive Regulation of Cell Cycle G2/M Phase Transition (GO:1902751) | 1.31870 |
| Regulation of CD4-positive, Alpha-Beta T Cell Proliferation (GO:2000561) | 1.30896 |
| Translational Termination (GO:0006415) | 1.30896 |
| Negative Regulation of Cell Division (GO:0051782) | 1.30896 |
| Negative Regulation of Vascular Endothelial Growth Factor Receptor Signaling Pathway (GO:0030948) | 1.30896 |
| Positive Regulation of Chromosome Segregation (GO:0051984) | 1.30896 |
| Positive Regulation of Cilium Movement (GO:0003353) | 1.30896 |
| Positive Regulation of Cilium-Dependent Cell Motility (GO:2000155) | 1.30896 |
| Positive Regulation of Flagellated Sperm Motility (GO:1902093) | 1.30896 |
| Protein Localization to Nuclear Envelope (GO:0090435) | 1.30896 |

**F**

LNCaP AR/METTL3 Shared Gene Candidates (GO Biological Processes)

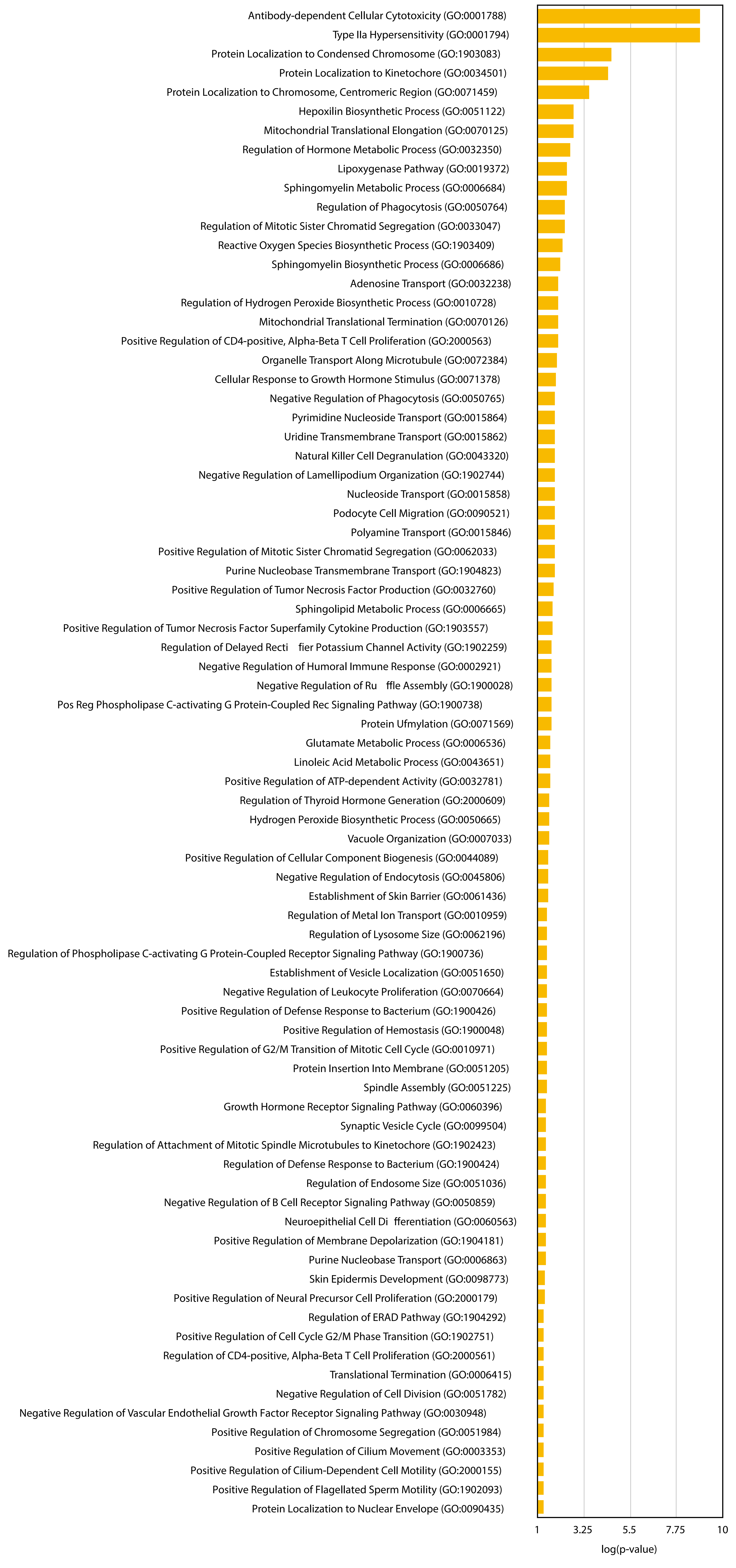


**G**

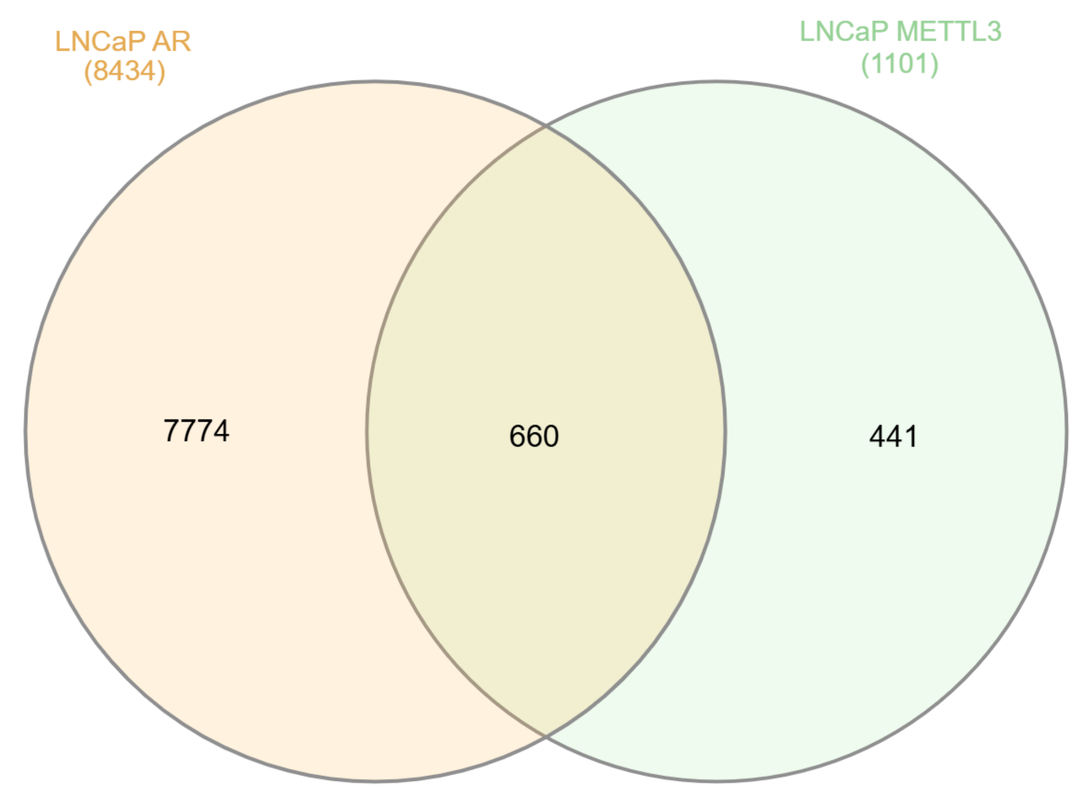


LNCaP ChIP-seq Gene Candidates (adjRPscore >0.01)

**H**

22Rv1 AR Gene Candidates (GO Biological Processes)

| Gene Ontology (p<0.05) | -log(p-value) |
|---|---|
| Regulation of Fatty Acid Oxidation (GO:0046320) | 2.873944518 |
| Antibody-dependent Cellular Cytotoxicity (GO:0001788) | 2.728396648 |
| Type IIa Hypersensitivity (GO:0001794) | 2.728396648 |
| Positive Regulation of Mesenchymal Stem Cell Differentiation (GO:2000741) | 2.273562322 |
| Regulation of Skeletal Muscle Contraction by Calcium Ion Signaling (GO:0014722) | 2.273562322 |
| Negative Regulation of VEGF Receptor Signaling Pathway (GO:0030948) | 1.904189876 |
| Regulation of Tube Diameter (GO:0035296) | 1.892014645 |
| Vascular Process in Circulatory System (GO:0003018) | 1.892014645 |
| Response to Fatty Acid (GO:0070542) | 1.698513856 |
| Negative Regulation of Hydrogen Peroxide-Mediated Programmed Cell Death (GO:1901299) | 1.64562505 |
| Negative Regulation of Very-Low-Density Lipoprotein Particle Remodeling (GO:0010903) | 1.64562505 |
| Positive Regulation of Toll-Like Receptor 2 Signaling Pathway (GO:0034137) | 1.64562505 |
| Regulation of Establishment of T Cell Polarity (GO:1903903) | 1.64562505 |
| Regulation of Adenylate Cyclase-Activating GPCR Signaling Pathway (GO:0106070) | 1.632371626 |
| Endoplasmic Reticulum Tubular Network Membrane Organization (GO:1990809) | 1.590285375 |
| Regulation of Skeletal Muscle Fiber Development (GO:0048742) | 1.590285375 |
| Regulation of Very-Low-Density Lipoprotein Particle Remodeling (GO:0010901) | 1.590285375 |
| Cellular Response to Fatty Acid (GO:0071398) | 1.494706251 |
| Bile Acid Metabolic Process (GO:0008206) | 1.461922969 |
| Mature B Cell Differentiation Involved in Immune Response (GO:0002313) | 1.3671109 |
| Androgen Metabolic Process (GO:0008209) | 1.36478218 |
| Positive Regulation of Glycolytic Process (GO:0045821) | 1.36478218 |
| Positive Regulation of Purine Nucleotide Catabolic Process (GO:0033123) | 1.36478218 |
| Alanine Transport (GO:0032328) | 1.328283488 |

**I**

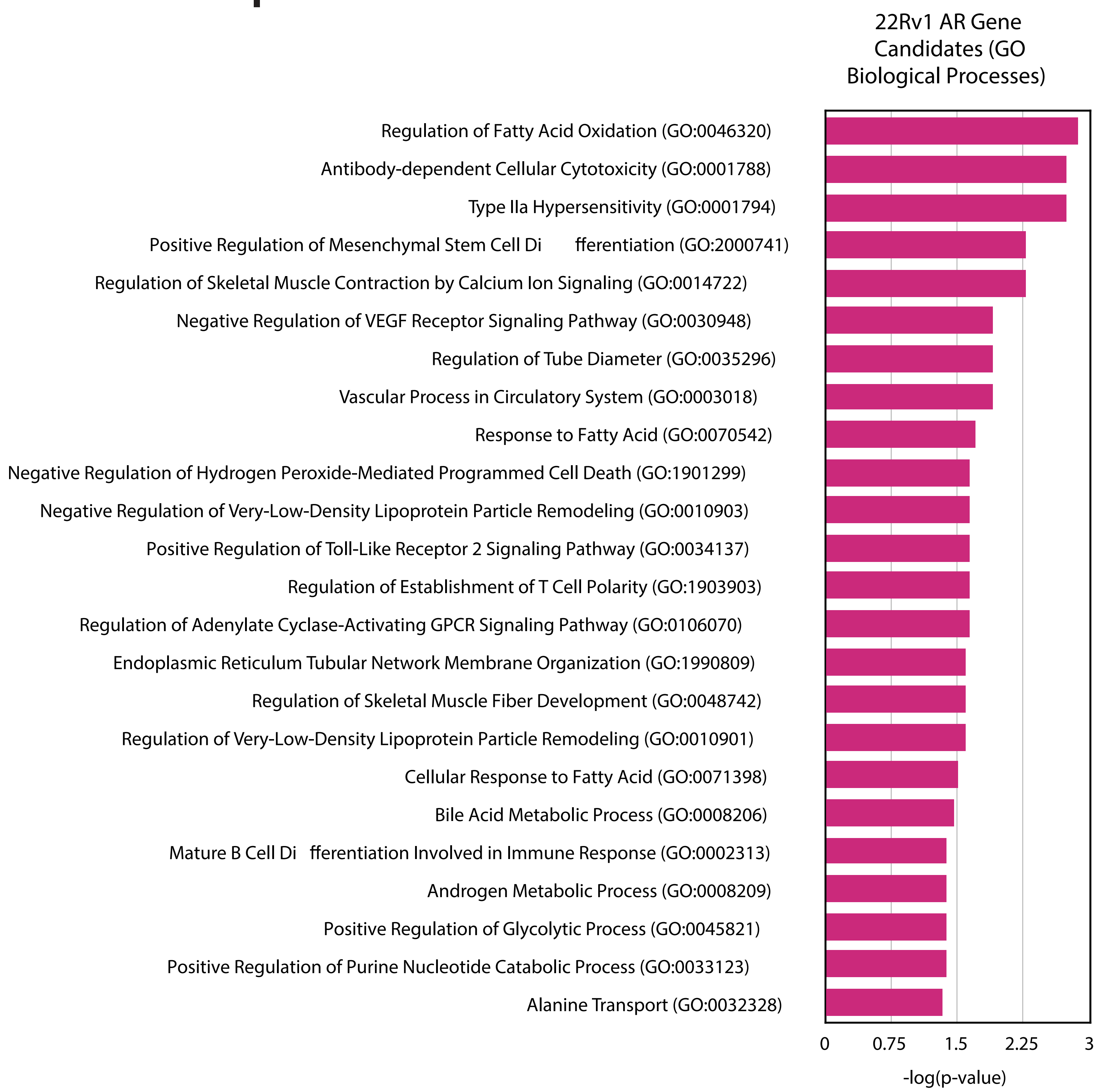

J

22Rv1 METTL3 Gene Candidates (GO Biological Processes)

| Gene Ontology (p<0.05) | -log(p-value) |
|---|---|
| Mitochondrial Translation (GO:0032543) | 5.251389081 |
| Chromatin Remodeling (GO:0006338) | 4.805324939 |
| Mitochondrial Gene Expression (GO:0140053) | 3.837708387 |
| Negative Regulation of DNA-templated Transcription (GO:0045892) | 3.707597674 |
| Positive Regulation of Neuroblast Proliferation (GO:0002052) | 3.271724804 |
| Translation (GO:0006412) | 3.243218722 |
| Regulation of DNA-templated Transcription Elongation (GO:0032784) | 2.983889324 |
| Mitochondrial Respiratory Chain Complex Assembly (GO:0033108) | 2.974654085 |
| RNA Splicing (GO:0008380) | 2.829925868 |
| RRNA Metabolic Process (GO:0016072) | 2.693443175 |
| RRNA Processing (GO:0006364) | 2.689083022 |
| Ribosome Biogenesis (GO:0042254) | 2.675599762 |
| RNA Processing (GO:0006396) | 2.625349211 |
| MRNA Transcription by RNA Polymerase II (GO:0042789) | 2.618086442 |
| Regulation of Neuroblast Proliferation (GO:1902692) | 2.595541968 |
| Mitochondrial Translational Elongation (GO:0070125) | 2.586261973 |
| Regulation of G1/S Transition of Mitotic Cell Cycle (GO:2000045) | 2.537036046 |
| Negative Regulation of Catalytic Activity (GO:0043086) | 2.498560816 |
| Negative Regulation of Transcription by RNA Polymerase II (GO:0000122) | 2.49562031 |
| Mitochondrial Respiratory Chain Complex IV Assembly (GO:0033617) | 2.44490875 |
| Negative Regulation of RNA Biosynthetic Process (GO:1902679) | 2.416656889 |
| Regulation of Chromatin Organization (GO:1902275) | 2.39591678 |
| Positive Regulation of DNA-templated Transcription (GO:0045893) | 2.38727274 |
| MRNA Transcription (GO:0009299) | 2.333718862 |
| Positive Regulation of RNA Biosynthetic Process (GO:1902680) | 2.322406244 |
| MRNA Splicing, via Spliceosome (GO:0000398) | 2.311539468 |
| Intracellular Protein Transport (GO:0006886) | 2.282664963 |
| Ribosome Disassembly (GO:0032790) | 2.282229395 |
| Long-term Synaptic Depression (GO:0060292) | 2.2798177 |
| Respiratory Chain Complex IV Assembly (GO:0008535) | 2.247513008 |
| Alternative mRNA Splicing, via Spliceosome (GO:0000380) | 2.222296246 |
| Response to Fibroblast Growth Factor (GO:0071774) | 2.222296246 |
| Epigenetic Regulation of Gene Expression (GO:0040029) | 2.17945386 |
| Translational Termination (GO:0006415) | 2.153752597 |
| Protein Deneddylation (GO:0000338) | 2.153752597 |
| Negative Regulation of RNA Splicing (GO:0033119) | 2.129205516 |
| Regulation of Double-Strand Break Repair (GO:2000779) | 2.075330325 |
| Regulation of RNA Biosynthetic Process (GO:2001141) | 2.030745417 |
| Ribosomal Large Subunit Biogenesis (GO:0042273) | 2.004094457 |
| SnRNA 3'-End Processing (GO:0034472) | 1.999190017 |
| Negative Regulation of DNA-templated Transcription, Elongation (GO:0032785) | 1.999190017 |
| Chromatin Organization (GO:0006325) | 1.972695229 |
| Regulation of Intracellular Steroid Hormone Receptor Signaling Pathway (GO:0033143) | 1.932900667 |
| Positive Regulation of Neural Precursor Cell Proliferation (GO:2000179) | 1.932900667 |
| MRNA Export From Nucleus (GO:0006406) | 1.889532163 |
| Protein Import Into Nucleus (GO:0006606) | 1.8889696 |
| SnRNA Transport (GO:0051030) | 1.8808482 |
| Vesicle Tethering to Golgi (GO:0099041) | 1.8808482 |
| Negative Regulation of Cell Size (GO:0045792) | 1.8808482 |
| Negative Regulation of Pinocytosis (GO:0048550) | 1.8808482 |
| RNA Splicing, via Transesterification Reactions (GO:0000375) | 1.870052827 |
| RRNA Methylation (GO:0031167) | 1.870052827 |
| U2-type Prespliceosome Assembly (GO:1903241) | 1.870052827 |
| Negative Regulation of DNA Damage Response, Signal Transduction by P53 Class Mediator (GO:0043518) | 1.846698232 |
| Negative Regulation of Stem Cell Differentiation (GO:2000737) | 1.846698232 |
| Negative Regulation of Transport (GO:0051051) | 1.824477494 |
| Spliceosomal snRNP Assembly (GO:0000387) | 1.791830765 |
| Establishment of Protein Localization to Organelle (GO:0072594) | 1.774638736 |
| RNA Methylation (GO:0001510) | 1.769449301 |
| Negative Regulation of mRNA Splicing, via Spliceosome (GO:0048025) | 1.753554411 |
| Transcription by RNA Polymerase II (GO:0006366) | 1.748071126 |
| Protein Localization to Nucleus (GO:0034504) | 1.744784736 |
| MRNA Processing (GO:0006397) | 1.743694849 |
| Regulation of Translational Termination (GO:0006449) | 1.715675559 |
| Glycosyl Compound Biosynthetic Process (GO:1901659) | 1.715675559 |
| Wybutosine Biosynthetic Process (GO:0031591) | 1.715675559 |
| Negative Regulation of Fibroblast Migration (GO:0010764) | 1.715675559 |
| Negative Regulation of Lipid Transport (GO:0032369) | 1.715675559 |
| Nucleoside Triphosphate Biosynthetic Process (GO:0009142) | 1.715675559 |
| Positive Regulation of Kidney Development (GO:0090184) | 1.715675559 |
| Regulation of Intracellular Estrogen Receptor Signaling Pathway (GO:0033146) | 1.699422674 |
| Negative Regulation of mRNA Processing (GO:0050686) | 1.699422674 |
| Nuclear-transcribed mRNA Catabolic Process, Nonsense-Mediated Decay (GO:0000184) | 1.699422674 |
| Regulation of Transcription by RNA Polymerase II (GO:0006357) | 1.68926359 |
| Ubiquitin-dependent Protein Catabolic Process via the C-end Degron Rule Pathway (GO:0140627) | 1.683860503 |
| Protein K6-linked Ubiquitination (GO:0085020) | 1.683860503 |
| RNA Splicing, via Transesterification Reactions With Bulged Adenosine as Nucleophile (GO:0000377) | 1.657277389 |
| Regulation of Protein Modification by Small Protein Conjugation or Removal (GO:1903320) | 1.647760212 |
| RRNA Catabolic Process (GO:0016075) | 1.611625639 |
| Regulation of Protein Neddylation (GO:2000434) | 1.611625639 |
| Translational Elongation (GO:0006414) | 1.611625639 |
| Negative Regulation of Transcription Elongation by RNA Polymerase II (GO:0034244) | 1.611625639 |
| Protein Localization to Cell-Cell Junction (GO:0150105) | 1.611625639 |
| Regulation of Transcription by RNA Polymerase III (GO:0006359) | 1.598388179 |
| Regulation of Cellular Response to Stress (GO:0080135) | 1.589087391 |
| Regulation of Blood-Brain Barrier Permeability (GO:1905603) | 1.580430272 |
| Clathrin-coated Vesicle Cargo Loading (GO:0035652) | 1.580430272 |
| Clathrin-coated Vesicle Cargo Loading, AP-3-mediated (GO:0035654) | 1.580430272 |
| Regulation of Mitochondrial Transcription (GO:1903108) | 1.580430272 |
| Regulation of Vascular Associated Smooth Muscle Cell Apoptotic Process (GO:1905459) | 1.580430272 |
| Rho-activating G Protein-Coupled Receptor Signaling Pathway (GO:0160221) | 1.580430272 |
| Selenocysteine Incorporation (GO:0001514) | 1.580430272 |
| Transcription-dependent Tethering of RNA Polymerase II Gene DNA at Nuclear Periphery (GO:0000972) | 1.580430272 |
| Translational Readthrough (GO:0006451) | 1.580430272 |
| Negative Regulation of Cyclin-Dependent Protein Kinase Activity (GO:1904030) | 1.580430272 |
| Negative Regulation of Cyclin-Dependent Protein Serine/Threonine Kinase Activity (GO:0045736) | 1.580430272 |
| Negative Regulation of Glial Cell Proliferation (GO:0060253) | 1.580430272 |
| Positive Regulation of Fibroblast Growth Factor Receptor Signaling Pathway (GO:0045743) | 1.580430272 |
| Positive Regulation of Sterol Transport (GO:0032373) | 1.580430272 |
| Response to UV (GO:0009411) | 1.545013669 |
| RRNA Modification (GO:0000154) | 1.481905137 |
| NLS-bearing Protein Import Into Nucleus (GO:0006607) | 1.481905137 |
| Positive Regulation of Mitotic Cell Cycle Phase Transition (GO:1901992) | 1.478874056 |
| Cellular Response to Interleukin-9 (GO:0071355) | 1.466338187 |
| Regulation of Organ Growth (GO:0046620) | 1.466338187 |
| Interleukin-9-mediated Signaling Pathway (GO:0038113) | 1.466338187 |
| Locomotor Rhythm (GO:0045475) | 1.466338187 |
| Mammary Gland Epithelial Cell Differentiation (GO:0060644) | 1.466338187 |
| U4 snRNA 3'-End Processing (GO:0034475) | 1.466338187 |
| Membrane Tubulation (GO:0097749) | 1.466338187 |
| Negative Regulation of Inflammatory Response to Wounding (GO:0106015) | 1.466338187 |
| Negative Regulation of Response to Type II Interferon (GO:0060331) | 1.466338187 |
| Negative Regulation of Type II Interferon-Mediated Signaling Pathway (GO:0060336) | 1.466338187 |
| Positive Regulation of Astrocyte Differentiation (GO:0048711) | 1.466338187 |
| Nuclear RNA Surveillance (GO:0071027) | 1.462490807 |
| Regulation of Cell Cycle Process (GO:0010564) | 1.429937071 |
| Regulation of Extrinsic Apoptotic Signaling Pathway (GO:2001236) | 1.42240611 |
| Regulation of Gene Expression (GO:0010468) | 1.407716949 |
| Regulation of Stem Cell Differentiation (GO:2000736) | 1.404125478 |
| Regulation of Transcription Elongation by RNA Polymerase II (GO:0034243) | 1.384185821 |
| Spliceosome Conformational Change to Release U4 (Or U4atac) and U1 (Or U11) (GO:0000388) | 1.380798517 |
| MRNA Transport (GO:0051028) | 1.368475487 |
| Positive Regulation of Ras Protein Signal Transduction (GO:0046579) | 1.368367 |
| Protein Neddylation (GO:0045116) | 1.368367 |
| Box C/D snoRNP Assembly (GO:0000492) | 1.368004067 |
| Regulation of Pinocytosis (GO:0048548) | 1.368004067 |
| Formation of Translation Preinitiation Complex (GO:0001731) | 1.368004067 |
| Response to Leptin (GO:0044321) | 1.368004067 |
| Synaptic Vesicle Fusion to Presynaptic Active Zone Membrane (GO:0031629) | 1.368004067 |
| Positive Regulation of Intracellular Estrogen Receptor Signaling Pathway (GO:0033148) | 1.368004067 |
| Positive Regulation of Smooth Muscle Cell Differentiation (GO:0051152) | 1.368004067 |
| Protein-DNA Complex Organization (GO:0071824) | 1.368004067 |
| Regulation of DNA Metabolic Process (GO:0051052) | 1.365054335 |
| Ribonucleoprotein Complex Biogenesis (GO:0022613) | 1.365054335 |
| Nuclear-transcribed mRNA Catabolic Process (GO:0000956) | 1.347704341 |
| Regulation of DNA-templated Transcription (GO:0006355) | 1.34554304 |
| Spliceosomal Complex Assembly (GO:0000245) | 1.342412076 |
| Regulation of Mitotic Metaphase/Anaphase Transition (GO:0030071) | 1.341651057 |
| Positive Regulation of Transcription by RNA Polymerase II (GO:0045944) | 1.317930369 |
| Regulation of Oxidative Stress-Induced Intrinsic Apoptotic Signaling Pathway (GO:1902175) | 1.31665315 |
| Cytoplasmic Translational Initiation (GO:0002183) | 1.305248292 |
| Intracellular Receptor Signaling Pathway (GO:0030522) | 1.305248292 |

K

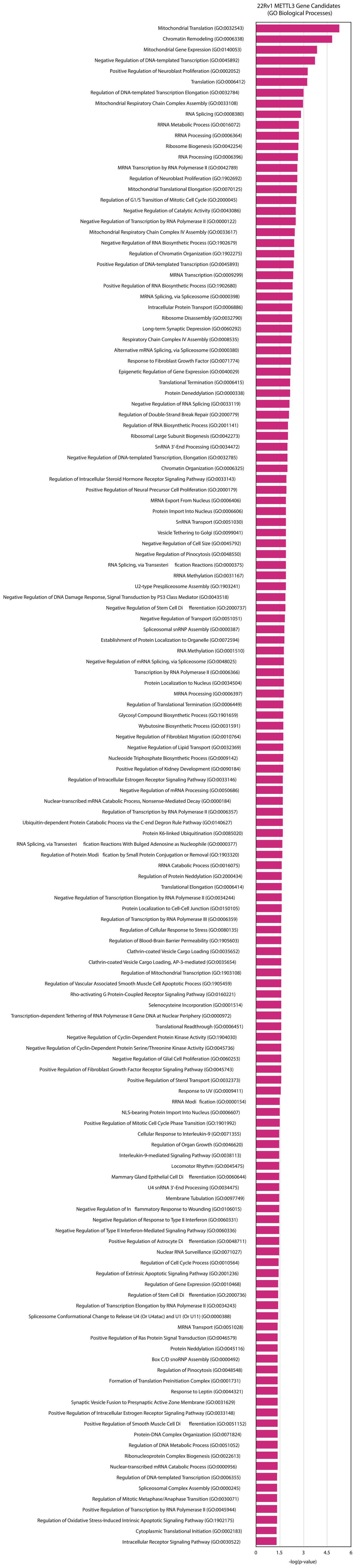

**L**

22Rv1 AR/METTL3 Shared Gene Candidates (GO Biological Processes)

| Gene Ontology (p<0.05) | -log(p-value) |
|---|---|
| Positive Regulation of Signal Transduction by P53 Class Mediator (GO:1901798) | 3.511626953 |
| Negative Regulation of Lens Fiber Cell Differentiation (GO:1902747) | 3.438460327 |
| Regulation of Lens Fiber Cell Differentiation (GO:1902746) | 3.204188068 |
| T-helper Cell Differentiation (GO:0042093) | 2.286112703 |
| TRNA Splicing, via Endonucleolytic Cleavage and Ligation (GO:0006388) | 2.248183324 |
| Spermine Metabolic Process (GO:0008215) | 2.164725124 |
| Positive Regulation of Cardiac Muscle Contraction (GO:0060452) | 2.164725124 |
| Negative Regulation of TGF-Beta Receptor Signaling Pathway (GO:0030512) | 2.127174577 |
| Alpha-beta T Cell Activation Involved in Immune Response (GO:0002287) | 2.088957728 |
| T Cell Differentiation Involved in Immune Response (GO:0002292) | 2.088957728 |
| Negative Regulation of MAPK Cascade (GO:0043409) | 2.065812077 |
| DNA Damage Response, Signal Transduction by P53 (GO:0043517) | 2.002747995 |
| TRNA Modification (GO:0006400) | 1.97610781 |
| Negative Regulation of Epithelial Cell Differentiation (GO:0030857) | 1.93219925 |
| Positive Regulation of ERAD Pathway (GO:1904294) | 1.93219925 |
| Negative Regulation of ERK1 and ERK2 Cascade (GO:0070373) | 1.930338963 |
| CD4-positive, Alpha-Beta T Cell Activation (GO:0035710) | 1.858040929 |
| Positive Regulation of Prostaglandin Biosynthetic Process (GO:0031394) | 1.858040929 |
| Positive Regulation of Striated Muscle Contraction (GO:0045989) | 1.858040929 |
| Negative Regulation of JNK Cascade (GO:0046329) | 1.85344837 |
| Regulation of Cell Development (GO:0060284) | 1.797554171 |
| Regulation of Collateral Sprouting (GO:0048670) | 1.740843069 |
| Endothelin Receptor Signaling Pathway (GO:0086100) | 1.740843069 |
| Intracellular Oxygen Homeostasis (GO:0032364) | 1.740843069 |
| Polyamine Biosynthetic Process (GO:0006596) | 1.740843069 |
| Protein Localization to Cytoplasmic Stress Granule (GO:1903608) | 1.740843069 |
| Regulation of Mitotic Nuclear Division (GO:0007088) | 1.735892232 |
| Negative Regulation of Epithelial to Mesenchymal Transition (GO:0010719) | 1.731585802 |
| Regulation of Anoikis (GO:2000209) | 1.690772571 |
| Regulation of Transforming Growth Factor Beta Receptor Signaling Pathway (GO:0017015) | 1.684739971 |
| Regulation of Prostaglandin Biosynthetic Process (GO:0031392) | 1.639421554 |
| Spermidine Metabolic Process (GO:0008216) | 1.639421554 |
| TRNA Pseudouridine Synthesis (GO:0031119) | 1.639421554 |
| White Fat Cell Differentiation (GO:0050872) | 1.639421554 |
| Phosphatidylserine Acyl-Chain Remodeling (GO:0036150) | 1.639421554 |
| Positive Regulation of Unsaturated Fatty Acid Biosynthetic Process (GO:2001280) | 1.639421554 |
| Positive Regulation of Vascular Associated Smooth Muscle Cell Migration (GO:1904754) | 1.639421554 |
| Regulation of Transferase Activity (GO:0051338) | 1.638643872 |
| Positive Regulation of Heart Contraction (GO:0045823) | 1.638643872 |
| Positive Regulation of Protein-Containing Complex Assembly (GO:0031334) | 1.632682636 |
| Negative Regulation of Translation (GO:0017148) | 1.593188319 |
| Positive Regulation of Lymphocyte Differentiation (GO:0045621) | 1.587025092 |
| Positive Regulation of Myoblast Differentiation (GO:0045663) | 1.587025092 |
| Regulation of Nervous System Development (GO:0051960) | 1.553657057 |
| Lipoxygenase Pathway (GO:0019372) | 1.55021649 |
| Regulation of ERAD Pathway (GO:1904292) | 1.542234278 |
| Cellular Response to Xenobiotic Stimulus (GO:0071466) | 1.542234278 |
| T-helper 1 Cell Differentiation (GO:0045063) | 1.542234278 |
| Regulation of Neurogenesis (GO:0050767) | 1.515832207 |
| Blood Vessel Endothelial Cell Migration (GO:0043534) | 1.497519189 |
| Vasoconstriction (GO:0042310) | 1.470753178 |
| Negative Regulation of Smooth Muscle Cell Apoptotic Process (GO:0034392) | 1.470753178 |
| Positive Regulation of Cell Morphogenesis (GO:0010770) | 1.470753178 |
| Positive Regulation of T Cell Differentiation (GO:0045582) | 1.46660621 |
| Positive Regulation of Cellular Component Biogenesis (GO:0044089) | 1.462789833 |
| GPI Anchor Biosynthetic Process (GO:0006506) | 1.454884067 |
| GPI Anchor Metabolic Process (GO:0006505) | 1.454884067 |
| Stress Granule Assembly (GO:0034063) | 1.454884067 |
| Regulation of Cell Cycle G1/S Phase Transition (GO:1902806) | 1.422982797 |
| Regulation of Cardiac Muscle Hypertrophy (GO:0010611) | 1.414170367 |
| T-helper 1 Type Immune Response (GO:0042088) | 1.414170367 |
| Cellular Response to Tumor Necrosis Factor (GO:0071356) | 1.403561437 |
| Negative Regulation of Cold-Induced Thermogenesis (GO:0120163) | 1.400939125 |
| RNA Splicing, via Endonucleolytic Cleavage and Ligation (GO:0000394) | 1.399240638 |
| Endocytosis Involved in Viral Entry Into Host Cell (GO:0075509) | 1.399240638 |
| Protein Localization to Condensed Chromosome (GO:1903083) | 1.399240638 |
| DNA-templated Transcription Initiation (GO:0006352) | 1.38445184 |
| Regulation of Nucleotide-Excision Repair (GO:2000819) | 1.375236903 |
| Ventricular Septum Morphogenesis (GO:0060412) | 1.375236903 |
| Protein Quality Control for Misfolded or Incompletely Synthesized Proteins (GO:0006515) | 1.375236903 |
| Negative Regulation of Mitotic Cell Cycle (GO:0045930) | 1.337957406 |
| Positive Regulation of Leukocyte Migration (GO:0002687) | 1.337957406 |
| V(D)J Recombination (GO:0033151) | 1.33434045 |
| Mitochondrial tRNA Modification (GO:0070900) | 1.33434045 |
| Protein Localization to Kinetochore (GO:0034501) | 1.33434045 |
| Regulation of JNK Cascade (GO:0046328) | 1.329448317 |
| Regulation of T Cell Differentiation (GO:0045580) | 1.319168056 |
| Negative Regulation of Multicellular Organismal Process (GO:0051241) | 1.315900585 |
| GPI Anchored Protein Biosynthesis (GO:0180046) | 1.302218487 |
| Positive Regulation of Response to Endoplasmic Reticulum Stress (GO:1905898) | 1.302218487 |

**M**

22Rv1 AR/METTL3 Shared Gene Candidates (GO Biological Processes)

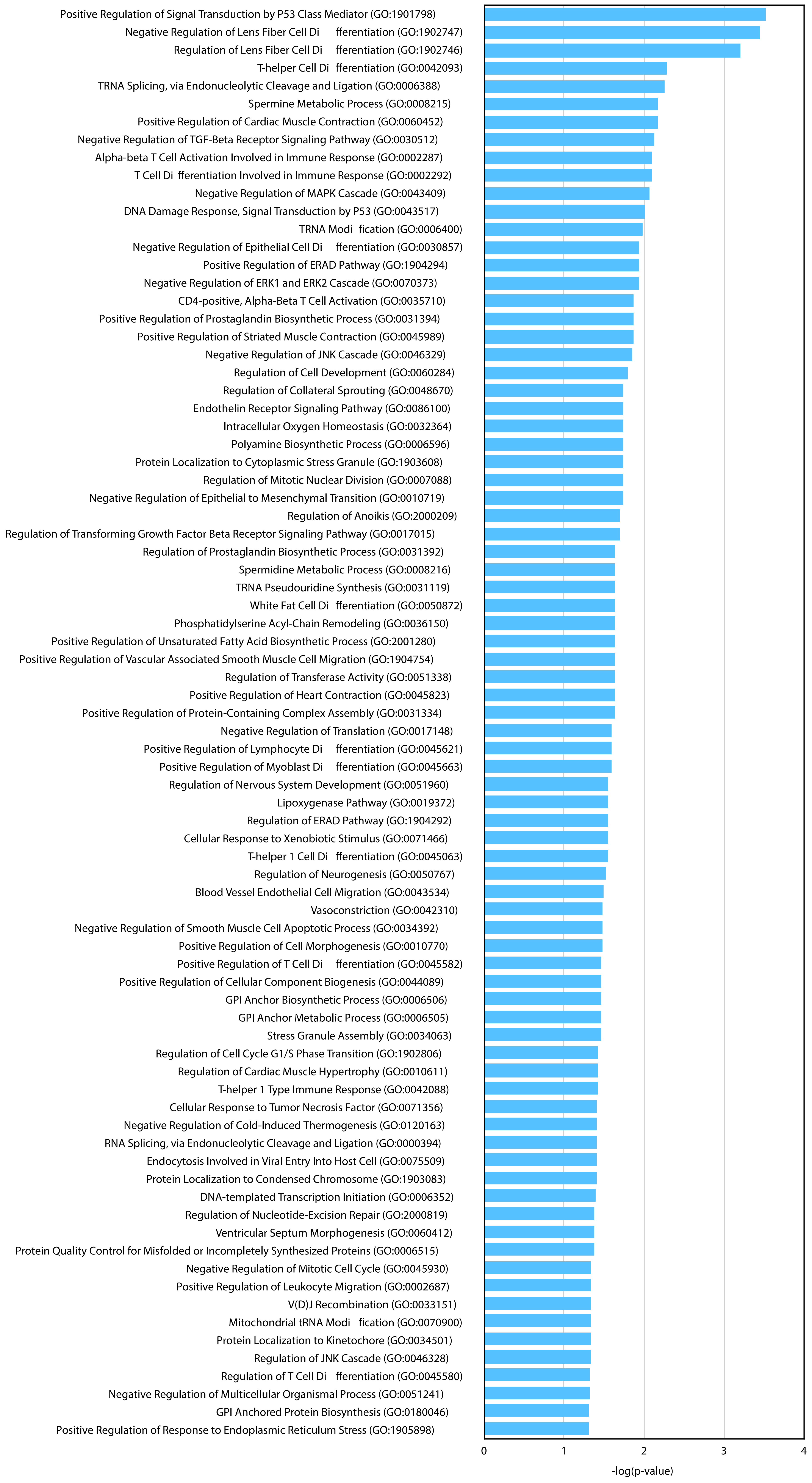


**N**

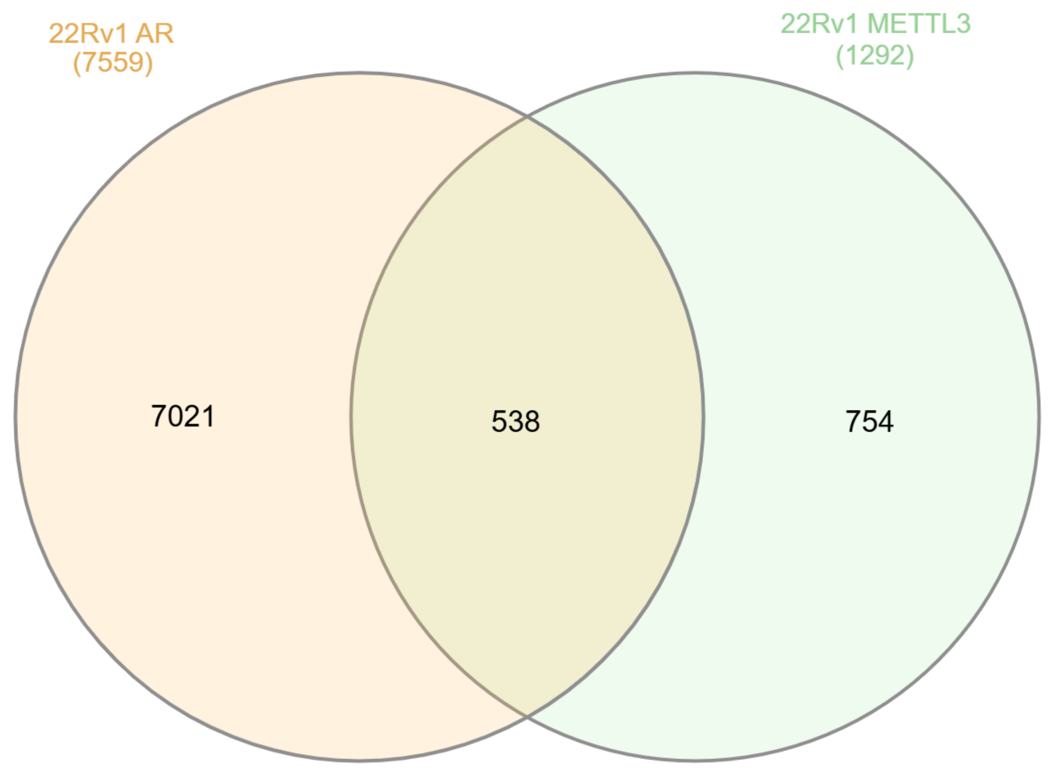


22Rv1 ChIP-seq Gene Candidates (adjRPscore >0.01)

**O**

AR/METTL3 Shared Gene Candidates (LNCaP and 22Rv1) GO Biological Processes

| Gene Ontology (p<0.05) | -log(p-value) |
|---|---|
| Alpha-beta T Cell Activation Involved in Immune Response (GO:0002287) | 2.7907 |
| T Cell Differentiation Involved in Immune Response (GO:0002292) | 2.7907 |
| Protein Localization to Condensed Chromosome (GO:1903083) | 2.4213 |
| T-helper Cell Differentiation (GO:0042093) | 2.3689 |
| Protein Localization to Kinetochore (GO:0034501) | 2.3510 |
| Astral Microtubule Organization (GO:0030953) | 2.2264 |
| Positive Regulation of ERAD Pathway (GO:1904294) | 2.0695 |
| Protein Localization to Chromosome, Centromeric Region (GO:0071459) | 2.0234 |
| Positive Regulation of ATP-dependent Activity (GO:0032781) | 2.0234 |
| Regulation of ERAD Pathway (GO:1904292) | 1.7925 |
| T-helper 1 Cell Differentiation (GO:0045063) | 1.7925 |
| SnRNA Metabolic Process (GO:0016073) | 1.7287 |
| T-helper 1 Type Immune Response (GO:0042088) | 1.6988 |
| Positive Regulation of Response to Endoplasmic Reticulum Stress (GO:1905898) | 1.6154 |
| Negative Regulation of Transforming Growth Factor Beta Receptor Signaling Pathway (GO:0030512) | 1.5526 |
| Mitochondrial Respiratory Chain Complex I Assembly (GO:0032981) | 1.4306 |
| NADH Dehydrogenase Complex Assembly (GO:0010257) | 1.4306 |
| Positive Regulation of Anoikis (GO:2000210) | 1.4157 |
| Positive Regulation of Dendritic Cell Differentiation (GO:2001200) | 1.4157 |
| Positive Regulation of Dopamine Receptor Signaling Pathway (GO:0060161) | 1.4157 |
| Negative Regulation of Translation in Response to Stress (GO:0032055) | 1.4157 |
| T-helper 2 Cell Differentiation (GO:0045064) | 1.4157 |
| Negative Regulation of CD4-positive, Alpha-Beta T Cell Differentiation (GO:0043371) | 1.4157 |
| Cellular Response to Heat (GO:0034605) | 1.4105 |
| Mitochondrial Respiratory Chain Complex Assembly (GO:0033108) | 1.3944 |
| Organelle Transport Along Microtubule (GO:0072384) | 1.3909 |
| Protein Monoubiquitination (GO:0006513) | 1.3909 |
| Mitochondrial Electron Transport, NADH to Ubiquinone (GO:0006120) | 1.3719 |
| Hemopoiesis (GO:0030097) | 1.3403 |
| Extracellular Exosome Biogenesis (GO:0097734) | 1.3382 |
| Regulation of RNA Polymerase II Transcription Preinitiation Complex Assembly (GO:0045898) | 1.3382 |
| Heat Acclimation (GO:0010286) | 1.3382 |
| Negative Regulation of Nuclear Receptor-Mediated Glucocorticoid Signaling Pathway (GO:2000323) | 1.3382 |
| Negative Regulation of Peroxisome Proliferator Activated Receptor Signaling Pathway (GO:0035359) | 1.3382 |
| Cellular Heat Acclimation (GO:0070370) | 1.3382 |
| TRNA Decay (GO:0016078) | 1.3382 |
| Dendrite Extension (GO:0097484) | 1.3382 |
| Mitochondrial Outer Membrane Permeabilization (GO:0097345) | 1.3382 |

**P**

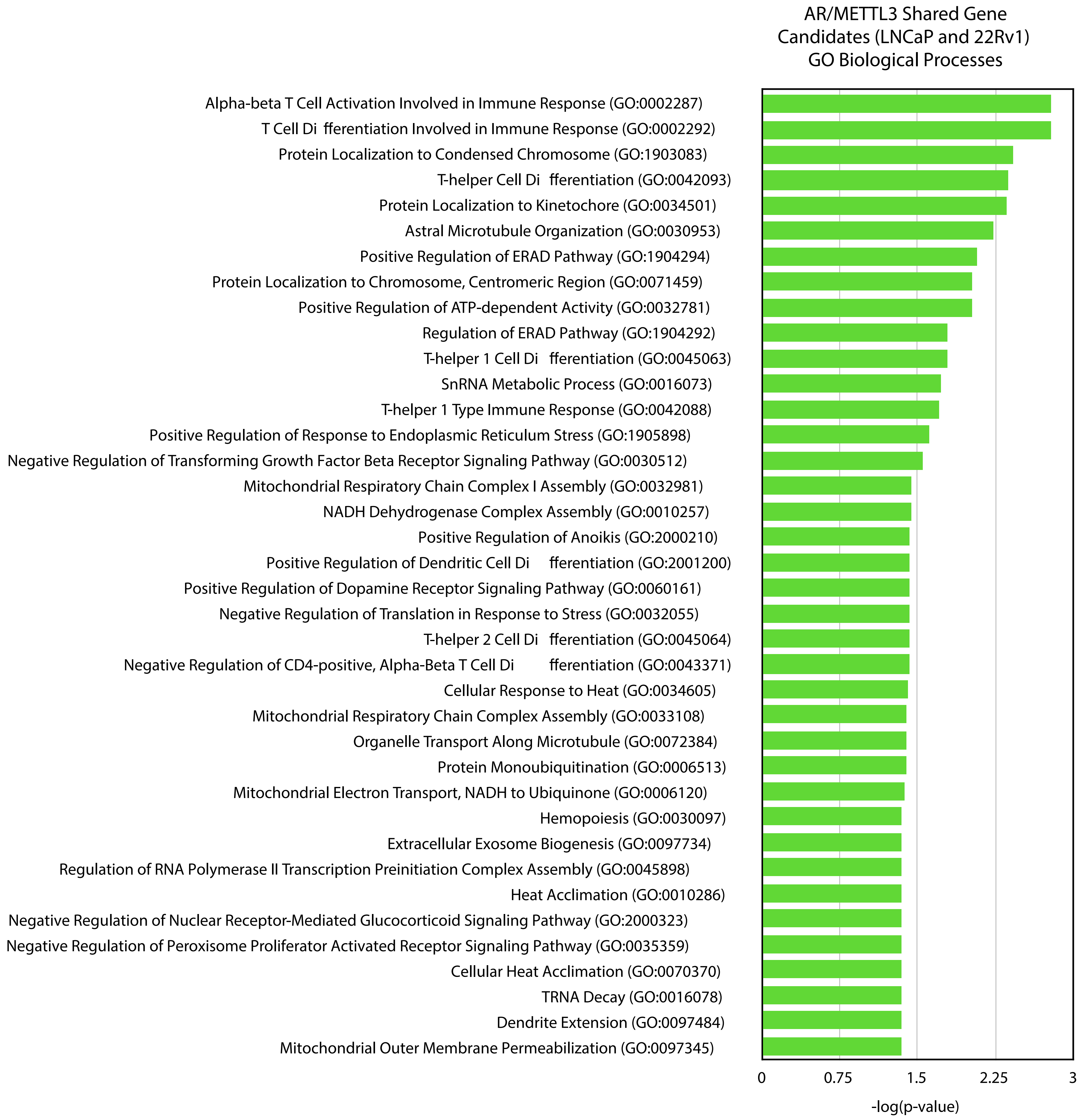


**Q**

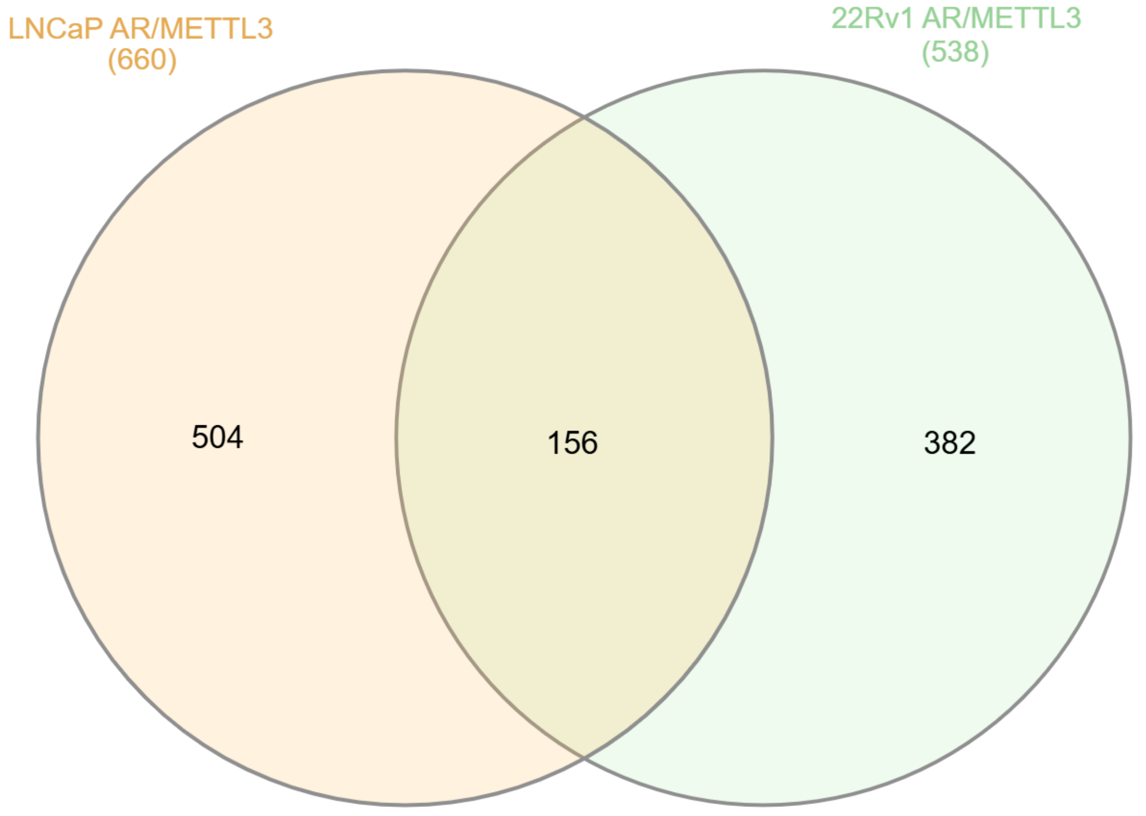


Nested LNCaP/22Rv1 AR/METTL3 Gene Candidates (adjRPscore >0.01)

**R**

AR/METTL3 Shared Gene Candidates
(LNCaP-shMETTL3-1 and LNCaP-shMETTL3-2) GO Biological Processes

| Gene Ontology (p<0.05) | -log(p-value) |
|---|---|
| Negative Regulation of Cell Division (GO:0051782) | 3.104156092 |
| Regulation of Phagocytosis (GO:0050764) | 2.733861163 |
| Regulation of Cell Division (GO:0051302) | 2.378829706 |
| Defense Response to Gram-positive Bacterium (GO:0050830) | 2.168190646 |
| Negative Regulation of Cell Cycle Process (GO:0010948) | 2.07219371 |
| Response to Glucose (GO:0009749) | 1.782143487 |
| Negative Regulation of G0 to G1 Transition (GO:0070317) | 1.718873582 |
| Neg Reg Intrinsic Apoptotic Signaling Pathway in Response to Hydrogen Peroxide (GO:1903751) | 1.718873582 |
| Ribonucleoside Monophosphate Biosynthetic Process (GO:0009156) | 1.718873582 |
| Eosinophil Activation Involved in Immune Response (GO:0002278) | 1.718873582 |
| Eosinophil Degranulation (GO:0043308) | 1.718873582 |
| Eosinophil Mediated Immunity (GO:0002447) | 1.718873582 |
| Antibody-dependent Cellular Cytotoxicity (GO:0001788) | 1.64051606 |
| Positive Regulation of Cholesterol Storage (GO:0010886) | 1.64051606 |
| Positive Regulation of Chromosome Separation (GO:1905820) | 1.64051606 |
| Negative Regulation of Cytokinesis (GO:0032466) | 1.64051606 |
| Negative Regulation of Hydrogen Peroxide-Mediated Programmed Cell Death (GO:1901299) | 1.64051606 |
| Heat Acclimation (GO:0010286) | 1.64051606 |
| Negative Regulation of Nuclear Receptor-Mediated Glucocorticoid Signaling Pathway (GO:2000323) | 1.64051606 |
| Positive Regulation of Mitotic Sister Chromatid Segregation (GO:0062033) | 1.64051606 |
| Cellular Heat Acclimation (GO:0070370) | 1.64051606 |
| Cholesterol Import (GO:0070508) | 1.64051606 |
| Neutrophil Degranulation (GO:0043312) | 1.64051606 |
| Type IIa Hypersensitivity (GO:0001794) | 1.64051606 |
| Natural Killer Cell Degranulation (GO:0043320) | 1.64051606 |
| Positive Regulation of Golgi to Plasma Membrane Protein Transport (GO:0042998) | 1.64051606 |
| De Novo' AMP Biosynthetic Process (GO:0044208) | 1.574392551 |
| Positive Regulation of Gastrulation (GO:2000543) | 1.574392551 |
| Regulation of Delayed Rectifier Potassium Channel Activity (GO:1902259) | 1.574392551 |
| Regulation of Histamine Secretion by Mast Cell (GO:1903593) | 1.574392551 |
| Regulation of Nuclear Receptor-Mediated Glucocorticoid Signaling Pathway (GO:2000322) | 1.574392551 |
| Positive Regulation of Attachment of Mitotic Spindle Microtubules to Kinetochore (GO:1902425) | 1.517223425 |
| GMP Biosynthetic Process (GO:0006177) | 1.517223425 |
| IMP Biosynthetic Process (GO:0006188) | 1.517223425 |
| Cleavage Furrow Formation (GO:0036089) | 1.517223425 |
| Dense Core Granule Cytoskeletal Transport (GO:0099519) | 1.517223425 |
| Detection of Molecule of Bacterial Origin (GO:0032490) | 1.517223425 |
| Negative Regulation of B Cell Apoptotic Process (GO:0002903) | 1.517223425 |
| Positive Regulation of Protein Transport (GO:0051222) | 1.470036642 |
| AMP Biosynthetic Process (GO:0006167) | 1.466893254 |
| Regulation of Gastrulation (GO:0010470) | 1.466893254 |
| Regulation of Lysosome Size (GO:0062196) | 1.466893254 |
| Regulation of B Cell Apoptotic Process (GO:0002902) | 1.421957639 |
| Regulation of Attachment of Mitotic Spindle Microtubules to Kinetochore (GO:1902423) | 1.421957639 |
| Negative Regulation of Lymphocyte Apoptotic Process (GO:0070229) | 1.421957639 |
| Regulation of Endoplasmic Reticulum Unfolded Protein Response (GO:1900101) | 1.421957639 |
| Regulation of Endosome Size (GO:0051036) | 1.421957639 |
| Negative Regulation of Protein Localization to Chromatin (GO:0120186) | 1.421957639 |
| Regulation of Hair Cycle (GO:0042634) | 1.421957639 |
| Cellular Response to Muramyl Dipeptide (GO:0071225) | 1.421957639 |
| Negative Regulation of Canonical NF-kappaB Signal Transduction (GO:0043124) | 1.418932682 |
| Positive Regulation of Attachment of Spindle Microtubules to Kinetochore (GO:0051987) | 1.381386348 |
| GMP Metabolic Process (GO:0046037) | 1.381386348 |
| Positive Regulation of Chromosome Segregation (GO:0051984) | 1.381386348 |
| SNARE Complex Assembly (GO:0035493) | 1.381386348 |
| Cellular Response to Oxidised Low-Density Lipoprotein Particle Stimulus (GO:0140052) | 1.381386348 |
| Negative Regulation of Vascular Endothelial Growth Factor Receptor Signaling Pathway (GO:0030948) | 1.381386348 |
| Neutrophil Activation Involved in Immune Response (GO:0002283) | 1.381386348 |
| Regulation of Mitotic Sister Chromatid Segregation (GO:0033047) | 1.381386348 |
| Mitotic Spindle Elongation (GO:0000022) | 1.381386348 |
| Mitotic Spindle Midzone Assembly (GO:0051256) | 1.381386348 |
| Negative Regulation of Sprouting Angiogenesis (GO:1903671) | 1.344418697 |
| Positive Regulation of Protein Depolymerization (GO:1901881) | 1.344418697 |
| Positive Regulation of Triglyceride Biosynthetic Process (GO:0010867) | 1.344418697 |
| Protein Localization to Condensed Chromosome (GO:1903083) | 1.344418697 |
| Purine Ribonucleoside Monophosphate Biosynthetic Process (GO:0009168) | 1.344418697 |
| Regulation of Golgi to Plasma Membrane Protein Transport (GO:0042996) | 1.310477015 |
| Positive Regulation of Mitotic Cytokinesis (GO:1903490) | 1.310477015 |
| Regulation of Mitotic Sister Chromatid Separation (GO:0010965) | 1.310477015 |
| Modification-dependent Macromolecule Catabolic Process (GO:0043632) | 1.310477015 |
| Protein Localization to Kinetochore (GO:0034501) | 1.310477015 |
| Regulation of Protein Localization to Chromatin (GO:1905634) | 1.310477015 |
| Purine Ribonucleoside Monophosphate Metabolic Process (GO:0009167) | 1.310477015 |
| Positive Regulation of DNA Metabolic Process (GO:0051054) | 1.301174188 |

**S**

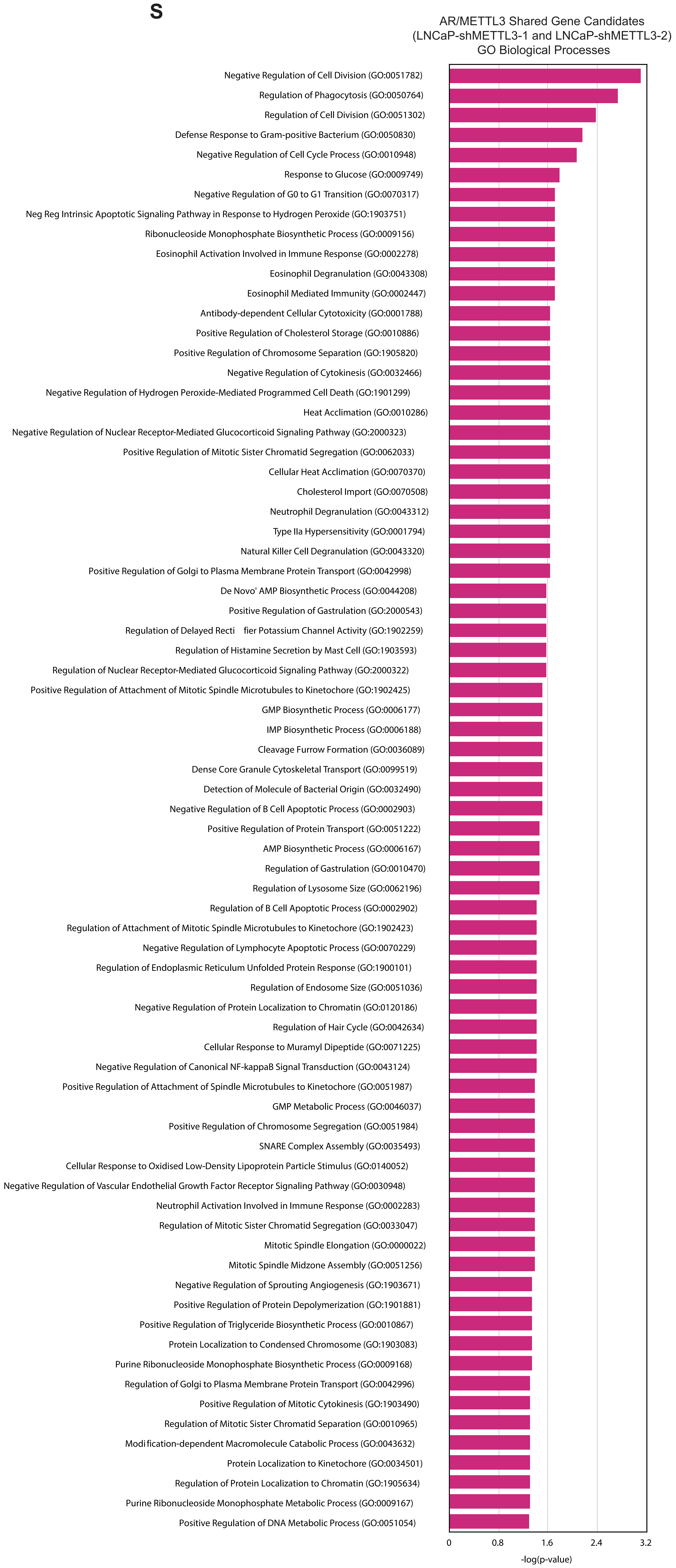
